# A Remarkable Adaptive Paradigm Of Heart Performance And Protection Emerges In Response To The Constitutive Challenge Of Marked Cardiac-Specific Overexpression Of Adenylyl Cyclase Type 8

**DOI:** 10.1101/2022.05.20.491883

**Authors:** Kirill V. Tarasov, Khalid Chakir, Daniel R. Riordon, Alexey E. Lyashkov, Ismayil Ahmet, Maria Grazia Perino, Allwin Jennifa Silvester, Jing Zhang, Mingyi Wang, Yevgeniya O. Lukyanenko, Jia-Hua Qu, Miguel Calvo-Rubio Barrera, Magdalena Juhaszova, Yelena S Tarasova, Bruce Ziman, Richard Telljohann, Vikas Kumar, Mark Ranek, John Lammons, Rostislav Beshkov, Rafael deCabo, Seungho Jun, Gizem Keceli, Ashish Gupta, Dongmei Yang, Miguel A. Aon, Luigi Adamo, Christopher H. Morrell, Walter Otu, Cameron Carroll, Shane Chambers, Nazareno Paolocci, Thanh Huynh, Karel Pacak, Robert G Weiss, Loren Field, Steven J. Sollott, Edward G Lakatta

## Abstract

Adult mice with cardiac-specific overexpression of adenylyl cyclase (AC) type VIII (TG^AC8^) adapt to an incessantly increased cAMP-induced cardiac workload (∼30% increases in heart rate, ejection fraction and cardiac output) for up to a year without signs of heart failure or excessive mortality. Here we show that despite markedly increased cardiac work, classical cardiac hypertrophy markers were absent in TG^AC8^, total left ventricular (LV) mass was not increased: a reduced LV cavity volume in TG^AC8^ was encased by thicker LV walls harboring an increased number of small cardiac myocytes and a network of small interstitial non-cardiac myocytes, manifesting increased proliferation markers and compared to WT. Protein synthesis, proteosome activity, autophagy, and Nrf-2, Hsp90α, ACC2 protein levels were increased in TG^AC8^, but LV ATP and phosphocreatine levels in vivo did not differ by genotype. 2,323 transcripts and 2,184 proteins identified in unbiased omics analyses, spanning a wide array of biological processes and molecular functions in numerous cellular compartments differed in TG^AC8^ vs WT; and over 250 canonical signaling pathways characteristic of adaptive survival circuitry of cancers, including PI3K and growth factor signaling, cytokine and T cell receptor signaling, immune responses, ROS scavenging, proliferation, protection from apoptosis, and nutrient sensing, were activated in TG^AC8^; and compared to WT there was a shift from fatty acid oxidation to increased aerobic glycolysis in the context of increased utilization of the pentose phosphate shunt and nucleotide synthesis. Thus, the adaptive paradigm, that becomes activated in the LV of TG^AC8^ in response to severe chronic, intense AC/PKA/Ca^2+^ signaling embodies many hallmarks of cancer.

## Introduction

Adaptations have evolved in all organisms to cope with both acute and chronic internal and environmental stress. For example, even during acute exercise increased autonomic sympathetic-mediated AC/cAMP/PKA/Ca^2+^ signaling, the quintessential mediator of both acute and chronic stress, also activates acute physiologic adaptations to moderate the exercise-induced increase in sympathetic signaling. In response to repeated bouts of acute exercise induced AC/cAMP/PKA/Ca^2+^ stress chronic adaptations emerge (endurance exercise conditioning).^1–3^

Prolonged and intense chronic cAMP-mediated stress in experimental animal models results in cardiomyopathy and death.^4, 5^ During chronic pathophysiologic states e.g. chronic heart failure (CHF) in humans, AC/cAMP/PKA/Ca^2+^ signaling progressively increases as the degree of heart failure progresses, leading to cardiac inflammation, mediated in part, by cyclic-AMP-induced up-regulation of renin-angiotensin system (RAS) signaling. Standard therapies for CHF include β-adrenoreceptor blockers and RAS inhibitors,^6, 7^ which although effective, are suboptimal in amelioration of heart failure progression.

One strategy to devise novel and better therapies for heart failure, would be to uncover the full spectrum of consilient cardio-protective adaptations that can emerge in response to severe, chronic AC/cAMP/PKA/Ca^2+^ -induced cardiac stress. The young adult TG^AC8^ heart in which cardiac-specific over expression in mice of Adenylyl Cyclase (AC) Type 8, driven by the α myosin heavy chain promoter, markedly enhances AC activity and cAMP signaling, may be an ideal model in which to elucidate the features of a wide-spread an adaptive paradigm that must become engaged in response to incessant chronic activation of cardiac AC/cAMP/PKA/Ca^2+^ signaling. Specifically, concurrent with chronically increased AC activity within the young adult TG^AC8^ sinoatrial node (SAN) and LV,^8, 9^ increased heart rate (HR), measured via telemetry in the awake, unrestrained state, increased by approximately 30%, persisting in the presence of dual autonomic blockade;^8^ and LV EF is also markedly increased in TG^AC8^.^10^ Thus, the cardiac phenotype of the adult TG^AC8^ mimics cardiac responses to AC/cAMP/PKA/Ca^2+^ sympathetic autonomic input during strenuous, acute exercise, but this stressful state persists incessantly. ^8^ Although this incessant high cardiac load imposes a severe chronic stress on the heart that might be expected to lead to the near term heart failure and demise,^11, 12^ adult TG^AC8^ mice maintain this remarkable, hyperdynamic cardiac phenotype up to about a one year of age,^9, 10, 13, 14^ when signs of CHF begin to develop and cardiomyopathy ensues.^10^

We hypothesized, (1) that a panoply of intrinsic adaptive mechanisms become *concurrently* engaged in order to protect the TG^AC8^ heart during the incessant, high level of cardiac work in several months, and (2) that, some of these mechanisms are those that become activated in the endurance, trained heart, including shifts in mechanisms of energy generation, enhanced protein synthesis and quality control, and increased defenses against reactive O_2_ species (ROS) and cell death.^15, 16^ We reasoned that a discovery bioinformatics approach in conjunction with a deeper phenotypic characterization of TG^AC8^ LV, would generate a number of testable hypotheses about the characteristics of some of these mechanisms utilized to sustain this adaptive paradigm of heart performance and protection in response to severe, chronic adenylyl cyclase-induced cardiac stress. To this end, we performed unbiased, RNASEQ and proteomic analyses of adult TG^AC8^ and WT LVs, and selectively validated genotypic differences in numerous transcripts and proteins. Our results delineate an emergent consilient pattern of adaptations within the TG^AC8^ heart in response to chronically increased AC/cAMP/PKA/Ca^2+^-signaling and offer numerous testable hypotheses to further define the details of mechanisms that underlie what we will show here to be a remarkable adaptive heart paradigm.

## Materials and Methods

### Mice

All studies were performed in accordance with the Guide for the Care and Use of Laboratory Animals published by the National Institutes of Health (NIH Publication no. 85-23, revised 1996). The experimental protocols were approved by the Animal Care and Use Committee of the National Institutes of Health (protocol #441-LCS-2019). A breeder pair of TG^AC8^ over expression mice, generated by ligating the murine α-myosin heavy chain promoter to a cDNA coding for human TG^AC8^, were a gift from Nicole Defer/Jacques Hanoune, Unite de Recherches, INSERM U-99, Hôpital Henri Mondor, F-94010 Créteil, France.^9^ Mice were bred adult (3 mo) and housed in a climate-controlled room with 12-hour light cycle and free access to food and water, as previously described.^8^

Detailed methodology for RNASEQ and LV Proteome analyses, WB and RT-qPCR analyses, echocardiography, heart and cardiac tissue isolation, isolated perfused working heart experiments, nuclear EdU and BrdU uptake in cells within LV tissue, adenyl cyclase activity, immunohistochemistry, protein synthesis, misfolding and degradation, autophagy, NMR spectroscopy, transmission electron microscopy, mitochondrial permeability transition threshold, are provided in the online supplement.

## Results

### Cardiac Structure and Performance

We performed echocardiography and cardiac histology for in-depth characterization of the TG^AC8^ heart structure and function. Representative echocardiograms of TG^AC8^ and WT are illustrated in **Fig.S1**, selected Echo parameters are shown in **Fig. 1**, (and a complete listing of parameters is provided in **Table S.1)**. Both EF and HR were higher in TG^AC8^ than in WT (**Fig. 1 A, B**), confirming prior reports.^8, 10^ Because stroke volume did not differ by genotype (**Fig. 1C**), cardiac output was elevated by 30% in TG^AC8^ (**Fig. 1D**) on the basis of its 30% increase in HR. Arterial blood pressure was only mildly increased in TG^AC8^ averaging 3.5 mmHg higher than in WT (**Fig. 1E**).

**Fig. 1.**
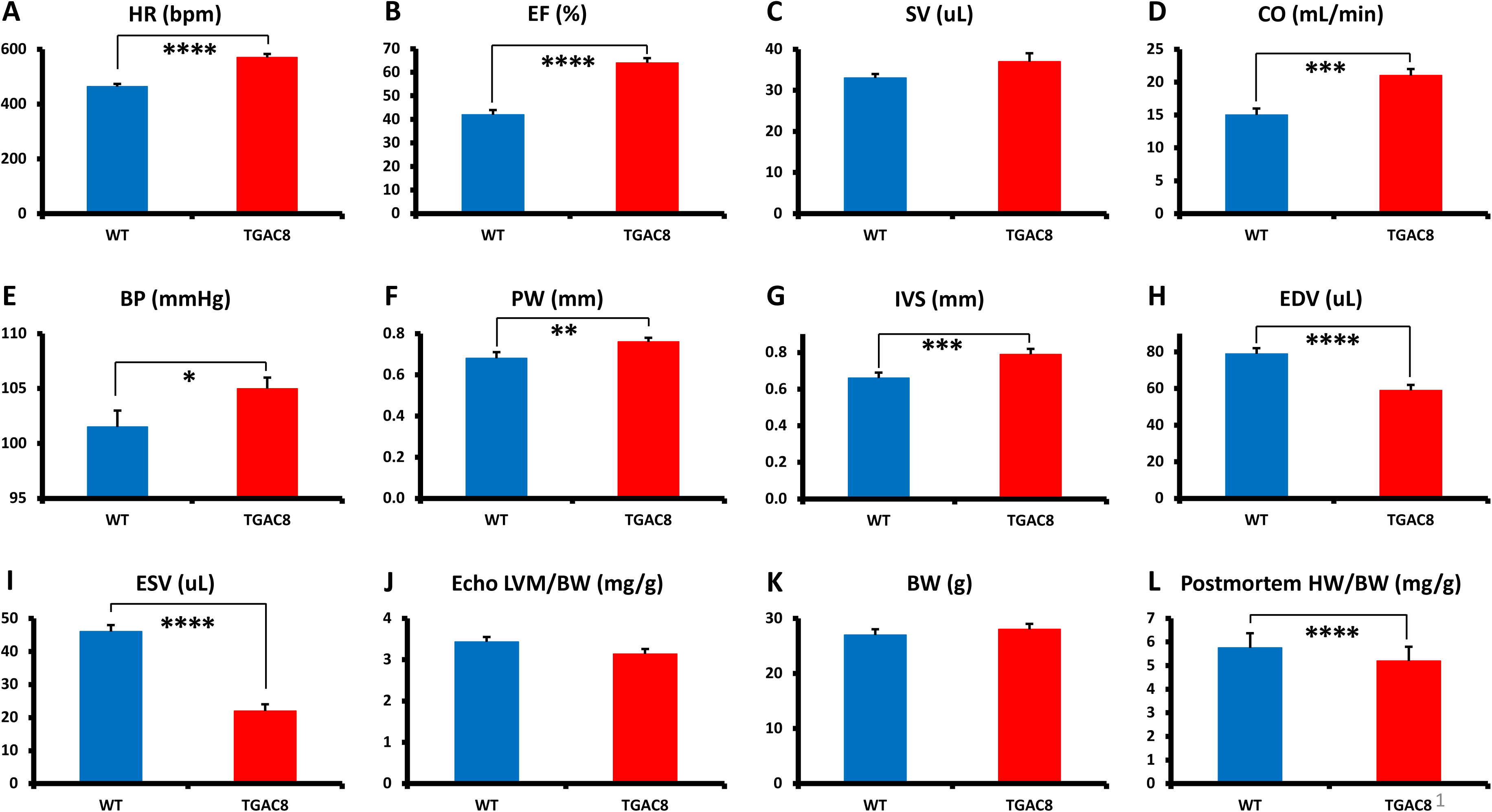
**(A-K), Echocardiographic parameters (N= 28 for TG^AC8^; WT=21); L heart weight/body weight at sacrifice (N=75 for TG^AC8^ and N=85 for WT). See Table S.1 for additional Echo parameters.**

A sustained high cardiac workload is usually expected to result in an increase of a total LV mass,^1, 17, 18^ i.e., cardiac hypertrophy. Although both the LV posterior wall and inter-ventricular septum were thicker in TG^AC8^ vs WT (indicative of an increased LV wall biomass), (**Fig. 1 F, G**) the LV cavity volume was markedly reduced (**Fig. 1 H, I**), and the echo derived **total** LV mass did not differ by genotype (**Fig. 1 J**). Postmortem measurements indicated that although the body weight did not differ between TG^AC8^ and WT (**Fig. 1 K**), the heart weight/body weight (HW/BW) was even actually, modestly reduced in TG^AC8^ vs WT (**Fig. 1 L**).

Pathological hypertrophy markers e.g., β-myosin heavy chain (MYH7), ANP (NPPA) or BNP (NPPB) were not increased in TG^AC8^ vs WT by Western Blot (WB) (**Fig. 2A**); curiously, α skeletal actin, commonly considered to be a pathologic hypertrophy marker, was increased in TG^AC8^ vs WT (**Fig. 2A**). Calcineurin (PP2B), which activates the hypertrophic response by dephosphorylating nuclear factor of activated T cells (NFAT),^19^ did not differ in WB of TG^AC8^ vs WT (**Fig. 2A)**.

**Fig. 2.**
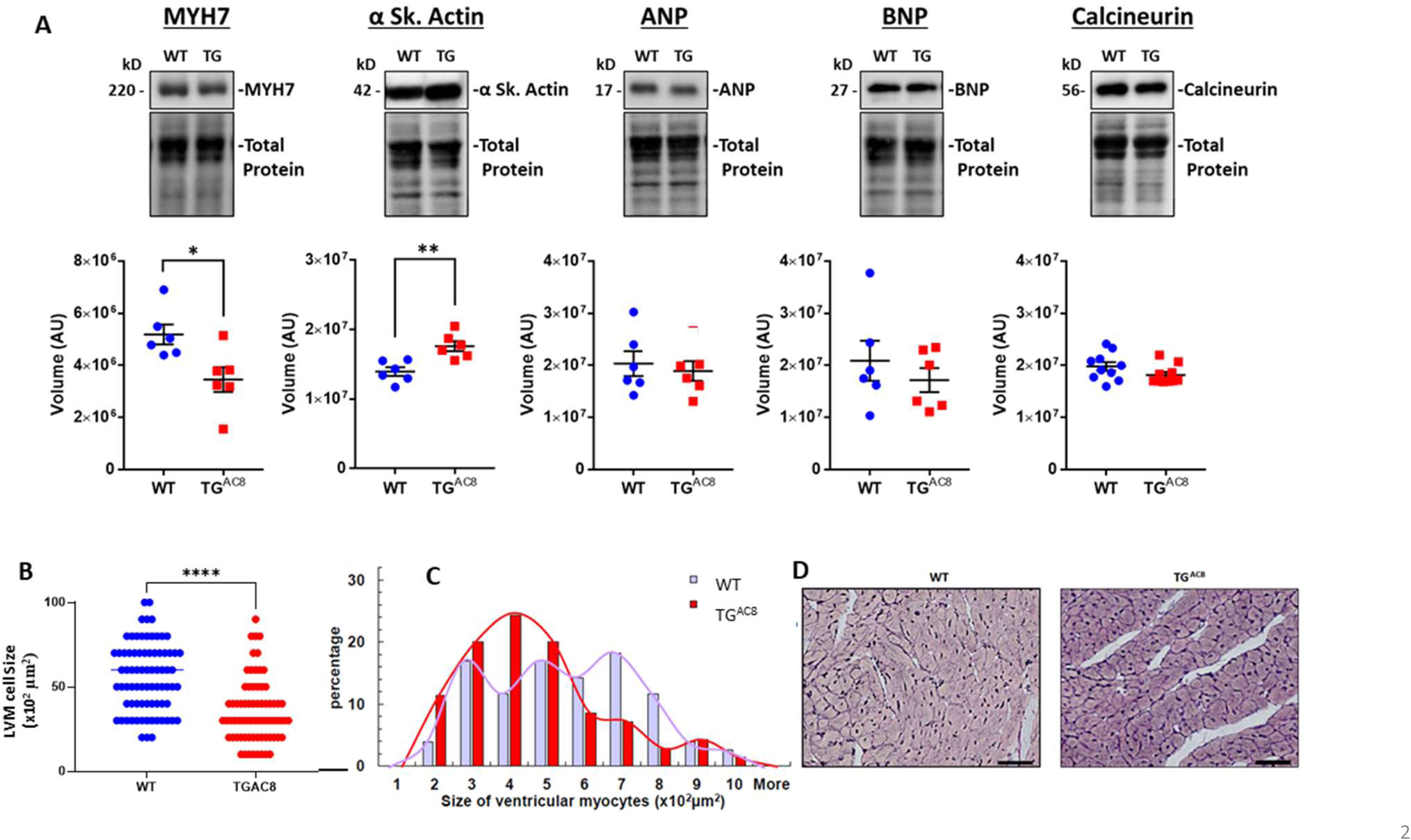
**(A) WB of Pathologic hypertrophy markers, *(B, C)* cardiac myocyte size and distributions of cardiac myocytes, (*D) Representative LV sections depicting cardiac myocyte diameters,***

### LV Histologic Analysis

The average LV cardiac myocyte size was smaller in TG^AC8^ compared to WT (**Fig. 2 B, C**), and LV myocyte size distribution was different by genotype (**Fig. 2 D**). LV collagen content in TG^AC8^ was not increased vs WT (**Fig. S. 2).**

To address the issue of DNA synthesis within LV cardiac myocyte we loaded EdU for 28-days. EdU labeled nuclei, detected in both TG^AC8^ and WT whole mount ventricular preparations were randomly scattered throughout the LV from the mitral annulus to the apex. LV myocyte nuclei, however, were rarely EdU labeled. Rather nearly all EdU labelling was detected in small interstitial cells that expressed vimentin or in cells that enclosed the capillary lumina (**Fig. 3 A-G)**, suggesting that EdU was also incorporated within the DNA of endothelial cells in both WT and TG^AC8^. However, total nuclear EdU labeling was 3-fold higher in TG^AC8^ than in WT **Fig. 3 H**.

**Fig. 3.**
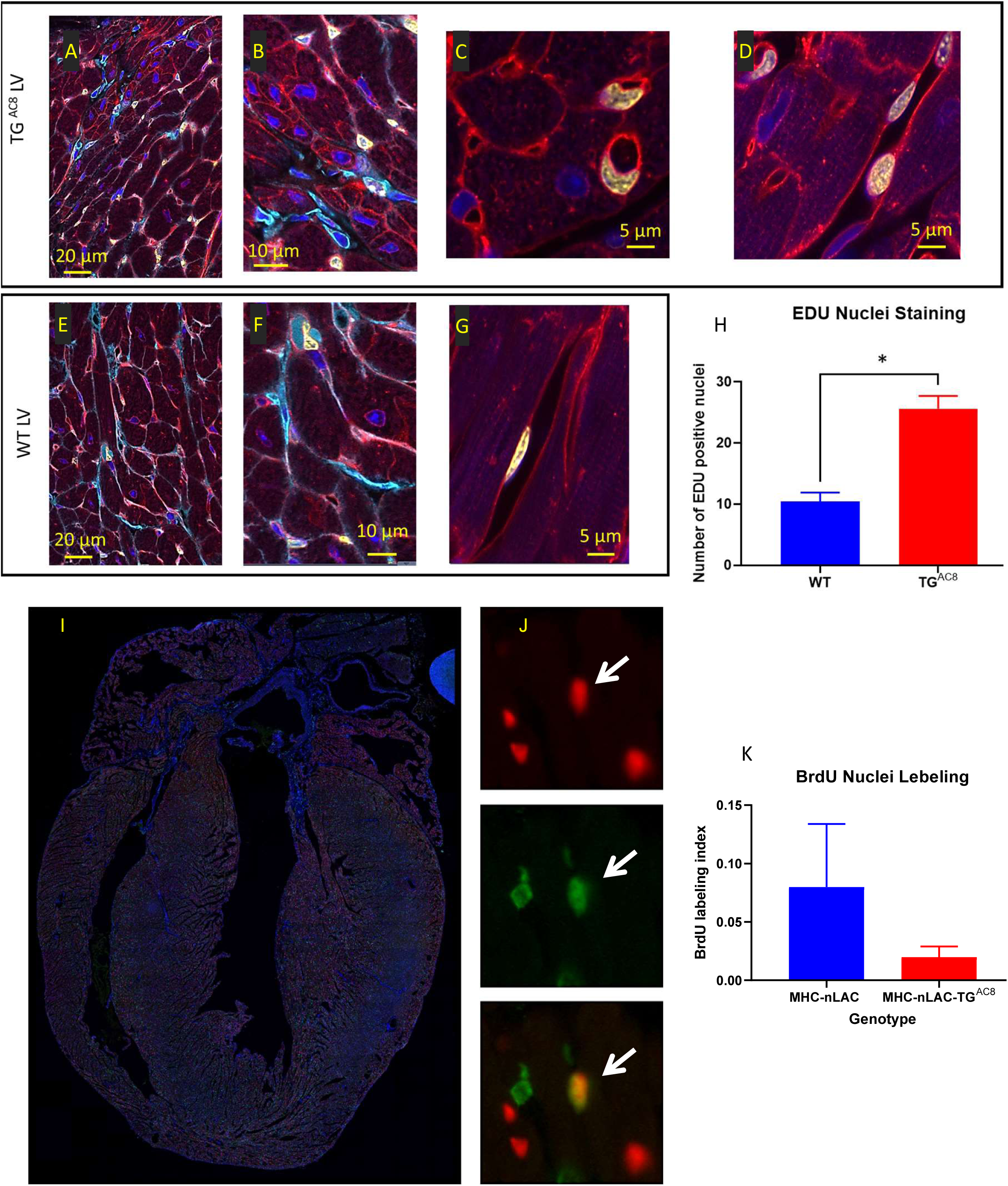
**(A-G) representative examples of confocal images (400x) of LV WGA (red), vimentin (Cyan), EdU (yellow), DAPI (blue), (H) average number of EdU labeled nuclei positive field counts in LV TG^AC8^ vs WT (N=3 mice in each genotype). Detection of cardiomyocyte S-phase activity. (I) Section from the heart of a TG^AC8^, MHC-nLAC double-transgenic mouse subjected to 12 days of BrdU treatment. The section was processed for β-galactosidase (to identify cardiomyocyte nuclei, red signal) and BrdU (to identify DNA synthesis, green signal) immune reactivity, and then counterstained with Hoechst (which stains all nuclei, blue signal). (J) Example of an S-phase cardiomyocyte nucleus detected with this assay. The upper panel shows β-galactosidase immune reactivity (red channel), the middle panel shows BrdU immune reactivity (green channel), and the lower panel shows a red and green color combined image of the same field. The arrow identifies an S-phase cardiomyocyte nucleus, as evidenced by the overlay of β-galactosidase and BrdU immune reactivity, which appears yellow in the color combined image. (H) Graph representing S-phase activity in the TG^AC8^, MHC-nLAC double-transgenic vs. the MHC-nLAC single transgenic animals (mean +/- SEM, p=0.315; 5 mice per genotype and 3 sections per mouse were analyzed).**

To monitor cardiomyocyte S-phase activity, TG^AC8^ mice were crossed with MHC-nLAC mice (which express a nuclear-localized β-galactosidase reporter under the transcriptional regulation of the mouse α-cardiac MHC promoter; these mice are useful to identify cardiomyocyte nuclei in histologic sections).^20^ The resulting TG^AC8^, MHC-nLAC double-transgenic mice and MHC-nLAC single-transgenic mice were identified and sequestered. At 28-to-30 days of age, the mice were administered BrdU via drinking water (0.5 mg/ml, changed every 2nd day) for a total of 12 days. There was no difference in the level of ventricular cardiomyocyte S-phase activity in the TG^AC8^, MHC-nLAC double-transgenic vs. the MHC-nLAC single transgenic animals (**Fig. 3 H-K**).

Thus, the adult TG^AC8^ heart has a hyper-dynamic LV with thicker walls, harboring not only an increased number of small **cardiac** myocytes, but also an increased number of small interstitial cells and endothelial cells with increased EdU labeling vs WT. The LV cavity volumes at both end-diastole and end-systole were markedly reduced, and LV EF was markedly increased in TG^AC8^ vs WT. But, neither **total** LV mass, nor collagen content were increased in TG^AC8^ vs WT and the profile of pathologic cardiac hypertrophy markers in TG^AC8^ was absent.

### AC/cAMP/PKA/Ca^2+^ signaling

Given that the transgene in our study was an AC, we next focused on expected differences in AC/cAMP/PKA/Ca^2+^ signaling in TG^AC8^ vs WT. Immunolabeling of single LV myocytes showed that AC8 expression was markedly increased (by 8-9-fold in TG^AC8^ vs WT), and AC activity (measured in membranes isolated from LV tissue) in TG^AC8^ was 50% higher than that in WT (**Fig. 4 A-C**). Expression of the PKA catalytic subunit was increased by 65.6%, and expression of the regulatory subunit was decreased by 26.7% in TG^AC8^ by WB (**Fig. 4 D, E**). PKA activity was increased by 57.8 % in TG^AC8^ vs WT (**Fig. 4 F**).

**Fig. 4.**
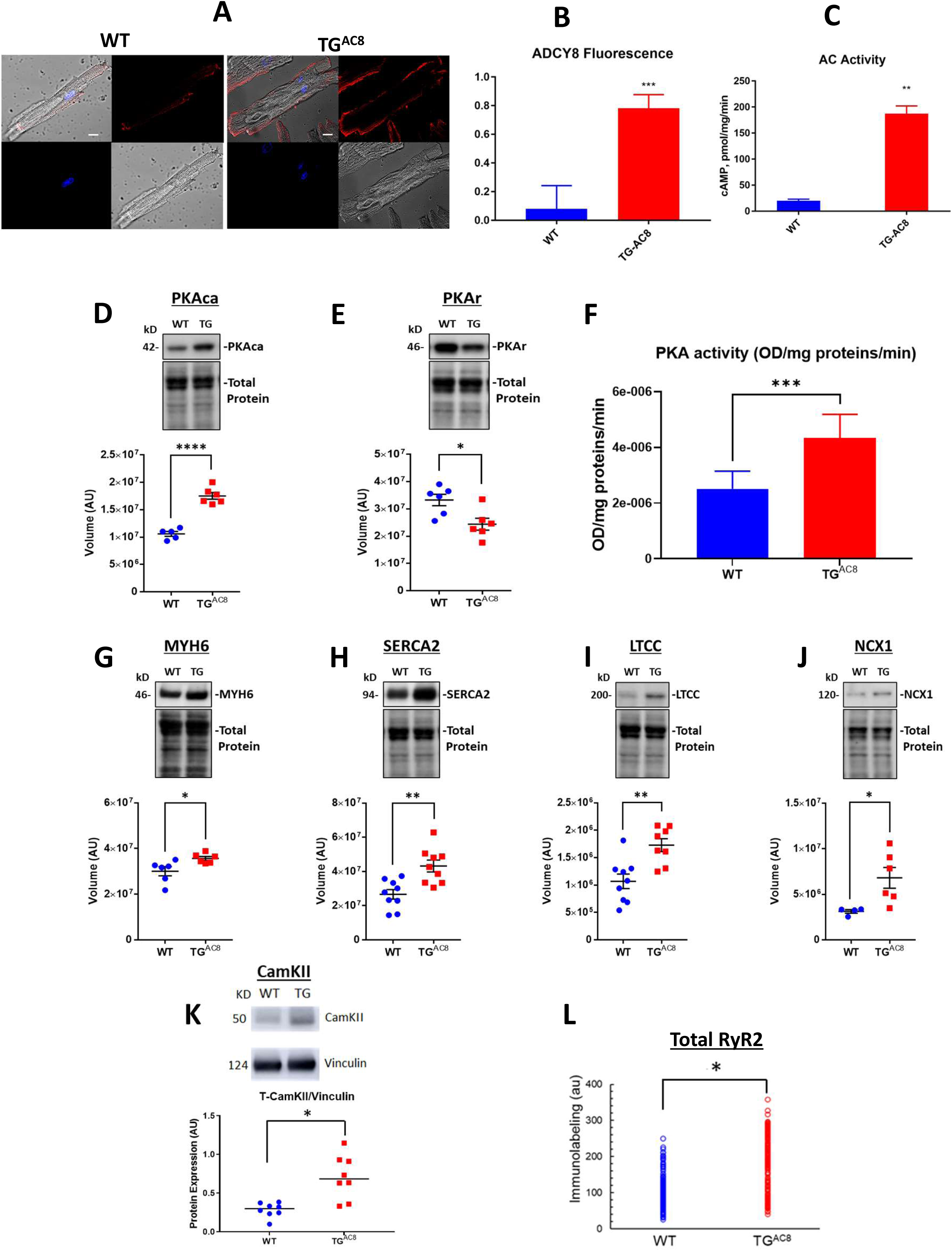
**(A)** Representative examples of ADCY8 immunolabeling in TG^AC8^ and WT LV cardiomyocytes, **(B)** Average AC8 fluorescence in LV cardiomyocytes and **(C)** Average AC activity in TG^AC8^ vs WT, **(D, E)** Expression levels of PKA catalytic and regulatory subunits, and **(F)** PKA activity in TG^AC8^ vs WT, **(G-M)** Western Blot analysis of selected proteins involved in excitation - Ca release – contraction-relaxation coupling TG^AC8^ vs WT LV. **(L)** RyR2 immunolabeling. Antibodies employed are listed in supplemental methods. (N=199 WT cells and 195 TG^AC8^ cells (each from 3 mice).

To search for mechanisms that increase contractility within cardiac myocytes we examined the expression of selected proteins downstream of PKA signaling that are involved in excitation/Ca^2+^ release/contraction/relaxation. Western Blot analysis revealed that a number of proteins that determine increase cardiac myocyte performance were upregulated in TG^AC8^ vs WT, including αMHC (MYH6 by 18.7%), SERCA2 (ATP2A2 by 62.0%), L-type Ca Channel (Cav1.2, by 61.8%), and NCX1 (AKA SLC8A1) by 117.6%, and CaMKII by 35.0% (**Fig. 4 G-K**).

Immunolabeling of total RyR2 was increased by 66% in TG^AC8^ vs WT (**Fig. 4 L**). This pattern of increased protein expression consistent with increased Ca^2+^ flux into and out of cardiac myocytes, an increased Ca^2+^ cycling between SR and cytosol, and increased myosin cross-bridge kinetics during heart contraction in TG^AC8^ vs WT.

Thus, as would be expected in the context of markedly increased transcription of AC type VIII, AC and PKA protein levels and activities, and levels of proteins downstream of PKA signaling were markedly increased and associated with a chronic, marked increase in LV performance.

### Myocardial and plasma catecholamines

It is well known, that PKA signaling activates a cascade of enzymes beginning with tyrosine hydroxylase (TH), that results in a production of catecholamines (**Fig. S.3 A**). Among these enzymes, dopamine decarboxylase, which converts DOPA to dopamine, was increased in TG^AC8^ vs WT (**Fig. S.3 B**), dopamine b-hydroxylase, which converts dopamine to norepinephrine did not differe by genotype, and phenylethanolamine N-methyltransferase (PNMT), which converts norepinephrine to epinephrine was reduced in TGAC8 vs WT. In the context of these genotypic differences in enzyme levels LV tissue dopamine was increased and DHPG was reduced and NE was borderline reduced (p=0.08) **Fig. S.3 B**.

Interestingly, as noted previosly ^8^, plasma levels of dopa and dopamine were also incresed in TG^AC8^ vs WT, whereas norepinephrine, epinephrine, DOPAC and DHPG were reduced (**Fig. S.3 C**).

### Protein Synthesis, Degradation and Quality Control

Because PKA signaling-driven increased cardiac work is known to be associated with increased protein synthesis,^21, 22^ we next compared the rate of protein synthesis in TG^AC8^ and WT LV lysates. Despite the absence of increase of total LV mass, the rate of protein synthesis was 40% higher in TG^AC8^ than in WT (**Fig. 5 A**). WB analysis indicated that expression or activation of p21Ras, p-c-Raf, MEK1/2, molecules downstream of PKA signaling that are implicated in protein synthesis, were increased in TG^AC8^ vs WT (**Fig. 5 B-C**). The transcription factor CREB1, involved in PKA signaling directed protein synthesis was increased by 58.1% in TG^AC8^ vs WT in WB analysis (**Fig. 5 D).** Expression of CITED4 (family of transcriptional coactivators that bind to several proteins, including CREB-binding protein (CBP)) was increased by 51 % in TG^AC8^ vs WT in WB analysis (**Fig. 5 E).** Expression of protein kinases, that are required for stress-induced activation of CREB1, MSK1 and MNK1, direct substrates of MAPK, were increased by 40% and 48% respectively in TG^AC8^ vs WT (**Fig. 5 F** and **G**).

**Fig. 5.**
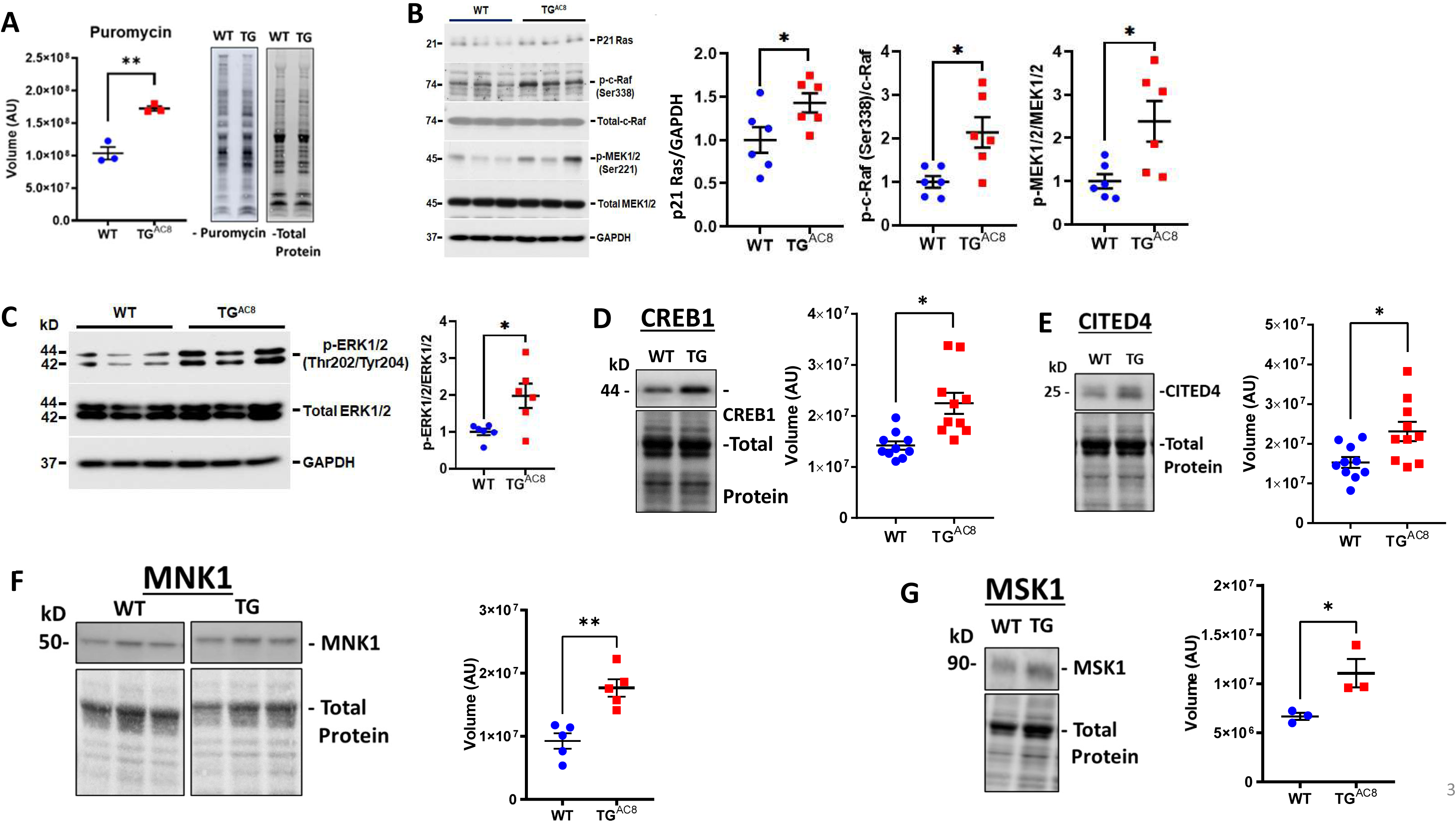
**(A)** Rate of global protein synthesis and **(B-G)** mechanisms downstream of PKA signaling involved in protein synthesis in the TG^AC8^ and WT.

### Autophagy

Proteasome activity within LV lysates was increased in TG^AC8^ vs WT **(Fig. 6 A)**. However, although there was a significant increase in the amount of soluble misfolded proteins in TG^AC8^ vs WT **(Fig. 6 B)**, insoluble protein aggregates did not accumulate within the TG^AC8^ LV (**Fig. 6 C**). This suggest that an increase in micro-autophagy of the TG^AC8^ LV circumvents the potential proteotoxic stress of aggregated protein accumulation, which can negatively impact cardiac cell health and function. A 24% increase in the expression of HSP90α (**Fig. 6 D**) in TG^AC8^ also suggested that chaperone-mediated autophagy was involved in preventing an accumulation of insoluble protein aggregates.

**Fig. 6.**
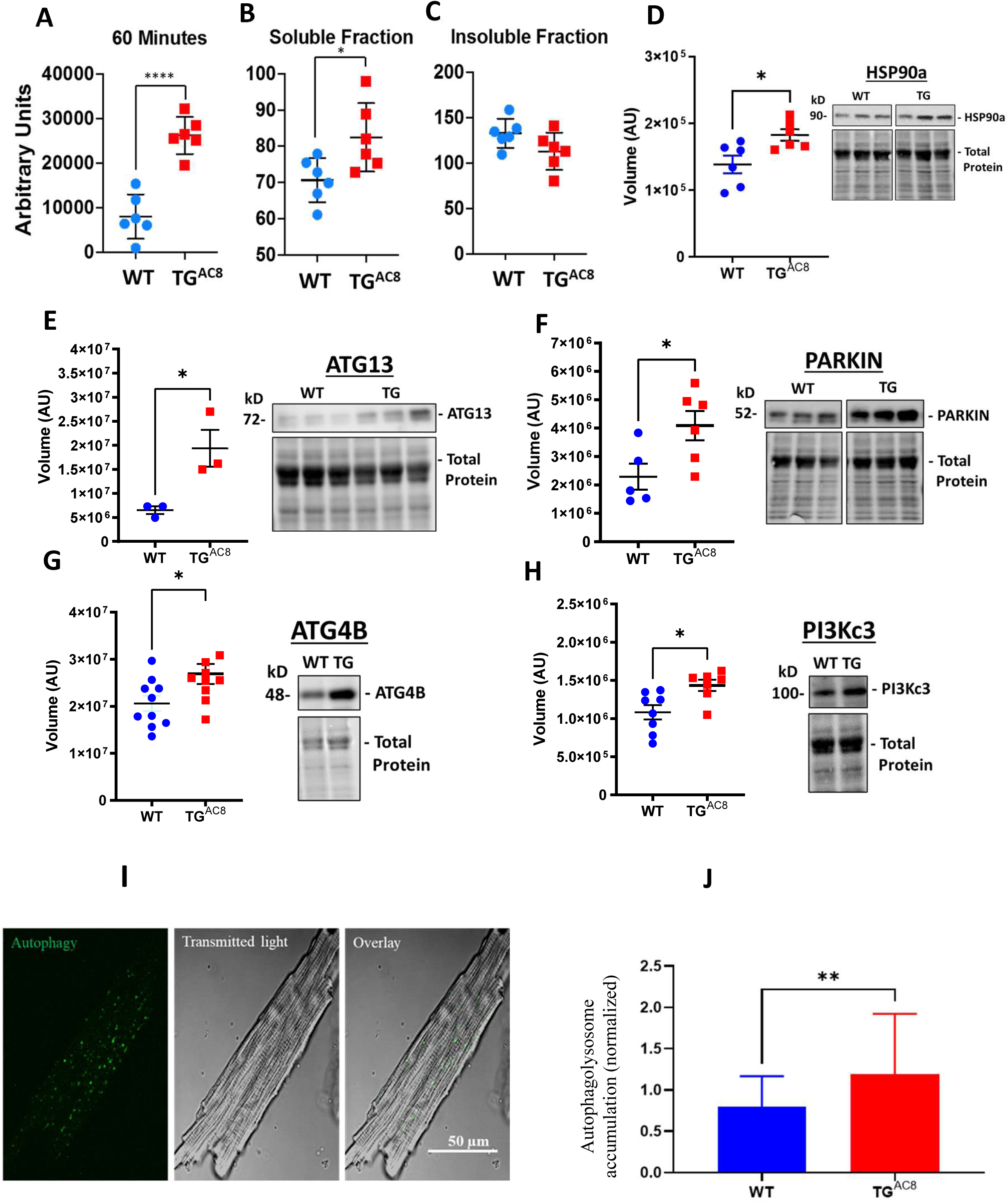
**(A)** Proteosome activity assay and **(B, C)** accumulated proteins in soluble and insoluble fractions of LV lysates in TG^AC8^ vs WT. **(D)** WB of HSP90 in TG^AC8^ and WT, **(E-H)** Expression levels of selected proteins involved in the autophagy process. **(I, J)** Autophagolysosome accumulation is enhanced in AC8 mice. The statistical significance is indicated by **(p<0.01) and t (p<0.01 in one-tailed t test).

We reasoned that another type of protein quality control, macro-autophagy, might also be activated in TG^AC8^ vs WT. Indeed, protein levels of several members of the autophagy machinery were increased in TG^AC8^ vs WT: both ATG13, a factor required for autophagosome formation and mitophagy, and the ubiquitin-protein ligase PARKIN, which protects against mitochondrial dysfunction during cellular stress, by coordinating mitochondrial quality control mechanisms in order to remove/replace dysfunctional mitochondrial components, were significantly increased in TG^AC8^ vs WT (**Fig. 6 E, F)**. Furthermore, protein levels of the cysteine protease ATG4B, involved in processing, and the lipidation/delipidation steps (conjugation/removal of phosphatidylethanolamine) and insertion of MAP1LC3, GABARAPL1, GABARAPL2 and GABARAP into membranes, during autophagy and endocytosis, were also increased (**Fig. 6 G)**. In addition, the catalytic subunit of the PI3Kc3, involved in initiation and maturation of autophagosomes and in endocytosis, was also significantly higher in the TG^AC8^ vs WT (**Fig. 6 H)**. Finally, direct measurements in single LV myocytes showed that autophagolysosomes were appreciably increased in TG^AC8^ vs WT, indicating that autophagy (mitophagy) was activated to the greater extent in the TG^AC8^ vs WT. (**Fig. 6 I, J)**.

### Mitochondrial Structure

We employed transmission electron microscopy (TEM) to directly visualize ultrastructural details of the mitochondria and cardiac myofibers in the LVs of TG^AC8^ and WT. Representative panoramic electron micrographs of LV cardiac muscle fibers and mitochondria in TG^AC8^ and WT are illustrated in **Fig. 7 A and B**. Cardiac myocytes presented a very distinctive morphology with high content of myofibrils and a large number of high-electron dense mitochondria and several capillaries surrounding cardiac myocytes are depicted (see white arrows). Cardiac myocyte ultrastructure is depicted in **Fig. 7** panels **C** and **E**, for WT mice, and in panels **D** and **F**, for TG^AC8^. Asterisks show swollen, disrupted mitochondria with lighter cristae compared to the surrounding healthy mitochondria. **Fig. 7** Panels **G-J** present quantitative stereological analyses of normal and damaged mitochondria. Although there was a mild increase in the number of damaged mitochondria (0.3 %), and in the percent of cell volume occupied by damaged mitochondria (0.4%) in TG^AC8^, the numbers of healthy mitochondria and the percent of cell volume occupied by healthy mitochondria did not differ between TG^AC8^ and WT. Nevertheless, the presence of mitochondrial deterioration at a young age is uncommon and may be further evidence for enhanced cleaning and recycling mechanisms such as autophagic signaling, (**Fig.s. 6, 7**).

**Fig. 7.**
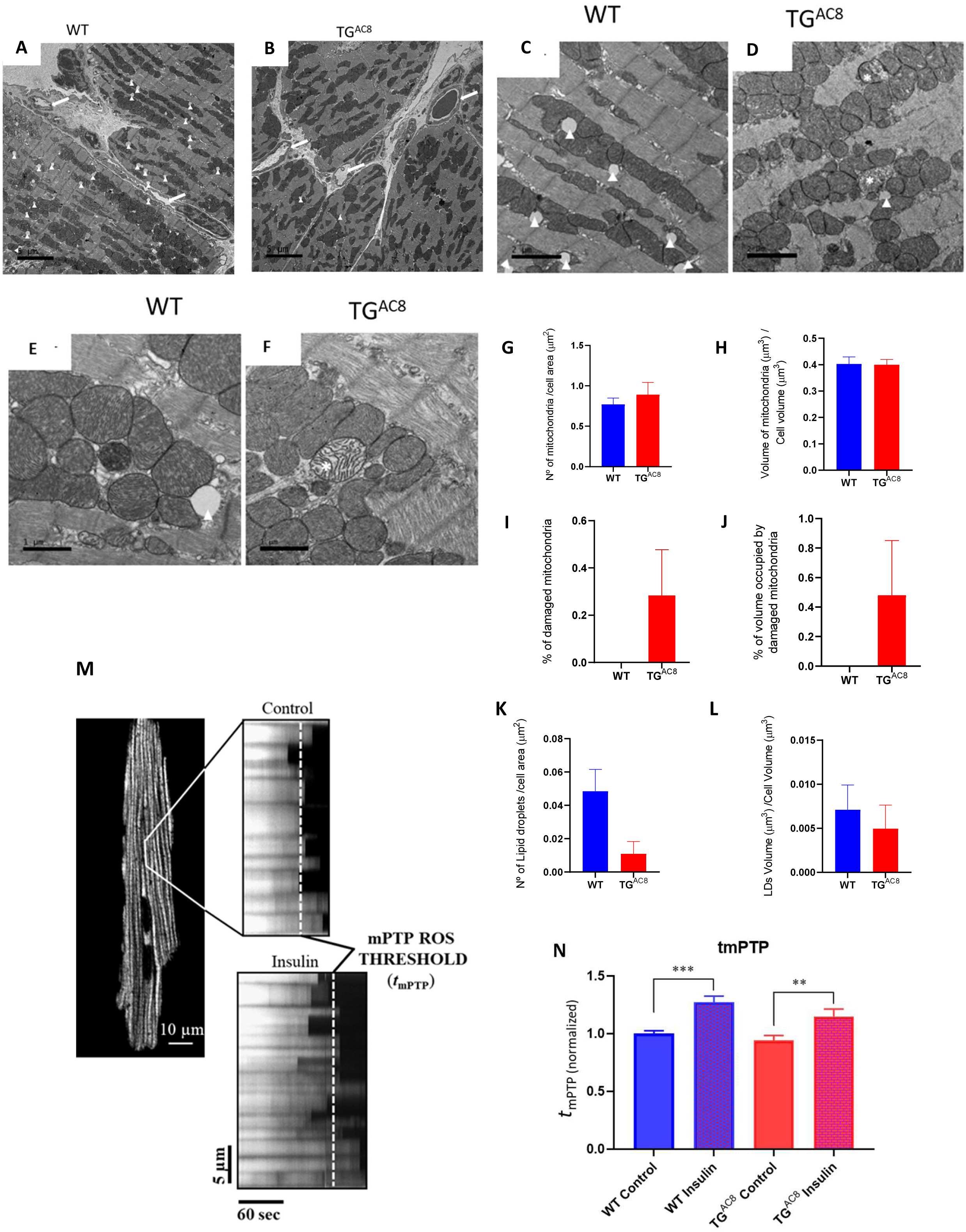
**(A, B)** Representative panoramic electron micrographs and **(C-F)** higher resolution images of LV cardiac muscle fibers and mitochondria in TG^AC8^ and WT. White arrowheads depict lipid droplets; asterisks show swollen, disrupted mitochondria with lighter cristae compared to the surrounding, healthy mitochondria. **(G, H)** Average mitochondrial number of quantitative stereological analyses of normal mitochondrial number and volume, **(I, J)** damaged mitochondria**, (K, L)** number of lipid droplets per cell area and volume of lipid droplets. **(M, N)** mPTP-ROS threshold, measured in a single LV cardiac myocyte, did not differ between TG^AC8^ and WT mice. Insulin was employed as a positive control. (N=3 in each genotype) (** p<0.01, *** p <0.001).

### Mitochondrial Fitness

The healthy functioning and survival of cardiac myocytes during severe, chronic myocardial stress requires close coordination of survival mechanisms and numerous mitochondrial functions that require a high level of mitochondrial fitness.^23–26^ The mitochondrial permeability transition pore (mPTP) is a key regulator of mitochondrial functions, including energy metabolism (e.g., with the pore performing as a “safety valve”, opening transiently and reversibly, to prevent: 1) the excessive accumulation of certain regulatory species, such as Ca^2+^; and 2) bioenergetic byproducts/damaging reactive species, such as free radicals, from achieving toxic levels. The mPTP also regulates cell fate: enduring and irreversible pore opening, plays decisive mechanistic roles in mitochondrial and cell life vs death decisions during normal development or pathological stress (e.g., involving excess and damaging free radical exposure). Measurement of the pore susceptibility or resistance to being induced/opened can serve as a biomarker of mitochondrial fitness. **Fig. 7** panels **M** and **N** shows that the mPTP ROS threshold did not differ in TG^AC8^ vs WT, suggesting a comparable degree of mitochondrial fitness in both genotypes.

### ROS levels and NRF Signaling

Given the incessantly elevated myocardial contractility and heart rate, and increased protein synthesis and quality control mechanisms in TG^AC8^ vs WT, it might be expected that ROS levels are increased in TG^AC8^. To this end we measured of superoxide radical accumulation in isolated, perfused, isometrically contracting TG^AC8^ hearts that maintained markedly enhanced cardiac workload observed *in vivo* (**Fig. 8 A-B)**. Superoxide radical accumulation did not differ by genotype, suggesting that mechanisms to scavenge ROS are increased in TG^AC8^. Indeed, the level of NRF2 protein, a key regulator of ROS defense signaling was increased by 24% in TG^AC8^ vs WT, suggesting that increased NRF signaling in the TG^AC8^ LV may be a factor that prevents the accumulation of superoxide ROS.

**Fig. 8.**
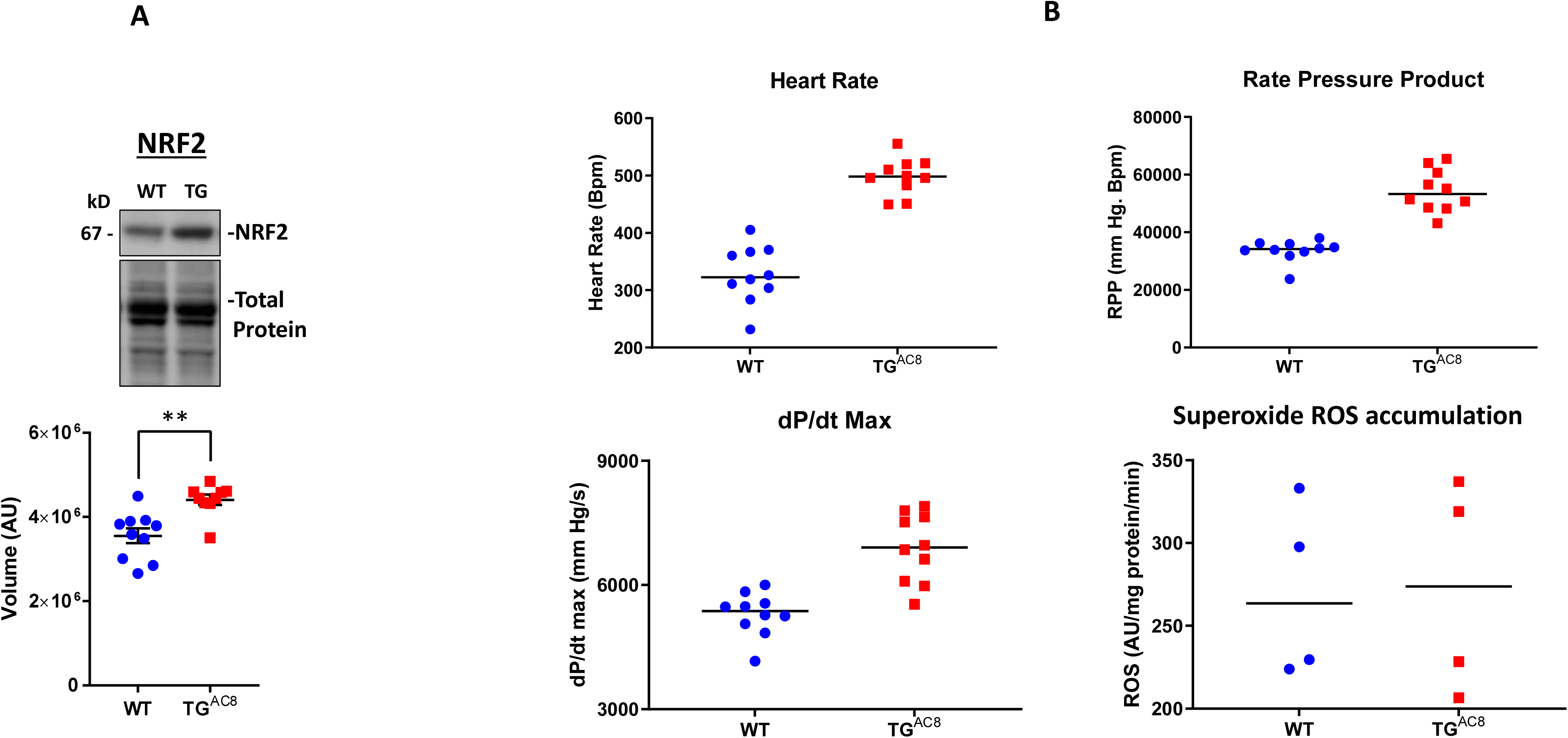
(A) WB analysis of Nrf2 expression in LV TGA^C8^ vs WT. (B) LV performance and the rate of superoxide (ROS) generation in isolated working TG^AC8^ and WT hearts.

### High Energy Phosphates

Given the fact, that increased protein synthesis and quality control mechanisms **(Fig. 6**), maintenance of normal ROS levels (**Fig. 8**) and the incessant high cardiac performance of the TG^AC8^ (**Fig. 1)** require increased energy production, it is reasonable to assume that the total energy requirements of the TG^AC8^ LV are probably considerably higher than those in WT. It was important, therefore, to assess high energy phosphate levels in TG^AC8^ and WT. A schematic of ATP-creatine energy system is depicted in **Fig. 9A**. Steady state levels of ATP, phosphocreatine and the ATP: phosphocreatine assessed in *vivo,* were maintained at the same level in TG^AC8^ as in WT (**Fig. 9 B-E**), suggesting that the rate of ATP production in the TG^AC8^ LV is adequate to meet its increased energy demands at least when animals rest.

**Fig. 9.**
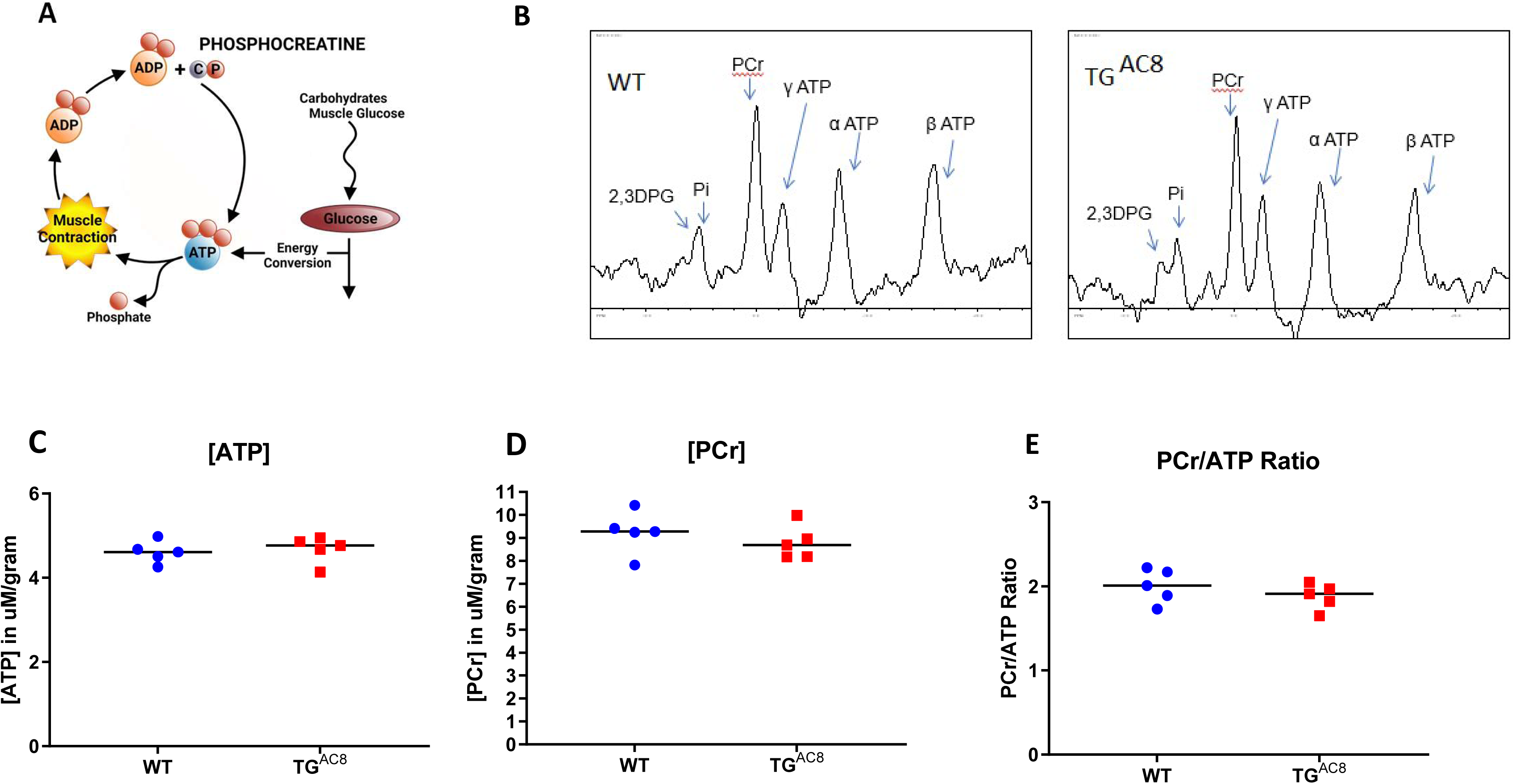
(A) A schematic of ATP creatine energy system, (B) Representative P31 NMR spectra of TGAC8 and WT hearts. (C-E) average levels of ATP, PCr, and ATP/PCr in TGAC8 and WT hearts derived from NMR spectra.

### Unbiased RNASEQ and Proteome Analyses

We performed unbiased, RNASEQ and proteome analyses of adult TG^AC8^ and WT left ventricles (LV) in order to realize facets of the consilience of adaptive mechanisms beyond those, identified in our experiments illustrated in **Fig.s. 1-9**. We reasoned that taking advantage of the knowledge base within bioinformatic tools would also generate a number of testable hypotheses regarding the components of adaptive paradigm of heart performance and protection in response to the constitutive challenge of marked cardiac-specific overexpression adenylyl cyclase type 8.

### LV Transcriptome (RNASEQ)

RNA sequencing identified 11,810 transcripts in LV lysates (**Fig. 10 A**, **Table S.2**); of these, 2,323 were differentially expressed in TG^AC8^ vs WT (**Fig. 10 A** and T**able S.3**): 1,201 were significantly upregulated in TG^AC8^ vs WT, and 1,117 were significantly downregulated. A volcano plot and heatmap of these transcripts are shown in **Fig. S.4 panels A and C**. The transcript abundance of human *ADCY8* in TG^AC8^ (LV) myocardium was more than 100-fold higher than the endogenous mouse isoform (**Fig. S.4 E**).

**Fig. 10.**
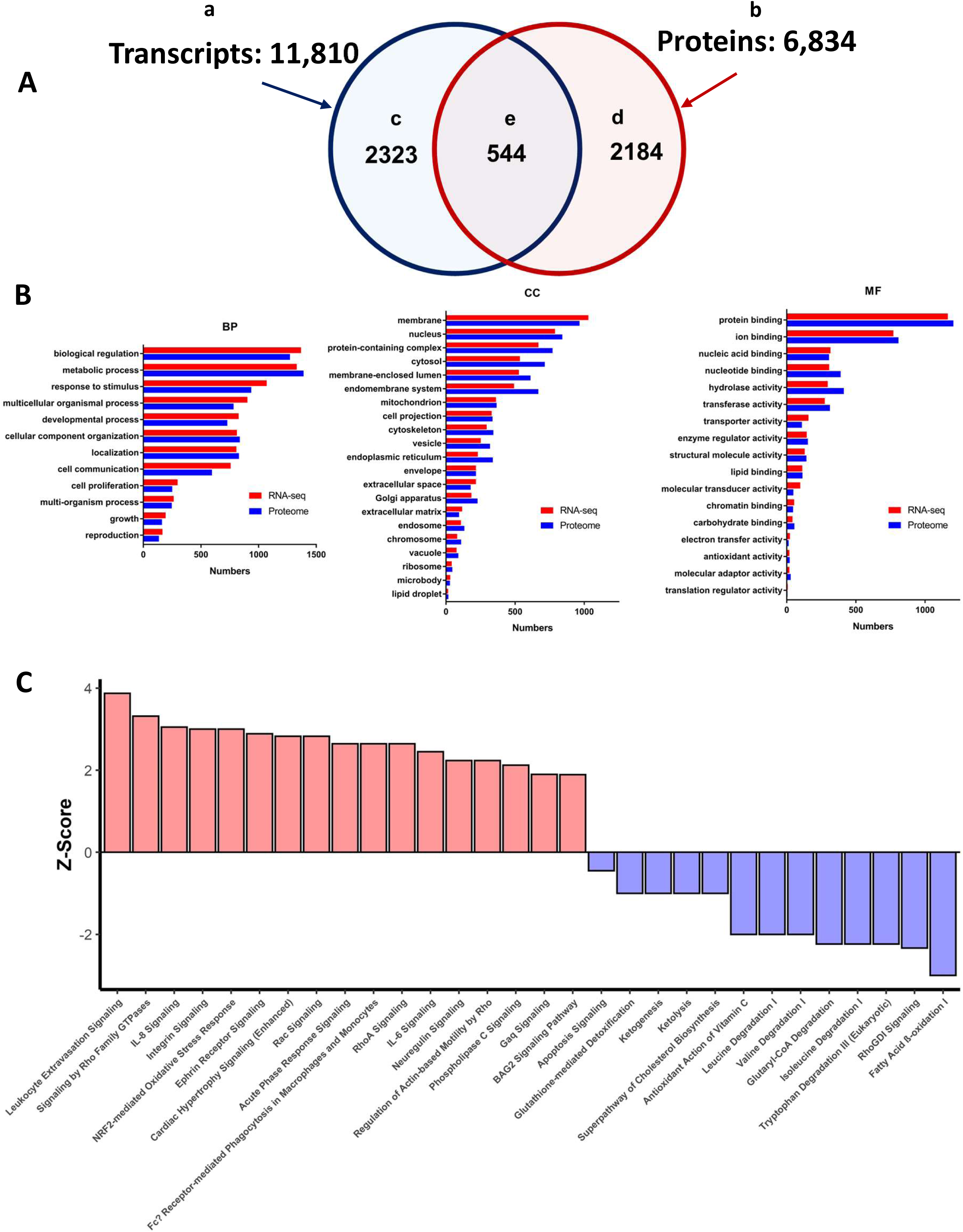
(A) Schematic of the total number of transcripts (subset “a” – 11,810), and proteins (subset “b” 6,834), identified in LV lysates, the number of transcripts (subset “d” 2,323), and proteins (subset “d” 2,184), that differed by genotype, and number of identified transcripts and proteins that both differed by genotype (subset “e” - 544). (B) WEBGESTALT analysis of the XX transcripts (Panel A subset “c”) and YY proteins (Panel A subset “d”) that significantly differed by genotype. Biological Processes (BP), Cell Compartment (CC), Molecular Functions (MF). (C) Top canonical signaling pathways differing in enrichment (-log(pvalue) >1.3) and activation status by genotype (Z-score: Fig. key) in IPA analysis of transcripts and proteins.

### The LV Proteome

6,834 proteins were identified in the LV proteome (**Fig. 10 A**, **Table S.4**); of these, 2,184 were differentially expressed (**Table S.5**) in TG^AC8^ vs WT: 2,026 were upregulated and 158 were downregulated in TG^AC8^. A Volcano plot and heatmap of the proteome are shown in the **Fig. S.4** panels **B and D**.

### Bioinformatic Analyses of the *Total* Transcriptome and Proteome Datasets

We employed WEB-based GEne SeT AnaLysis Toolkit (WebGestalt) and Ingenuity Pathway Analysis (IPA), online platforms for in depth analyses of the total 2,323 expressed transcripts (**Fig. 10 A-a**) and the total 2,184 **(Fig. 10 A-b)** proteins that differed by genotype.

In Gene Ontology (GO) analysis WebGestalt classifies submitted molecules into three main categories, i.e., Biological Process (BP), Cellular Component (CC), Molecular Function (MF), (**Fig. 10 B)**. Remarkably, transcriptome and proteome gene ontology (GO) terms that differed by genotype in response to overexpression of a single gene, AC Type 8, covered nearly all the biological processes and molecular functions within nearly all compartments of the TG^AC8^ LV myocardium.

In order to discover genotypic differences in canonical signaling pathway enrichment, we performed QIAGEN Ingenuity Pathway Analysis (IPA) of the total number of transcripts (2,323) that were differentially expressed by genotype **(Fig. 10 A-c)** and the total number of proteins (2,184 **Fig. 10 A-d**). In IPA analyses 248 canonical pathways in the transcriptome analyses and 308 canonical pathways in the proteome differed by genotype p<0.05 in *enrichment* (listed in **Tables S. 6A, B**). 154 pathways in transcriptome or proteome were *activated* by genotype (Z-score calculated) and Spearman’s correlation coefficient r_s_ was 0.57, p < 0.0001 **Fig. S.5**, and showed moderate correlation between enriched and activated pathways in transcriptome and proteome Although 60 molecules were represented in only a single enriched pathway, 22 molecules were represented in 10 or more pathways, the highest among these being: MAP3K – (68 pathways); MAP2K2 – (51 pathways); and PLCG2 – (33 pathways) (**Table S.6C**). Representation of several differentially expressed molecules in multiple canonical signaling pathways that differed by genotype is a plausible explanation accounting for the consilient pattern of pathway enrichment and activation in the TG^AC8^ heart identified in IPA analyses.

The top enriched and activated or inactivated pathways are illustrated in **Fig. 10 C**. Of note, several pathways, e.g., Nrf2 signaling, Integrin Signaling, and Cardiac Hypertrophy Signaling were among the top activated pathways that were highly enriched in TG^AC8^ vs WT, whereas fatty acid β-oxidation, tryptophan degradation III, isoleucine degradation and glutaryl Co-A degradation were markedly suppressed in TG^AC8^ vs WT (**Fig. 10 C**). Surprisingly, the top differentially regulated pathways between TG^AC8^ and WT was “leukocyte extravasation signaling” (**Fig. 10 C**).

### Bioinformatic Analyses of Transcripts and Proteins that *Both* Significantly Differed by Genotype

544 transcripts and proteins **both** differed by genotype **(Fig. 9A-e, Table S.7A)**. Of these 544, 339 (62.32%) were significantly **upregulated** in TG^AC8^; and 99 (18.2%) were significantly downregulated in TG^AC8^. Thus, of the 544 transcripts and proteins that **both** significantly differed by genotype, 80.5% differed **concordantly** (in the same direction, **Fig. S.6**).

We next subjected these 544 molecules to IPA analysis. 170 of the **same** canonical pathways in the transcriptome and proteome differed by genotype in enrichment. Of these, 118 pathways also differed by genotype and activation status (**Table S.7B**).

In addition to IPA analyses we also used PROTEOMAP platform^27^ to visualize functional categories of the 544 proteins that differed by genotype in **Fig. 10 A**-**e, Fig. S.6**. PROTEOMAP displays a protein data set as functional trees (**Fig. S.7**), each consisting of a number of polygons, the areas of which reflect genotypic differences in protein abundances. Genotypic differences were most marked by increases in TG^AC8^ of: ENVIRONMENTAL INFORMATION PROCESSING (informing on marked differences by **Rap1** and **PI3K-AKT** signaling pathways, harboring the marked increases in expression of ADCY8 and TNC (tenascin) proteins); GENETIC INFORMATION PROCESSING (informing on PROTEIN **translation, folding, sorting and degradation** signaling pathways, harboring marked increases in RNF128, MMP2 and WFS1); and CELLULAR PROCESSES (informing on **vesicular transport** and **exosome**, harboring increased NCF1, LCP1 and CTSZ proteins).

PROTEOMAP also identified major genotypic changes in BIOSYNTHESIS AND CENTRAL CARBON METABOLISM (**Fig. S.7**), informing on genotypic differences in: **membrane transport,** harboring increased SLC4A2 (regulates intracellular pH, biliary bicarbonate secretion, and chloride uptake); **other metabolic enzyme proteins** (PDK3, SULF2 and ALPK2), and carbohydrate metabolism; and **glycolysis**, harboring reduced ALDH1B1.

### Regulatory Networks Centered on cAMP and Protein Kinase A Signaling

Having confirmed that both AC8 protein and AC activity were markedly increased in TG^AC8^ vs WT (**Fig. 4 B, C**) it was reasonable to infer that the multitude of genotypic differences that were identified in WebGestalt, IPA and PROTEOMAP (**Fig.s. 10 B, S6; Table S.6**) might be ultimately (directly or indirectly) linked to increased signaling driven by the high levels of AC activity. Three main downstream targets of cAMP generated by AC activity are: protein kinase A (**PKA**), guanidine-nucleotide exchange proteins activated by cAMP (**EPACs**), and cyclic nucleotide-gated ion channels (**CNGCs**). We found no evidence of activation for 2 of these 3 targets. In fact: 1) Neither CNGC subunits α 1-4, nor β 1-2 transcripts or proteins were identified in our omics analysis; 2) omics analysis (**Table S.2, S4**) provided no evidence to suggest that cAMP directed **Epac** signaling was upregulated in the TG^AC8^ LV. More specifically, transcripts of *Rapgef* 1 thru 6 were significantly **downregulated** in TG^AC8^ vs WT (**Table S.3**) and RAPGEF 2, 3 and 5 proteins identified in proteomic analyses did not differ by genotype (**Table S.4**). On the contrary, we found clear evidence of activation of PKA. In fact, we had shown that PKA catalytic subunit and PKA activity are substantially higher in TG^AC8^ vs WT (**Fig. 4 D, F**), and PKA signaling was among the top pathways increased and activated in IPA analysis We therefore next focused the IPA knowledge base on the PKA complex as the center of a number of **interacting** pathways. We noticed that in addition to cAMP, another upstream regulator of PKA signaling, ITGA5, was increased in TG^AC8^ vs WT in both the transcriptome and proteome (**Fig. 11**), suggesting increased crosstalk between intra and extra cellular signaling.

**Fig. 11.**
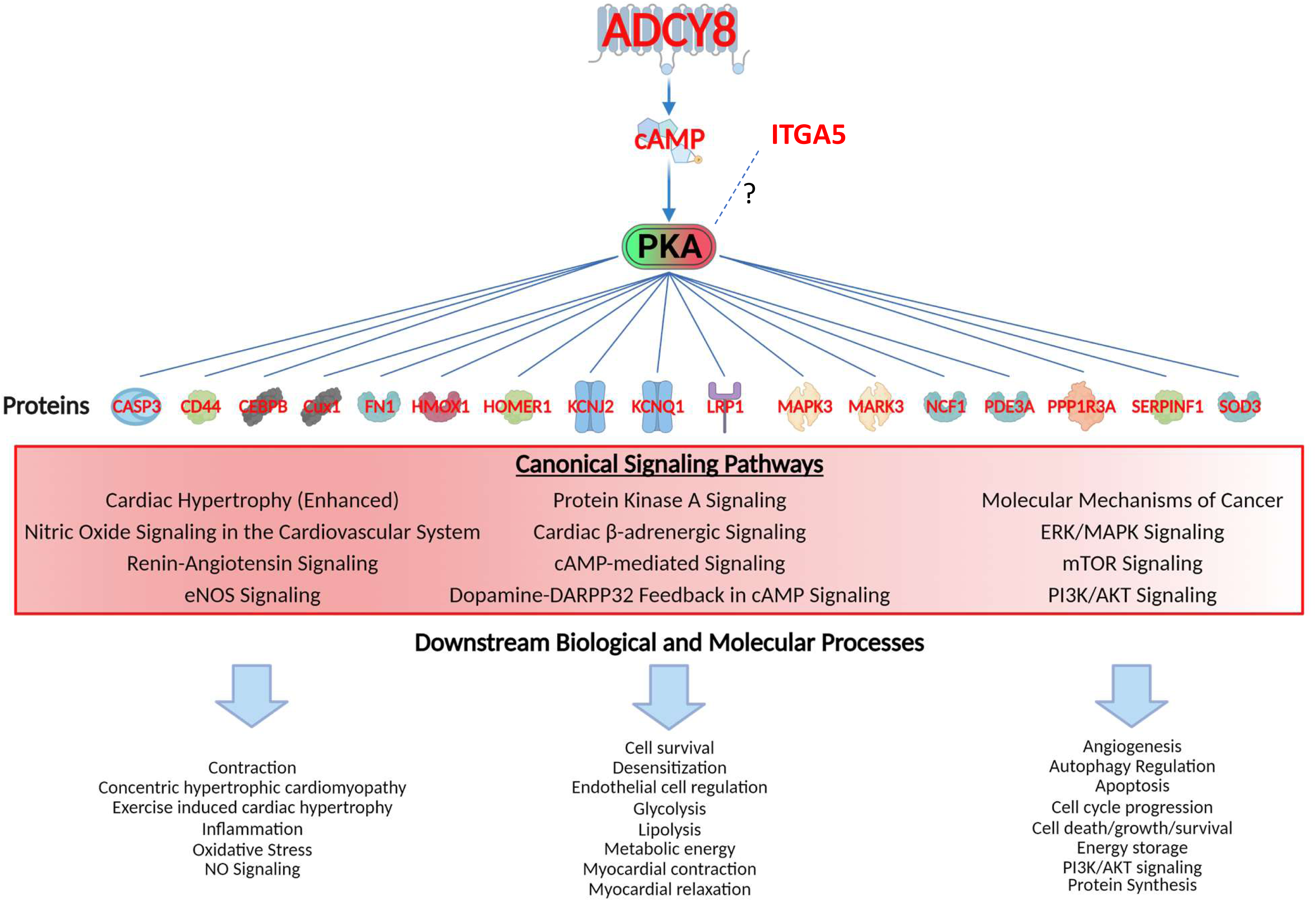
Regulatory Networks Centered on Adenylyl Cyclase and Protein Kinase A Signaling. Gene families of proteins regulated by PKA ranged from transcription regulators, kinases, peptidases and other enzymes, transmembrane receptors, ion channels and other gene families. Canonical signaling pathways in which these proteins operate and downstream biological and molecular processes of proteins in these pathways are also displayed (lower part). See **Table S.8** for the full list of these downstream effects of PKA signaling.

To further investigate the role of PKA-signaling in the extreme TG^AC8^ phenotype, we next established the protein interaction network centered on PKA (**Fig. 11**). Gene families of these proteins ranged from transcription regulators, kinases, peptidases and other enzymes to transmembrane receptors, ion channels and other protein families (CFH, CD44, HOMER1, and SERPINF1). Although CREB1 protein was not identified in proteome analyses, as noted this canonical transcription factor, and its phosphorylated form activated by PKA were markedly increased in WB analyses (**Fig.s. 5 D**, **S.11 B**). Canonical signaling pathways in which these proteins operate and downstream biological and molecular processes of proteins in these pathways are listed in **Fig. 11** (lower part). See **Table S.8** for the full list of these downstream effects.

### Shifts in Metabolism

The enrichment or activation of the large number of signaling pathways in TG^AC8^ vs WT (**Fig. 10 C, S.6; Table S.6 A, B**) suggested that some aspects of metabolism differed by genotype. A detailed schematic of the genotypic differences in transcripts and proteins that related to metabolism circuits is illustrated in **Fig. 12**, and WB validations of selected proteins are presented in **Fig. S.8.** Bioinformatic analyses suggested that nutrient sensing pathways, including AMPK, insulin and IGF signaling, that induce shifts in aerobic energy metabolism were activated in TG^AC8^ vs WT LV. A close inspection of the bioinformatic analyses of central carbon metabolic processes (**Fig. 12**) suggested that catabolism of glucose is markedly increased in TG^AC8^ vs WT, while the fatty acid β oxidation pathway is concurrently reduced.

**Fig. 12.**
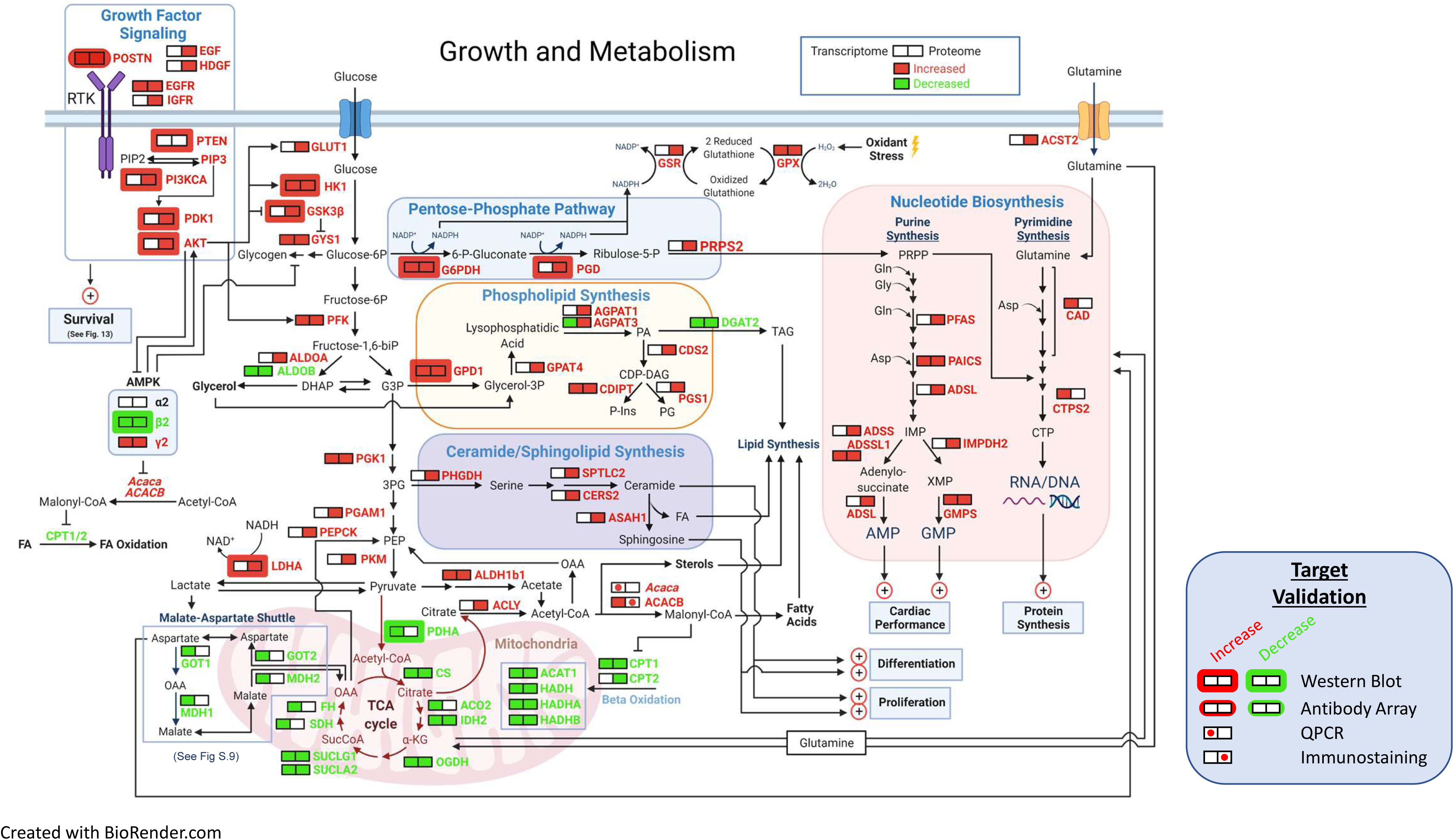
Growth and Metabolism. A schematic of growth and metabolism circuitry based on signals derived from bioinformatic analyses of the transcriptome and proteome and on selected WBs. Catabolism of glucose is markedly increased in TG^AC8^ vs WT as reflected in increased expression of Glut1, HK1 and PFK and other glycolytic enzymes, whilst fatty acid β oxidation pathway is concurrently reduced, as reflected in reduced expression of Cpt1, Cpt2, Acat1, Hadh, Hadha, and Hadhb. in TG^AC8^ vs WT LV. Shifts in the transcription of genes and translation of proteins that are operative within PPS pathway, i.e. G6PDH, PGD and PRPS2 suggests that PPS is more highly activated in the TG^AC8^ vs WT. The combination of increased expression of the glucose transporter, lactic acid dehydrogenase type A and the glutamine transporter in TG^AC8^, suggests that, relative to WT, the TG^AC8^ heart utilizes aerobic glycolysis to fulfill part of its energy needs. Enhanced growth factor and other PI3/AKT driven signaling processes increased in TG^AC8^ vs WT are also known to be linked to aerobic glycolysis.

An increase in myocardial glucose metabolism in TG^AC8^ vs WT was suggested by increases in GLUT1, HK1, GSK3Β, GYS1 and PFK, PGK1, PGA and PKM. Interestingly, bioinformatic analyses suggested that, both transcript (down by 86%) and protein (down by 38%) expression of ALDOB were reduced in TG^AC8^ vs WT. Signals related to reduced fatty acid oxidation (FAO) likely stem from reductions in CPT1, CPT2, ACAT1, HADH, HADHA, and HADHB (**Fig. 12**). Further, signals from bioinformatic analyses pointed to reduced utilization of the TCA cycle and mitochondrial respiration within the TG^AC8^ LV vs WT **(Fig. 12)**. Interestingly, omics signals suggested that the malate-aspartate shuttle (MAS), which results in the net transport H+ from cytosol back to the mitochondria, may be suppressed in TG^AC8^ vs WT, because transcripts of *Ldhb, Mpc1, Mpc2, Mdh1, Got1, Slc25a11 Slc25a13*, and SLC25A13 protein was also downregulated (**Fig.s. 12, S.9**). Reduced shuttling of aspartate into mitochondria in conjunction with increased level of ACC2 (**Fig. S.10 A, B**) would favor with increased nucleotide synthesis. ^28, 29^

The results of bioinformatic analyses also pointed to increased expression, in TG^AC8^ vs WT, of the glucose transporter, lactic acid dehydrogenase type A and the glutamine transporter, that are required for enhanced aerobic glycolysis^30^ (**Fig. 12**). **Enhanced growth factor** and other PI3K/AKT driven signaling processes in TG^AC8^ suggested by omics analyses (**Fig. 12)** and validated by WB (**Fig. S.8)** are tightly linked to aerobic glycolysis and utilization of pentose phosphate shunt (PPS).^30^ Increased utilization of the PPS in TG^AC8^ vs WT is suggested by increases in both the expression of transcripts and proteins of G6PDH, PGD and PRPS2 (**Fig. 12**).

Because Acetyl-CoA carboxylase (ACC), a complex multifunctional enzyme system that catalyzes the carboxylation of acetyl-CoA to malonyl-CoA, which limits CPT1 transport of FA into the mitochondria, we inquired whether, in addition to reduction in CPT1 transcripts and reductions in CPT1 and CPT2 proteins in TG^AC8^ vs WT, ACC2 might also be increased in TG^AC8^ LV myocytes. Indeed, immunolabeling of isolated LV cardiomyocytes indicated a clear increased in ACC2 (AKA ACACB) protein expression (**Fig. S.10 A**); and in increase *Acc1* (AKA *Acaca*) mRNA was documented in LV tissue by RT-qPCR (**Fig. S.10 B**). Such an increase of ACC2 expression in TG^AC8^ may not only explain a shift from FAO to glucose utilization within the TG^AC8^ LV but may also explain increased utilization of Aerobic glycolysis in TG^AC8^ vs WT, which is linked enhanced utilization of PPS to increase anabolic processes such as nucleotide synthesis (**Fig. 12**). Because it is known that *Acc2* deletion increases FAO and suppresses glucose utilization in the context of LV pressure overload, ^28, 29^ it is important to recall that cardiac myocytes within TG^AC8^ LV are smaller in size, and not enlarged as those in pathologic cardiac hypertrophy in response to LV pressure overload, ^31^ and in which ACC2 promotes glucose utilization and reduces FAO.^28, 29^ In other terms, an increase in ACC2 appears to promote enhanced glucose utilization, enhanced aerobic glycolysis, enhanced utilization of the PPS, and reduced FAO, and may be involved in the reduction of average LV cardiac myocyte size in TG^AC8^, and the absence of markers of pathologic hypertrophy within the TG^AC8^ LV.

### A Schematic of the Circuitry of Enhanced Cardiac Performance and Adaptive Mechanisms within the TG^AC8^ LV Derived from Phenotypic Characterization and Signals from Bioinformatic Analyses

**Fig. 13** depicts a hypothetical scheme of physiologic performance and protection circuits that appeared to be concurrently engaged within the TG^AC8^ LV. Because many of the perspectives depicted in the scheme in **Fig. 13** were derived from bioinformatic analyses of cell lysates, it is not implied that all of these circuits are present within the same cell types that reside in the heart (see Discussion). Although the pathways/specific targets and the effector functions/outcomes culminating from the regulation of the components within the circuits in the pathways are represented according to published literature with respect to cardiac-specific context. The circuits (**Fig. 13**) are based upon defined LV phenotypic characteristics (**Fig.s. 1 - 9**, **Fig. S.1, S2, S.9** and **Table S.1**), on signals derived from genotypic differences in transcriptome and proteome and IPA analyses (**Fig.s. 10** and **11, Tables S.2-S.6**), on selected WB, RT-qPCR and immunolabeling analyses performed prior to and following transcriptome and proteome bioinformatic analyses, and WB analyses (**Fig.s. 2, 4-6, 8, S.10, S.15**), and on additional selected post-hoc WB, RT-qPCR (**Fig. S.8-S.12**), performed having visualized the consilience of pathways depicted within the circuit schematic in **Fig. 13**.

**Fig. 13.**
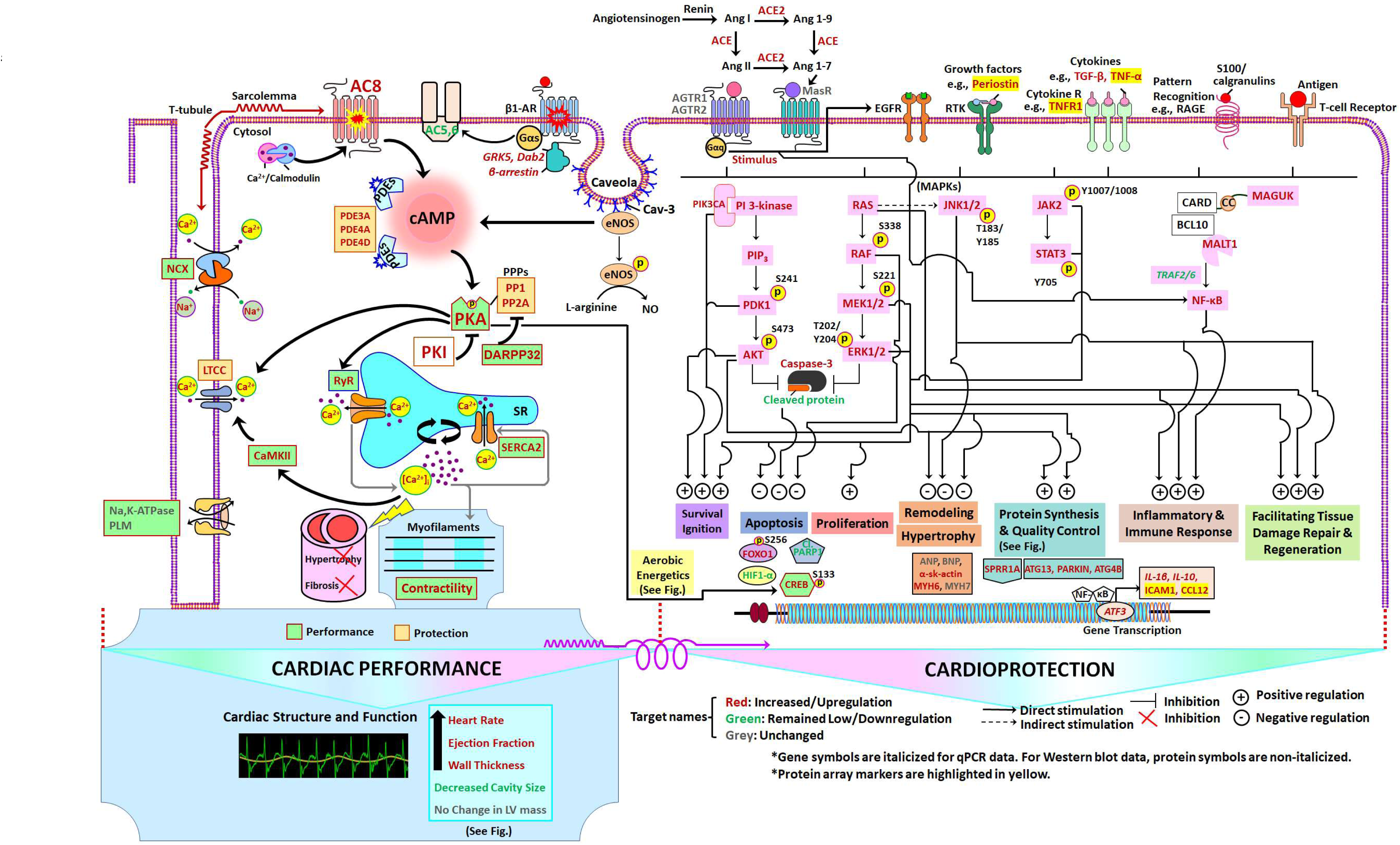
Schematic of TG^AC8^ Heart Performance and Protection Circuits that appeared to be concurrently engaged in the TG^AC8^ LV. The pathways/specific targets and the effector functions/outcomes culminating from the regulation of the components within the circuits in the pathways are represented according to published literature with respect to cardiac-specific context (See Discussion). **Pink colors** represent proteins that differed by genotype in WB. Molecular targets or components in **red**, **green,** and **grey** represent molecular targets or components that are increased or upregulated, decreased or downregulated, or unchanged, respectively, in TG^AC8^ vs WT.

## Discussion

### Cardiac Structure and Contractile Performance Circuitry

Because it is well known that alterations in heart structure and increased performance in the TG^AC8^ vs WT measured in vivo (**Fig. 1 and S.1, Table S1)** and depicted in the scheme in **Fig. 13** (left) can result from enhanced cAMP/PKA/Ca^2+^ signaling, we may conclude that the marked increases in AC8 protein level and activity and in PKA expression and activity in TG^AC8^ directly or indirectly lead to the differences in LV structure and performance in TG^AC8^ vs WT (**Fig. 4 A-F**).

### Left ventricular structure

Although the TG^AC8^ LV wall thickness was increased, neither LV mass assessed via echocardiograms nor post-mortem LV weight, differed from WT (**Fig. 1**), because the thicker LV walls in TG^AC8^ encompassed an LV cavity size that was markedly reduced at both end-diastole and end-systole compared to WT. Thus, because left ventricular hypertrophy, is strictly defined as an increase in LV mass, ^17, 18^ the TG^AC8^ LV is not technically hypertrophied. Furthermore, pathologic hypertrophy markers were not increased in TG^AC8^. It’s of interest, however, that KO of, Protein Tyrosine Phosphatase Non-Receptor Type 1 (PTP1B AKA PTPN1), which protects against pathologic hypertrophy and increase in fetal gene markers in response to transverse aortic constriction^32^ was increased by 2.5-fold in RNSEQ and 25% in proteome (**Table S.3, S.5**). The lack of an increase in LV mass in TG^AC8^ may be attributable, in part, to reduced LV wall stress (diastolic stretch), due to reduced filling volume, in TG^AC8^ vs WT, linked to the reduction in diastolic filling time, on the bases of the increased heart rate. Of note in this regard, the left atrial size was not enlarged in TG^AC8^ vs WT.

Histologic analysis of the thickened TG^AC8^ LV posterior wall revealed a cardiac myocyte profile that was shifted to cells that were smaller in size than WT (**Fig. 2**). Thus, an increased density (per unit area) of smaller cardiac myocytes contributed to the increase in LV posterior wall thickness in TG^AC8^. The LV wall collagen fraction did not differ by genotype (**Fig. S2**), even though TGF β protein, TGF β receptor, and downstream signaling molecules were all upregulated (**Fig. 13**). This suggests that additional adaptations that uncouple TGF β signaling from its effect increase fibrosis may become activated at this age in TG^AC8^ LV.

### Left Ventricular Contractile Performance

Neither the early, nor the late diastolic filling rates, or early/late ratio differed by genotype (**Table S.1**), indicating that the smaller LV EDV in TG^AC8^ vs WT was not associated with reduced diastolic functional measures. LV myocardial contractility was markedly increased in TG^AC8^ vs WT as evidenced by markedly increased EF *in vivo* (**Fig. 1**), as previously reported.^10^ A 30% increase cardiac output in TG^AC8^ vs WT is attributable to the 30% increase in heart rate, because stroke volume did not differ by genotype (**Fig. 1**).

As expected, expression of proteins that underly numerous cellular mechanisms driven by PKA signaling that determine cardiac myocyte contractile performance, were also upregulated in the transcriptome (**Table S.3**), proteome (**Table S.5**) or in both transcriptome and proteome **(Fig. S.5**, **Table S.7**) or in WB analyses (**Fig. 4 G-J**) in TG^AC8^ vs WT. Both KCNJ2 and KCNQ1 K^+^ channels which modulate cardiomyocyte membrane potential and action potential characteristics were upregulated in TG^AC8^ vs WT (**Fig. 11**): KCNQ1 transcripts increased by 2.3-fold and protein increased by 26%; KCNJ2 transcripts increased by 3.2-fold and protein 32% (**Tables S.3, S5**). Of note, KCNJ2 regulation is mechanical sensitive and its increase in TG^AC8^ likely relates, at least in part, to the hypercontractile state of the TG^AC8^ LV.

It bears emphasis that the cardiac structural and resting performance profiles of the TG^AC8^ LV (**Fig. 1**), therefore differ from those, observed in LV remodeling induced by chronic exercise endurance training, in which the diastolic cavity size, and stroke volume are increased at rest, ^33^ resting HR is reduced and cardiac output is unchanged. The cardiac myocyte profile in TG^AC8^ also differs from that induced in chronic exercise in which the LV myocytes size are increased but the myocyte number is unchanged.^34^ The marked differences in cardiac structure and function between TG^AC8^ and the endurance-trained heart, in large part, likely to be attributable to increased vagal tone at rest in the latter, but not in the former,^8^ in which an exercise functional profile is maintained around the clock at rest. In other terms the TG^AC8^ heart does not have an opportunity to rest between bouts of dynamic exercise as do the hearts of endurance trained organisms.

### Cell Cycle, Proliferation, and Growth and Developmental Circuits

PKA signaling links many pathways involved in protein synthesis, (**Fig. 11**) which was increased in the TG^AC8^ vs WT LV (**Fig. 5A)**. Regulation of the synthesis of numerous proteins related to cell cycle, cell proliferation, and growth and development were increased in TG^AC8^ (**Tables S.3, S.5, S.10**). For example, the *Mras* oncogene transcript increased by 40%, and as noted, RAS protein was increased by 40% in TG^AC8^ vs WT (**Fig. 5 B)**. Numerous molecules downstream of RAS were increased and activated in TG^AC8^ LV including: RAF, MEK1/2 and ERK1/2 **(Fig. 5 B, C**), indicating that MEK/ERK signaling, associated with enhanced cardiac proliferation^35^ is activated to a greater extent in TG^AC8^ compared to WT (**Fig. 13**, ERK arm).

AKT2 protein level and phosphorylation state were markedly increased in TG^AC8^ vs WT (see WB in **Fig. S.8 A**). Numerous growth factor signaling molecules upstream of AKT were upregulated in TG^AC8^ vs WT (**Fig. S.8, Fig.s. 12, 13**) including periostin, which was increased by more than 100% in the transcriptome, and by 39% in proteome **(Tables S.10)** and by antibody array (**Fig. S.14**). Of note, periostin plays a role in tissue regeneration, including wound healing, and ventricular remodeling following myocardial infarction. ^36–38^ *Egfr*, *Hdgf*, and *Igf1* transcripts were also increased in TG^AC8^, and IGF1R protein was also borderline increased (**Table S.10**). Interestingly, some features of endurance exercise conditioning involve upregulation of IGF signaling via PIK3CA.^39^ PIK3CA expression was increased in TG^AC8^ **(Fig. S.7 A)**, as well as IGF signaling (**Table S.6A, B**).

Other molecules related to cell cycle, cell proliferation and growth and development that were increased in TG^AC8^ vs WT included: CDK2 (increased by 19%), CDK6 (increased by 36%), NOTCH1 (increased by 12%), beta1 catenin protein (increased by 17%), and PCNA (increased by 12%), *Hand2* (increased by 100%), and *Tbx5* and *Tbx20* transcripts both increased by 30% **(Table S.10)**.

Nuclear EdU labeling in cardiac myocytes did not differ by genotype. EdU labeling of non-cardiac myocytes, however, was substantially increased (**Fig. 3**) in TG^AC8^ vs WT. Leading candidates of this regenerating cell network include endothelial cells, fibroblasts, immune cells, pericytes and most importantly, telocytes.^40^

### Shifts in Metabolism

A greater **rate** of energy production in TG^AC8^ vs WT is likely required to support, not only the higher HR and increased contractility during each heartbeat compared to WT, but also increased growth factor and other PI3K-directed signaling pathways, e.g. autophagy, protein synthesis and protein quality control (**Fig.s. 5, 6, 13**). The increased “energy requirements” of the TG^AC8^ LV are apparently being paid in full, because there is no indication that the TG^AC8^ LV is chronically hypoxic, as HIF-1α transcripts are down regulated (**Table S.2**) and HIF-1α protein level is reduced (**Fig. S.8 B**). Further, steady levels of ATP and phosphocreatine in the WT LV *in vivo* at rest did not differ in TG^AC8^ vs WT (**Fig. 9**). A prior study, in fact, indicates that exercise capacity of TG^AC8^ exceeds that of WT. ^41^

Bioinformatic signals suggested that catabolism of glucose is markedly increased in TG^AC8^ vs WT (**Fig. 12**). Enhancement of myocardial glucose catabolism was suggested by increases in GLUT1 (AKA Slc2a1) HK1 and PFK in TG^AC8^ vs WT LV (**Fig. 11**). Of note, GLUT1 is the embryonic type of glucose transporter; ^42, 43^ and not the canonical heart form, GLUT4 (AKA Slc2a4) observed in the adult heart in response to myocardial stress.^44, 45^

Phosphorylation of GSK3B by AKT1 deactivates GSK3B. Inactivation of GSK3B via its phosphorylation is a key conversion point of a numerous processes that confer cardioprotection. ^24^ Although WB analysis indicated that GSK3B phosphorylation did not differ between genotypes, total GSK3B protein was significantly increased in TG^AC8^ vs WT (**Fig. S.8 A**). Gys1 and phosphorylase kinase (PhK) were increased in the TG^AC8^ vs WT both the transcriptome and proteome (**Tables S.3 and S.5**) suggesting that a high turnover (increased synthesis and increased degradation) of glycogen may be involved in the marked increase in glucose catabolism within the in TG^AC8^ LV (**Fig. 12**). LV glycogen staining was modestly increased in the TG^AC8^ vs WT **(Fig. S.13 B).**

Enhanced growth factor and other PI3K/AKT-driven signaling processes in TG^AC8^ suggested by omics analyses and validated in **Fig. 12, 13, S.8, S.11** are tightly linked to ***aerobic glycolysis***.^30, 46^ It is well established that the **rate** at which ATP is generated in aerobic glycolysis is *greater* than that generated in oxidative phosphorylation via the TCA cycle and respiration.^30^

Bioinformatic analyses also indicated that in addition to increased expression of the glucose transporters, lactic acid dehydrogenase type A (cytosolic isoform of LDH) and the glutamine transporter, which are required for ***aerobic glycolysis***^30^ are also increased in TG^AC8^ vs WT (**Fig.s. 12, S.9**), suggesting that, relative to WT, the TG^AC8^ heart may utilize ***aerobic glycolysis*** to fulfill part of its energy needs. This idea is underscored by the absence of an increase in HIF1α (**Fig. S.11 B**), suggesting the absence of hypoxia in the TG^AC8^ LV.^47, 48^ Aerobic glycolysis promotes utilization of the pentose phosphate shunt (PPS). PPS activation facilitates amino acid and nucleotide synthesis. Overexpression in TG^AC8^ LV of genes and proteins that are operative within the glucose metabolic pathway, e.g. G6PDH, PGD and PRPS2. (**Fig.s. 12, S.8 C**), suggests that the PPS is more highly utilized in the TG^AC8^ vs WT LV.

Enhanced utilization of PPS in TG^AC8^ may be involved in re-synthesis of crucial amino acids and nucleotides following their degradation, (via increased autophagy, mitophagy and proteosome activity in TG^AC8^ vs WT, **Fig. 6 D, G, I, J)**. PPS and may also be involved in catalyzing the replenishment of NADPH in TG^AC8^ via LDH and increased G6PDH and PGD (**Fig. 12**). Further ACLY, which catalyzes the **synthesis** of fatty acids (FA), leading to the formation of AcCoA, linking to synthesis of FA, lipids, phospholipids, and ceramides, was increased in TG^AC8^ vs WT proteome analysis (**Fig.s. 12, S.7, Table S.5**). Enhancement of these processes in TG^AC8^ via increased utilization of PPS may be linked to signals pointing to increased cell cycle, cell proliferation and growth and development in TG^AC8^ vs WT. In this context aerobic glycolysis and increased utilization of PPS in TG^AC8^ resembles heart **embryonic** differentiation and growth prior to the onset of the fetal stage when respiratory enzymes, e.g., COX7 are induced.^49^ Thus, increased utilization of the PPS, in TG^AC8^ may be a crucial factor that underlies increased cardiac biomass (thicker LV walls vs WT containing an increased number of small myocytes, and an increased number of EdU-labeled non-cardiac myocyte nuclei, **(Fig. 1 and 2)** in the absence of increased collagen deposition (**Fig. S.2**)).

Bioinformatic signals suggested that the fatty acid β oxidation pathway and pathways that degrade branched chained amino acids are concurrently reduced, possibly due to reductions in CPT1, CPT2, ACAT1, HADH, HADHA, and HADHB (**Fig. 12**). Bioinformatic analyses also suggested that utilization of TCA cycle and mitochondrial respiration within the TG^AC8^ LV are reduced compared to WT, even though the mitochondrial number and volume do not differ by genotype (**Fig. 7 G, H**).

Downregulation of transcripts or proteins operative within the MAS, (*Ldhb, Mpc1, Mpc2, Mdh1, Got1, Slc25a11 Slc25a13*, and SLC25A13) **Fig. 12 and Fig. S.9** suggested that in conjunction with reduced TCA cycle and mitochondrial respiration in TG^AC8^ vs WT the translocation of H+ from cytosol to mitochondria to restore the level of mitochondrial NADH is also reduced in TG^AC8^. An interaction between cytosolic lactate and the MAS has been proposed: cytosolic lactate formed during glycolysis may be used as a mitochondrial energy source via translocation of lactate via lactate-MAS within the same cell. Lactate can also be exported from one cell and taken up by other cells. ^50^ In heart, lactate can also regenerate cytosolic NAD^+^ from cytosolic LDH.^51, 52^ Under conditions when shuttling of NADH from cytosol into the mitochondria via the MAS is limited, an increase in cytosolic lactate concentration might be expected to favor cytosolic NAD^+^ regeneration via the LDH reaction. Thus, it might be expected that persistent high cardiac workloads that are characteristic of TG^AC8^ heart, in conjunction with reduced MAS and reduced expression of proteins involved in TCA cycle and mitochondrial respiration in TG^AC8^ vs WT, are associated with increased replenishment of cytosolic NADH from NAD+, at the expense of replenishment of mitochondrial NADH from NAD^+^ within mitochondria. It is important to note that both transcripts and protein levels of IDH, which catalyze the oxidative decarboxylation of isocitrate to α-ketoglutarate, are reduced in TG^AC8^ vs WT, because during high cardiac workloads, as in the TG^AC8^ heart, the α-ketoglutarate/malate transporter located within the inner mitochondrial membrane does not compete favorably with mitochondrial matrix α -ketoglutarate dehydrogenase for α-ketoglutarate their common substrate.^52^ It is well known that even in the presence of adequate oxygen during strenuous exercise requiring increased energy utilization, plasma lactate concentration becomes increased, suggesting that cytosolic lactate is increased^53^ in the TG^AC8^ heart, favoring NAD^+^ regeneration in the cytosol. ^54^

### ROS Regulation

Although increased rates and amounts of ATP production and utilization in the TG^AC8^ heart to support simultaneous increased cardiac performance and growth factor signaling are likely to be associated with a substantial increase in ROS generation, ROS levels were not increased in the TG^AC8^ LV (**Fig. 8 B**). Because ATP production via aerobic glycolysis generates less ROS compared respiration,^30^ increased utilization of aerobic glycolysis would be one mechanism to limit ROS production within the TG^AC8^ LV. Other mechanisms to reduce ROS levels that appeared to be utilized to the greater extent in TG^AC8^ vs WT include NRF2 signaling, one of the top enriched and activated pathways **(Fig.s. 8 A, 10 C),** SOD3 (transcripts increased by 3.2-fold and protein by 20%), and HMOX1 (transcripts increased by 54% and protein by 32%; **Table S.7**) which not only protects tissue against oxidative stress, but also protects against apoptosis.^55, 56^ HMOX1 is an important activator of NRF1, which has been recently discovered to be involved in heart regeneration and repair, promoted by NRF1 signaling.^57^ Importantly, four members of the family of glutathione peroxidases involved in the termination reaction of ROS pathways were increased in TG^AC8^ vs WT: Gpx3 (transcripts by 103%, proteins by 30%); Gpx7 (transcripts by 40%, proteins by 21%); Gpx8 (transcripts by 20%, proteins by 16%); Gpx1 (transcripts by 39%, proteins by 12%). Several members of glutathione transferases were differentially expressed in TG^AC8^ vs WT: Gsta3 (transcripts and proteins upregulated by 55% and 26% respectively); GSTM5 (proteins upregulated by 23%), GSTM1 (proteins upregulated by 14%); mitochondrial glutathione reductase GSR was also upregulated, while GSTM7 (transcripts and proteins were decreased by 21%, and 9%) and GSTO1 (protein was decreased by 9%).

### Cardiac Protection Circuitry

#### Negative feedback on βAR-cAMP-PKA-Ca signaling

Numerous molecules that **inhibit** βAR signaling, (e.g. Grk5 by 2.6 fold in RNASEQ and 30% in proteome; Dab2 by 1.14 fold in RNASEQ and 18% in proteome; and β-arrestin by 1.2 fold in RNASEQ and 14% in proteome) were upregulated in the TG^AC8^ vs WT LV (**Table S.3, S.5** and **S.9)**, suggesting that βAR signaling is downregulated in TG^AC8^ vs WT, and prior studies indicate that βAR stimulation-induced contractile and HR responses are blunted in TG^AC8^ vs WT.^8, 11^ The heart, itself, produces catecholamines, (**Fig. S.3 A**). It is interesting to note that βARs become desensitized in TG^AC8^ (**Fig. S.3 B**), even though neither plasma norepinephrine, nor epinephrine are increased, but reduced in TG^AC8^ vs WT^8^ (**Fig. S.3 C**).

Both the RNASEQ and proteome analyses indicated that PI3K/AKT signaling, which promotes survival,^58^ is activated in TG^AC8^ vs WT (**Fig. 11** and **13**). Numerous studies have indicated that PI3K signaling is involved in βAR internalization,^59, 60^ and is also involved in cAMP metabolism by acting as an essential component of a complex controlling PDE3B phosphodiesterase-mediated cAMP destruction.^61, 62^

In cultured cardiomyocytes, βAR stimulation increases PI3K activity.^63, 64^ Both the β1AR and β2AR have been reported to transactivate PI3K in vitro.^65, 66^ Moreover, βAR stimulation-induced increases in heart weight, contractile abnormalities, and myocardial fibrosis, and cardiac “fetal” genes were markedly attenuated in the absence of PI3K expression.^67^ Although this formidable evidence in most cases has identified PI3Kγ to be specifically involved in cardioprotection with response to G-protein coupled receptor signaling, PI3Kγ was not identified in our transcriptome or proteome analyses, but RT-qPCR assay detected significant increase of catalytic subunit of PI3Kγ (*Pi3kcg)* expression (**Table S.9**). Tyrosine receptor coupled PI3KCA, AKT and PDK1 were increased in TG^AC8^ vs WT proteome, and these proteins, as well as PTEN (WB) and PIP3 (ELISA) were increased TG^AC8^ vs WT (**Fig. 12, Fig. S.7**).

A blunted response to βAR stimulation in a prior report was linked to a smaller increase in L-type Ca^2+^ channel current in response to βAR stimulation in the context of increased PDE activity.^13, 14^ WB analyses showed that PDE3A and PDE4A expression increased by 94% and 36%, respectively in TG^AC8^ vs WT, whereas PDE4B and PDE4D did not differ statistically by genotype (**Fig. S.11 A**). In addition to mechanisms that limit cAMP signaling, the expression of endogenous PKI-inhibitor protein (PKIA), which limits signaling of downstream of PKA was increased by 93% (p<0.001) in TG^AC8^ vs WT (**Table S.3**). Protein phosphatase 1 (PP1) was increased by 50% (**Fig. S.11 A**). The Dopamine-DARPP-32 feedback on cAMP signaling pathway was enriched and also activated in TG^AC8^ vs WT (**Fig. 12**), the LV and plasma levels of dopamine were increased, and DARPP-32 protein was increased in WB by 269% (**Fig. S.11 A**).

Thus, mechanisms that limit signaling downstream of AC-PKA signaling (βAR desensitization, increased PDEs, PKI inhibitor protein, and phosphoprotein phosphatases, and increased DARPP-32, cAMP (dopamine- and cAMP-regulated phosphoprotein)) are crucial components of the cardio-protection circuit that emerge in response to chronic and marked increases in AC and PKA activities (**Fig. 4 C, F**).

#### Upregulation of stress response pathways

Beyond the adaptations to limit the degree to which cAMP/PKA signaling is activated, numerous canonical stress-response signaling circuits were concurrently enriched in TG^AC8^ (**Fig. 13**). Multiple parallel cascades of stress response receptor signaling within circuits downstream of AC signaling (**Fig. 13)**, include: RTKs (PI3K/AKT, ERK-MAPK); cytokine receptors (JNK1/2, JAK2/STAT3), pattern recognition receptor RAGE (S100/calgranulins) and T-cell receptor (NF-κB) (**Fig. 13**). The main functional category of genotypic changes identified in PROTEOMAP analysis, **Environmental Information Processing** (**Fig. S.7**), points to an integration of these stress response pathways. The consilience of cardio-protective adaptations that result from integrated activation of signaling cascades that were enriched in TG^AC8^ included: cell survival initiation, protection from apoptosis, proliferation, prevention of cardiac-myocyte hypertrophy, increased protein synthesis and quality control, increased inflammatory and immune responses, facilitation of tissue damage repair and regeneration and increased aerobic energetics (**Fig. 13**).

Pathways downstream of receptor tyrosine kinase (RTK), a classical cognate receptor, amongst others, transmits signals to promote the activation of PI3 kinase and RAS-RAF-MEK1/2-ERK1/2 (**Fig. 13**). Receptor activation by adrenergic agonists and growth factors (such as periostin, increased by 39% in omics and validated by antibody array (**Table S.5, Fig. S.14**) bind to β-AR and RTK, respectively. Ras-p21 functions as a molecular link for membrane GPCR and RTK to transduce signals from these receptors to downstream signaling machinery (**Fig. 13**, ERK arm). Ras-Raf-MEK-ERK pathway plays a key role in cardioprotection in the context of ischemia-reperfusion injury and oxidative stress.^68^ Downstream of Ras-p21, c-Raf is subsequently induced, which, in turn, sequentially activates MEK1/2 and ERK1/2 to achieve their downstream effects (**Fig. 13**, ERK arm, **Fig. 4 B, C**). MEK/ERK1/2 activation promotes tissue repair that is essential for repair of damaged cells or cell regeneration (see below) in response to cardiac stress in vivo (**Fig. 13**, ERK arm), especially those observed in genetically engineered animal models.^69^

Ras-p21 signaling also exerts protective influences on cell reparative and regenerative capacity, presumably via activation of ERK-independent downstream pathways of PI3K/AKT signaling^35, 70^. Increased expression of the catalytic subunit PIK3CA in TG^AC8^ by 15 % in WB (**Fig. S.8 A**) may be linked to the activation of the downstream targets, including PDK1 phosphorylation on serine 241, subsequent phosphorylation of AKT2 (i.e., threonine 308 and serine 473) and of the transcription factor, Foxo1, at serine 256 (**Fig. S.11 B, Fig. 13**, PI3K arm, bottom panel). Activation of AKT inhibits hypertrophic signaling in adult hearts in vivo, ^71 72 73 18^ while inducing protein synthesis and activating protein quality control signaling ^74^ (**Fig. 13**, PI3K arm). This anti-hypertrophic effect of AKT in TG^AC8^ may be linked to the lack of increased myocyte size and absence of increased LV mass in TG^AC8^ (**Fig.s. 1J, 2B, C, D**), (see below), even in the presence of elevated α-skeletal-actin and MYH6 (**Fig. 4 G, Fig. 13**, bottom panel).

The tumor suppressor PTEN is a lipid phosphatase that regulates cell growth, survival and migration by catalyzing the dephosphorylation of the phospholipid phosphatidylinositol (3,4,5)-trisphosphate PtdIns (3,4,5)P3 or PIP3, an integral second messenger molecule for the PI3K-AKT signaling, thus antagonizing this pathway. PTEN protein abundance was increased by approximately 28% (p<0.01) in the TG^AC8^ LV compared to WT by WB (**Fig. S.8 E**), in concordance with proteomics data, which showed a 22% increase in PTEN (p=0.053) in the TG^AC8^ LV. Phosphorylation of sites within the PTEN C-terminal domain, including S380, T382 and T383, have been demonstrated to be involved in the regulation of stability and activity, with a loss in phosphorylation leading to membrane recruitment and greater activity followed by rapid proteosome-mediated degradation.^75–77^ Phosphorylation of Ser380 residues was significantly lower in TG^AC8^ vs WT (−35%, p<0.05) as was the ratio of p-PTEN Ser380/Total PTEN (−50.15%, p<0.005) (**Fig. S.8**), suggesting that PTEN activity is increased in the TG^AC8^ vs WT. A 2-fold higher expression of PIP3 in TG^AC8^ vs WT, determined by ELISA, along with increases in PI3K catalytic subunit abundance and AKT activity, strongly suggest that PTEN does not exert **a net** negative regulatory effect in TG^AC8^ on this pathway. Rather, an increase in PTEN in TG^AC8^ may be necessary to maintain the available pool of PIP2 that is required for phospholipid metabolism, and other signaling pathways, such as e.g. PLC/IP3/DAG signaling (important in Ca^2+^ regulation), or in functional processes like e.g. Endocytosis and Actin cytoskeleton remodeling,^78^ that are enriched and activated in TG^AC8^ vs WT (**Fig. 10 C and Table S.6 A,B**).

Integral components of the PI3K-AKT and RAS-RAF-MEK1/2-ERK1/2 survival-associated pathways converge to suppress apoptosis^79–81^ (**Fig. 13**, PI3K and ERK arms). Caspase-3 levels were increased (**Fig. S.11 B**). The barely detectable or low protein expression of hallmarks of apoptosis in TG^AC8^ LV, including *cleaved* caspase-3 and its downstream *cleaved* PARPs, suggest that apoptosis suppression signaling mechanisms become activated in the TG^AC8^ LV (**Fig. S.11 B and Fig. 13**, PI3K and ERK arms top and bottom panels). That full-length caspase-3 expression was increased in TG^AC8^ vs WT, but not cleaved (**Fig. S.11 B and Fig. 12**),^82^ provides strong evidence of activation anti-apoptotic functions of caspase-3 activation in the TG^AC8^ LV. Further, there was no indication of increased apoptotic nuclei in TG^AC8^ LV tissue (**Fig. S.13 A**).

### Proteostasis

Many upregulated transcripts and proteins in TG^AC8^ vs WT (**Fig. S.7, Tables S3, S.5**), including the serine/threonine-protein kinases, MSK1 and MNK1, might signal within cardio-protective pathways (**Fig.s. 5 F, G; 13**). Specifically, MSK1 is involved in regulation of the transcription factors CREB1, ATF1, RELA and STAT3. MNK1 is involved in the initiation of protein translation by interacting with eIF4G and phosphorylating eIF4E.

The Small Proline Rich Protein 1 **Sprr1a,** which is linked to induction of protein synthesis (**Fig. 13**, bottom panel), was the most abundantly overexpressed transcript **and** protein in the TG^AC8^ **LV ( Table S.7 A).** Immunolabeling of Sprr1a in single LV myocytes isolated from TG^AC8^ is illustrated in **Fig. S.15**. Sprr1a is associated with inflammation, cellular stress, and repair^83, 84^ and has been linked to protection against cardiomyocyte death in the setting of ischemia– reperfusion injury.^85, 86^ Sprr1a stimulates the expression of Rtn4 (**Fig. S.15**), a member of the reticulon encoding gene family associated with endoplasmic reticulum that is involved in neuroendocrine secretion or in membrane trafficking in neuroendocrine cells.^87, 88^ Rtn4, which was also overexpressed in TG^AC8^ vs WT (transcripts increased by 50% and proteins increased by 30%, **Fig.** S.**15**, **Table S.7 A)**.

Derlin1, a protein that was also markedly overexpressed in TG^AC8^ vs WT (by 132%, **Table S.5**) participates in the ER associated protein degradation by recognizing and selecting misfolded or unfolded proteins and translocating these from the ER lumen to the cytosol for proteasomal degradation. Calnexin, and calreticulin both members of the CALR/CN cycle, which ensures the quality and folding of newly synthesized molecules within the endoplasmic reticulum, were also both increased by WB in TG^AC8^ (**Fig. S.11 D**). Numerous other molecules involved in unfolded protein response signaling significantly differed by genotype in omics analyses: transcripts included ATF4, ATF6, Caspase 3, CALR, HSP90 (**Table S.3**); proteins included: CASP3, CASP6, HSP90α (**Table S.5**).

### Autophagy

Forced changes in energy metabolism, downstream of markedly increased levels of AC/cAMP/PKA signaling in TG^AC8^, and resultant adaptations to this stress (**Fig. 13)**, may produce excess wear-and-tear damage in cellular components, requiring an adaptively higher autophagy/mitophagy machinery. The preserved level of mitochondrial fitness (**Fig. 6 I, J; Fig. 7 M, N**) in TG^AC8^, in the context of marked chronic AC-driven cellular stress, likely reflects one such autophagic adaptation. Protein levels of ATG13, ATG4B and PARKIN, members of the autophagy machinery were upregulated in TG^AC8^ vs WT (**Fig. 6 E-G**). Specifically, the cysteine protease ATG4B processes pro-MAP1LC3 and pro-GABARAP in the early stages of autophagy and endocytosis pathways, before their lipidation (addition of the phosphatidylethanolamine) by ATG7 and ATG3. Being also involved in MAP1LC3/GABARAP delipidation, it plays a key role in their turnover. The autophagy-related protein ATG13 is a factor required for autophagosome formation and part of the initiation complex with ULK1 and RB1CC1. It is also involved in mitophagy, together with the E3 ubiquitin-protein ligase PARKIN, which acts downstream of PINK1. Upregulation of these proteins protects against mitochondrial dysfunction during cellular stress by coordinating mitochondrial quality control mechanisms that remove and replace dysfunctional mitochondrial components.

### Inflammation/Immune Signaling

The myocardium is intimately connected with the immune system and activation of the immune system has been shown to have both protective and maladaptive effects on the heart. ^89^ This inherent duality of the immune system has spurred a quest for tools to harvest the protective effect of the innate and adaptive immune response without experiencing their detrimental effects. ^90^ On this background, it is remarkable that the most upregulated pathway in the TG^AC8^ hearts was “leukocyte extravasation signaling” (**Fig. 10 C**) and that the TG^AC8^ was characterized by up regulation of several inflammatory molecules and pathways, both in the “omics analysis” and in western blot based or RT-qPCR based validation analyses. TG^AC8^ hearts in fact showed upregulation of *IL-6*, *IL-10*, ICAM1, and CCL12 (**validated in Fig. S.12 A**). Furthermore, they showed upregulation of MAGUK and MALT1 of the CBM complex together with its downstream NF-κB signaling (NF-κB coactivators TRAF2/6 and ATF3 (**Fig. 13**, NF-κB arm, **Fig. S.7**, **Table S3, S.5 and Fig. S.12 A**)). The TG^AC8^ heart was also characterized by phosphorylation of JNK1/2 on threonine 183 and tyrosine 185 (**Fig. 13**, JNK arm, and **Fig. S.8**) that paralleled gene expression changes consistent with activation of the JNK1/2 signaling pathway in TG^AC8^ vs WT (**Tables S.3, S.5, S.6 B**).

The fact that we detected upregulation of T cell receptor signaling and B cell receptor signaling (**Table S.6 A, B**) together with the fact that we detected upregulation of the CBM complex (central in B and T cell activation) suggests that the AC8 heart might be characterized by recruitment of T and B cells to the myocardium. ^91^ Specific subclasses of lymphocytes have been shown to have protective effect on the heart. ^89^ JNK1/2 activation is centrally involved in inflammatory signaling^92^ but also linked to inhibition of cardiac hypertrophy^93^ and promotion of reparative and regenerative capacities. ^94^ Taken together, these observations raise the intriguing possibility that the AC8 overexpressing heart might manage to cope with continuous stress also through the activation of a cardio-protective immune responses.

### TG^AC8^ LV protection circuits resemble some adaptive mechanisms that accompany disease states

We next performed IPA analyses in order to ascertain the extent to which cardiac protection circuits identified in the TG^AC8^ LV (**Fig. 13**) might be utilized as adaptations to the stress of various disease states. The top disease categories identified in both the TG^AC8^ transcriptome and proteome were **Organismal Injury and Abnormalities and Cancer (Fig. S.16),** the latter being subsumed within the former category. This is not surprising, because the sequalae of marked chronically increased AC/cAMP/PKA signaling illustrated in TG^AC8^ LV protection circuitry **(Fig. 13),** are very similar to the “**Hallmarks of Cancer**”,^15, 16^ and included: protection from apoptosis, survival, aerobic glycolysis, increased proliferation, enhanced protein synthesis and quality control, increased inflammatory and immune response, enhanced tissue damage repair and regeneration. Thus, the TG^AC8^ LV utilizes a consilient pattern of adaptive mechanisms that emerge in many cancer cell types for self-protection.^15, 16^ This protection circuitry allows the adult TG^AC8^ heart to cope (for at least up to a year),^10^ with numerous, marked cell-tissue stressors driven by the markedly augmented AC-cAMP-PKA-Ca^2+^ signaling. “Limited versions” of this cardio-protective profile that emerges within TG^AC8^ LV have also been previously discovered to be central to cardiac ischemic pre-conditioning^95^ and exercise endurance cardiac conditioning.^1, 96^

### Opportunities for Future Scientific Inquiry Afforded by the Present Results

By design our systems approach to elucidate some general foundations of the consilient adaptation profile that protects the chronically high performing TG^AC8^ heart, represents only a “first port of call”, defining some general features of the TG^AC8^ cardiac performance/protection profile. Because many of the perspectives depicted in the scheme in **Fig. 13** were derived from bioinformatic analyses of cell lysates, it is not implied that all of these circuits are present or become activated in all cell types that reside in the LV myocardium. It is apparent, however, from our results, consilient shifts in enrichment or activation of numerous stress response pathways accrue to allow for the chronic high-level performance of the TG^AC8^ LV. Similarly, a consilience of shifts in numerous stress response pathways may be of biological significance in pathologic states, for example, e.g. consilient adaptations may be allowed for proper healing in disease state, e.g. myocardial infarction or chronic heart failure.

A most important aspect of our results, is that they identify numerous hypotheses that can be tested in future, more in depth studies aimed at: (1) defining the precise cardiac cell types in which protection circuitry is activated in the chronically over-worked TG^AC8^ heart; (2) providing deeper definitions of the multitude of signaling pathways that differed significantly in enrichment and activation status among cell types in TG^AC8^ vs WT (**Fig.s. 10-13, Table S.6 A, B**). This will lead to identification of novel associations among the concurrently activated adaptive mechanisms within and among cell types in TG^AC8^ LV that are not intuitively linked to cardiac protection in the context of our current state of knowledge. For example, elucidation of specific details within the leukocyte extravasation pathway, the top activated IPA canonical pathway in TG^AC8^, and of other highly activated canonical signaling pathways in TG^AC8^ vs WT (**Fig. 10 C** and **Table S.6 A, B)**, that are not usually addressed in the field of cardiac research. This will permit detection of crosstalk between cardiac myocytes and other cell types such as immune cells and others that is likely to be critical to the cardiac protective and performance-enhancing circuitry that is harbored within TG^AC8^ heart (**Fig. 13**). (3) Identification post-translational protein modifications e.g. via delineations of changes in the TG^AC8^ LV phosphoproteome, ubiquitome, acetylome, and 14-3-3 interactome; (4) identifying specific types of intersticial cells that proliferate within the LV myocardium that was predicted by our omics results, and validated by EdU and BrdU labeling of the young adult TG^AC8^ heart between 2 and 3 mo (**Fig. 3**). (5) precisely defining shifts in metabolism within the cell types that comprise the TG^AC8^ LV myocardium via metabolomic analyses, including fluxomics.^97^ It will be also important that future metabolomics studies elucidate post-translational modifications (e.g. phosphorylation, acetylation, ubiquitination and 14-3-3 binding) of specific metabolic enzymes of the TG^AC8^ LV, and how these modifications affect their enzymatic activity.

Finally, chronic consilient utilization of numerous adaptive pathways, however, may present a double-edged sword, compensating for chronic stress in the “short run”, but becoming maladaptive when utilized over prolonged periods of time. In fact, TG^AC8^ mice have a reduced (about 30%),^10^ a median life span, and later in life (at about 12 mo) marked LV fibrosis and dilated cardiomyopathy occur. Future studies are required to precisely discover how the consilience of cardio-protective adaptations harbored within the TG^AC8^ heart at 3 mo of age that were defined in present study fails over a protracted time-course are extremely important. It is noteworthy in this regard, when our omics data were specifically filtered for “**cardiovascular system”** and “**cardiovascular disease”** in bioinformatic analyses, features common to variety of cardiomyopathies were predicted in the “super-performing” TG^AC8^ LV (**Table S.11 A, B**). The basis of this prediction may stem from the fact that omics “knowledge” generated by findings in prior cross-sectional studies of experimental heart failure, it seems to us may have often been misinterpreted to be **causal** factors of the heart failure, rather than adaptations that react to ameliorate the heart failure, due to a lack of longitudinal prospective. Thus studies, conducted *longitudinally* over the TG^AC8^ life course, to generate testable hypothesis about which of these correlated findings are *cause-and-effect* vs those that are simply *associated* are required to dissect out which factors that enable the “super performing heart” of the TG^AC8^ during young adulthood, gradually fail over a prolonged period of time, resulting not only in accelerated aging, but also in and frank, severe, dilated cardiomyopathy.^10^

## Acknowledgments

We thank the NIDA Confocal and Electron Microscopy Core for access to their equipment, and specifically Zhang Shiliang (Core manager) and Marisela Morales (Director). We wish to thank Loretta Lakatta for her superb editorial assistance.

## Supplements

### Supplemental Tables

**Table S.1. List of Echo parameters recorded in TG^AC8^ heart vs WT.**

**Table S.2. A Listing of all 11810 identified transcripts.**

**Table S.3. A listing of 2323 transcripts that significantly differed in expression by genotype.**

**Table S.4. A listing of all 6834 identified proteins.**

**Table S.5 A listing of 2184 proteins that significantly differed in expression by genotype.**

**Table S.6. A listing of canonical pathways in IPA analysis of the total number of (A) transcripts and (B) proteins that were differentially enriched or activated in TG^AC8^ vs WT. (C) A listing of molecules with indication of involvement in number of pathways.**

**Table S.7. (A) A listing of 544 molecules of which both transcripts and proteins differed by genotype. (B) A listing of canonical pathways that were differentially enriched by genotype in IPA analysis of 544 transcripts and proteins.**

**Table S.8. A Complete list of downstream effects of enriched canonical signaling pathways in TG^AC8^, depicted on Fig. 11.**

**Table S.9. RT-qPCR analysis of selected transcripts related to G-protein Coupled Receptor Signaling, that differed by genotype in RNASEQ.**

**Table S.10 Transcripts and proteins involved in Cell Cycle/Cell Proliferation, and Growth and Developmental Circuits that differed between TG^AC8^ vs WT.**

**Table S.11 IPA representation of top cardiovascular disease-related functions within the LV (A) transcriptome and (B) proteome of TG^AC8^ and WT.**

**Table S.12 Primers used in RT-qPCR analyses**

**Table S.13 Antibodies used in WB analyses and antibody arrays**

## Supplemental Methods

### Echocardiography

Mice underwent echocardiographic (Echo) examination (40-MHz transducer; Visual Sonics 3100; Fuji Film Inc, Seattle, WA) under light anesthesia with isoflurane (2% in oxygen) via nosecone, temperature was maintained at 37°C using a heating pad. Mice were placed in the supine position; skin hair in the chest area was shaved. Standard ECG electrodes were placed on the limbs and ECG Lead II was recorded simultaneously with acquisition of echo images. Each Echo examination was completed within 10 min. Parasternal long-axis views of the LV were obtained and recorded to ensure that mitral and aortic valves and the LV apex were visualized. From the parasternal long-axis view of the LV, M-mode tracings of LV were obtained at mid-papillary muscle level. M-mode tracing of Left atrium and basal aorta were recorded at aortic valve level and Left atrial dimension (LAD) and Aortic lumen dimension (AoD) were measured. Mitral valve blood flow velocity (E and A waves) was recorded at the tip of the mitral valves at an angle of 450. Parasternal short-axis views of the LV were recorded at the mid-papillary muscle level. Endocardial area tracings, using the leading-edge method, were performed in the 2D mode (short-axis and long-axis views) from digital images captured on a cine loop to calculate the end-diastolic and end-systolic LV areas. LV End-diastolic volume (EDV) and end-systolic volume (ESV) were calculated by a Hemisphere Cylinder Model method. Stroke volume (SV) was calculated as SV = EDV-ESV. Cardiac output (CO) was calculated as CO = SV*HR. Ejection Fraction (EF) was derived as EF = 100 * (EDV -ESV) / EDV. Cardiac Index (CI) was calculated as CI = CO / BW. LV Posterior Wall (PW) and Inter Ventricular Septal thicknesses (IVS) were measured from the LV M-mode tracing LV. LV mass (LVM) was calculated from EDV, IVS and PW. LV early (E) and late (A) diastolic filling rates, and early/late ratio (E/A) were calculated from mitral valve blood flow velocities.

All measurements were made by a single observer who was blinded to the identity of the tracings. All measurements were reported an average of five consecutive cardiac cycles covering at least one respiration cycle (100 times/min in average). The reproducibility of measurements was assessed by repeated measurement a week apart in randomly selected images; the repeated-measure variability was less than 5%.

Echocardiography data are expressed as mean ± SEM. Differences between two groups were assessed by a t-test. Statistical significance was assumed at p<0.05.

### Heart and cardiac tissue isolation

Mice were injected (intraperitoneally) with heparin and acutely anesthetized with pentobarbital-based euthanasia solution. The heart then was quickly removed and placed into cold PBS solution. The left ventricle free wall, without the septum, was identified anatomically, under a dissecting microscope, and pieces of tissue 2×3mm were dissected and snap frozen in liquid nitrogen.

### LV Histology

The LV free wall, (excluding the septum) was cut into base, mid-portion, and apex segments, fixed with formalin, embedded in paraffin, and sectioned (5 μm in thickness). Sections were stained with hematoxylin and eosin, silver (Reticulum stain kit, American MasterTech Scientific, Inc, Lodi, CA) and Masson’s trichrome (American MasterTech Scientific, Inc, Lodi, CA) as previously reported4. Myocyte cross-sectional area was measured from images captured from silver-stained 5-μm-thick sections of the LV mid portion sections as described5. Suitable cross sections were defined as having nearly circular-to-oval myocytes at the nuclear level. Outlines of ∼25 myocytes were traced in each section. Morphometric Analyses were performed using the computerized imaging program MetaMorph (MetaMorph Imaging System, Universal Imaging Corp) using light microscopy.

### EdU labeling for detection and Imaging of cardiac cell DNA synthesis

Mice were administered 0.35mg/L 5-ethynyl-2’-deoxyuridine (EdU) for 28 days via drinking water, changed every third day. Mice were then administered pentobarbital IP, heart removed and placed in PBS. The aorta was cannulated, and the heart perfused with PBS for 5 min followed by perfusion at ∼100mmHg with 4% paraformaldehyde for approximately 10 min or until flow rate was greatly reduced. Hearts were stored in fresh 4% formaldehyde for 24hr at 4 degrees. Hearts were washed with phosphate buffer and imbedded in 4% low melting point agarose. Sequential transverse sections from the heart of WT and TG^AC8^ transgenic mice were sectioned on the Leica Vibratome VT1000s from 200u-300uM. Sections were permeabilized with 0.2% triton, glycine, and 2% DMSO for 3 days. EDU labeling (Click Chemistry Tools), and primary antibodies Vimentin 1:500 (Synaptic Systems), Actinin (Sigma) 1:500, WGA 1:500 (Vector Labs), 4′,6-diamidino-2-phenylindole DAPI 1:300 (Sigma). Five microscopic fields, per mouse, in the left ventricle were visualized in cardiomyocytes via fluorescent imaging (Zeiss LSM 980) at 400 x magnification, and the number of cardiac nuclei staining positively for EdU was counted in each.

### BRDU labeling

To monitor cardiomyocyte S-phase activity, TG^AC8^ mice were crossed with MHC-nLAC mice (which express a nuclear-localized β-galactosidase reporter under the transcriptional regulation of the mouse α-cardiac MHC promoter; these mice are useful to identify cardiomyocyte nuclei in histologic sections) ^1^. The resulting TG^AC8^, MHC-nLAC double-transgenic mice and MHC-nLAC single-transgenic mice were identified and sequestered. At 28-to-30 days of age, the mice were administered BrdU via drinking water (0.5 mg/ml, changed every 2nd day) for a total of 12 days. Hearts were then harvested, fixed (1% paraformaldehyde, 50 mM cacodylate, 0.665% NaCl, pH 7.3) for 24 hours at 4°C, cryopreserved (30% sucrose) for 24 hours at 4°C, embedded and cryosectioned at 10 microns using standard methods.^2^ Sections were subjected to antigen retrieval (10 mM trisodium citrate, 0.05% Tween20, pH 6) for 30 min at 100°C, and non-specific signal was then blocked using MOM blocking reagent (Vector Labs, Burlingame California) following manufacture’s recommendations. Sections were then processed for β-galactosidase (#A-11132, Invitrogen Life Sciences, Grand Island New York) and BrdU (#11296736001, Roche, Indianapolis Indiana) immune reactivity; signal was developed using Alexa 555 goat anti-rabbit for β-galactosidase and Alexa 488 goat anti-mouse for BrdU (secondary antibodies were #A21429 and #A1100, respectively, Invitrogen). Sections were counterstained with Hoechst 33342 (Sigma-Aldrich, St. Louis Missouri, blue signal) and cover slipped. After processing, the sections were imaged sequentially for the red, green and blue signals using Surveyor software (version 9.0.4.5, Digital Imaging Systems Ltd., Buckinghamshire, UK) interfaced with a Leica DM5500 (Leica AG, Wetzlar, Germany) microscope. The percentage of S-phase cardiomyocyte nuclei (as evidenced by the overlay of red βGAL signal and green BrdU immune reactivity) was then quantitated.

### Electron Microscopy

Mice left ventricles were dissected and processed for transmission electron microscopy visualization. Fixation for electron microscopy was performed using 2.5% glutaraldehyde in 0.1 M sodium cacodylate buffer, pH 7-7.4. Samples were post fixed in 1% osmium tetroxide for 1 h at 4°C in the same buffer, dehydrated and then embedded in Embed 812 resin (Electron Microscopy Sciences, Hatfield, PA) through a series of resin resin-propylene oxide gradients to pure resin. Blocks were formed in fresh resin contained in silicon molds, and the resin was polymerized for 48-72 h at 65°C. Blocks were trimmed and sectioned in an EM UC7 ultramicrotome (Leica Microsystems, Buffalo Grove, IL) to obtain both semi-thick (0.5-1 µm width) and ultrathin (40-60 nm width) sections. Semi-thick sections were mounted on glass slides and stained with 1% toluidine blue in a 1% borax aqueous solution for 2 min. Micrographs were obtained using a Leica AXIO Imager light microscope with a Axiocam 512 color camera (Carl Zeiss, White Plains, NY). Ultrathin sections were stained with uranyl acetate and lead citrate, and then imaged on a FEI Tecnai G^2^ 12 Transmission Electron Microscope (TEM) with a Gatan OneView 16 Megapixel Camera.

### Electron microscopy image analysis

Micrographs at ×1,200 and x2,900 magnification were obtained from randomly selected areas of cardiomyocytes cytoplasm for illustration and quantitative analysis purposes of mitochondrial population and lipid droplets quantification. We determined two stereological parameters: (a) Na, which is the numerical profile density (number of figures of interest / μm2 of cell fraction), and (b) volume density of figures of interest (Vv; i.e., the volume fraction of cardiomyocyte cytoplasm occupied by figures of interest). Volume density was obtained following a point analysis using a simple square lattice test system.^3^ Stereological measurements were performed using ImageJ software (NIH). Mitochondria were identified as electron dense double membrane organelles vesicles with identifiable cristae. Damaged mitochondria presented swollen and disrupted electron-“lighter” cristae. Lipid droplets were denoted as electron-light, not-limited by any membrane vesicles with very clear and homogeneous content. For stereological analysis, only micrographs depicting longitudinal sections of cardiomyocytes with visible sarcomeres were utilized, and from each of these ten pictures were taken from four to six cells/fibers. From each picture, mitochondria were counted and measured to determine the number and area, doing the average of these metrics in each picture, and finally the average for each cell. With this procedure, a total of ∼500 mitochondria per animal were counted/measured.

### Adenylyl Cyclase Activity in Cell Membranes of LV Tissue

Pieces of left ventricular (LV) tissues from wild type and AC8-TG mice were frozen in liquid nitrogen, homogenized with Bel-Art™ SP Scienceware™ liquid nitrogen-cooled Mini Mortar and stored at −80°C till use in the AC assay. On the day of the assay, 1 ml of ice-cold Lysis buffer (LB) was added to each sample (LB composition: 10 mM Tris, pH 7.6, 0.5 mM DTT, 1 mM EGTA, 0.2 mM IBMX and 0.33% PIC). Samples were sonicated on ice (3 x 15 sec at setting 2, with 15 sec of rest between bursts). 1800 µl of lysates (combined from two mice) were used for membrane isolation. To each such combined sample a Sample Separation Buffer (SSB, composition: 10 mM Tris, pH 7.6, 0.5 mM DTT, 1 mM EGTA, and 0.01% PIC) was added to a total volume of 11 ml. These samples were centrifuged for 10 min at 1,000xg to remove big clumps. Supernatants were further centrifuged for 30 min at 48,254xg at 40C in the Ultra-Clear 14×89 mm ultracentrifuge tubes filled almost to the top with SSB. Membrane proteins precipitated on the bottom of the tubes were washed three times with SSB via ultracentrifugation in the same conditions. At the end of the last wash the pure pellets were resuspended in 300 µl of the Sample Reaction Buffer (SRB, composition: 70 mM Tris, pH 7.6, 0.5 mM DTT, 1 mM EGTA, 5 mM MgCl^2^, 0.2 mM IBMX, and 0.33% PIC) via sonication (on ice, 3 sec x 10 bursts on setting 2, rest between bursts 15 sec). Protein content in the membrane preparations was quantified using a Reducing Agent Compatible Pierce® Microplate BCA Protein Assay Kit # 23252.

Purified LV membranes were further diluted with SRB to a 0.2 µg/µl protein concentration and used in the AC reaction. For the AC activity detection reaction, Stock AC reaction media (SACRM) was prepared: 70 mM Tris (pH 7.6), 0.5 mM DTT, 1 mM EGTA, 5 mM MgCl^2^, 0.2 mM IBMX, 4 mM ATP, 20 mM Creatine Phosphate, and 240 U/ml Creatine phosphokinase. The end AC reaction composition was: 70 mM Tris (pH 7.6), 0.5 mM DTT, 1 mM EGTA, 5 mM MgCl2, 0.2 mM IBMX, 0.25% PIC, 1 mM ATP, 5 mM Creatine Phosphate, 60 U/ml Creatine phosphokinase, 0.2% DMSO, and 0.15 µg/µl membrane proteins.

To proceed the reaction, to the tube, preheated to 35°C with 25 µl of the SACRM, 75 µl of the membrane sample was added. The AC reaction lasted for 5 min at 35°C at 400 RPM, and was stopped by immersion of the tube into a 100°C steel shot for 5 min. Then tube was cooled down, centrifuged for 5 min at 40°C at 15,000xg, and the supernatant was used for cAMP measurement. 0 time samples were prepared the following way: first to the tubes 75 µl of membrane proteins were added, then proteins were denatured for 5 min at 100°C and cooled down, after that 25 µl of the SACRM was added to the tubes, and they were immediately immersed into 100°C steel shot for 5 min. Then these 0 time samples were processed exactly the same way as other samples.

The cAMP concentration was quantified via a LANCE kit protocol (Lance cAMP384 kit 500 points, Perkin Elmer, AD0262) in 96-well OptiPlates (Perkin Elmer). 15 µl of the sample was used in the LANCE assay in a total volume of 40 µl (including 5 µl of 2x cAMP antibodies and 20 µl of the Detection Mix). 2x cAMP antibodies stock preparation: 40 µl of Ab stock, 107 µl of 7.5% BSA, 1853 µl of Detection buffer. Preliminary experiments demonstrated that cAMP LANCE standard curves depend a lot on the buffer in which they were prepared; because of that cAMP standards were prepared in the same buffers that were used for samples (75% SRB, 25% SACRM), they were heated, cooled down and centrifuged. Right before fluorescence detection, plates were centrifuged for 2 min at 1000xg to remove bubbles from the wells and to increase accuracy of the measurements. All measurements were done in triplicate and the average of the 3 taken as the cAMP value of that sample. Three pairs of WT and three pairs of TG^AC8^ mice LV samples were used in this experiment. Statistic – values as mean ± St Error.

### Immunostaining of isolated intact mice ventricular myocytes for ADCY8, SPRR1A and ACACB detection

Immunolabeling was performed in freshly isolated LV mouse cells. Cells were plated on laminin coated MatTek dishes for 1h, 4% paraformaldehyde for 10 minutes, washed 3 times with PBS, and then permeabilized with 0.2 % Triton X-100 in PBS for 10 minutes at room temperature. The plates were washed two more times with PBS and then incubated with 10% goat serum for 1 hour to minimize nonspecific staining. Afterwards, samples were incubated at 4 °C overnight with primary antibodies against SPRR1 (ab125374 ), ADCY8 ( bs-3925R) and ACACB(sc-390344). Cells were then washed 3 times with PBS and incubated with fluorescence-conjugated secondary antibodies (1:1000) (Sigma, USA) for 45 min at 37 °C. Cell nuclei were labeled with DAPI (Sigma, USA). Cells were visualized using a LSM 710 laser-scanning confocal microscope (Carl Zeiss) and images were captured using the Carl Zeiss Zen software. Quantitative fluorescence image analysis was performed with ImageJ software, according to the following protocol: http://theolb.readthedocs.io/en/latest/imaging/measuring-cell-fluorescence-using-imagej.html.

Images of stained cells were transferred and analyzed with Image J software to calculate the basic characteristics of each image, including Area, Mean Gray Value and Integrated Density. To calculate the corrected total cell fluorescence (CTCF). Small areas of positively stained fluorescent cells were selected using a free hand selection tool. A background reading was created by selecting a negatively stained rectangular section near the analyzed cell. Total fluorescence per cell was calculated in Excel with the following formula: CTCF = Integrated Density – (Area of selected cell X Mean fluorescence of background readings)

### Immunostaining of isolated intact mice ventricular myocytes for RyR2 detection

Immunostaining was performed as previously described. Specifically, freshly isolated mice ventricular myocytes from TG^AC8^ and WT control mice were fixed with 4% paraformaldehyde, permeabilized with 1% Triton and incubated with blocking solution (1×PBS containing 2% IgG-free BSA+ 5% goat serum+0.02% NaN3+0.2% Triton). Then, the cells were incubated with primary antibody anti-total RyR (Santa Cruse, R128, 1:500) overnight. After several wash, secondary Atto 647N-conjugated anti-mouse IgG (Sigma-Aldrich, 1:500) antibody was used, and only secondary antibody was applied to negative controls, which displayed negligible fluorescence. Confocal images of middle section were obtained via Zeiss LSM 510 (Carl Zeiss Inc., Germany) using 633 nm laser to excite the fluorophore Atto 647N. The images were analyzed using ImageJ software (1.8V, Wayne Rasband, National Institutes of Health). Please note that the RyR2 immunolabeling in TG^AC8^ mice ventricular myocytes was saturated if using the same sampling setting as WT controls. So, the sampling condition was adjusted when sampling ventricular myocytes from TG^AC8^ mice, and the density was converted to the same setting as WT control for comparison. ^4^

### Protein synthesis

Protein synthesis was assessed by SUnSET-Western Blot as previously described 6. Briefly, the puromycin solution was prepared in PBS, sterilized by filtration, and a volume of 200µl was injected in mice intraperitoneally, to achieve a final concentration of 0.04µmol/g of body mass. After 30 minutes, mice were sacrificed, the LV was harvested and snap frozen in liquid nitrogen. Protein extraction was performed using Precellys, quantified with BCA assay 25µg of total protein were separated by SDS-PAGE; proteins were then transferred onto PVDF membrane and incubated overnight in the anti-puromycin primary antibody (MABE343, Sigma-Aldrich, St. Louis, MO). Visualization of puromycin-labelled bands was obtained using horseradish peroxidase conjugated anti-mouse IgG Fc 2a secondary antibody (Jackson ImmunoResearch Laboratories Inc., West Grove, PA, USA), using Pierce Super Signal ECL substrate kit (Pierce/Thermo Scientific Rockford, IL). Chemiluminescence was captured with the Imager AI600 and densitometry analysis was performed using ImageQuantTL software (both by GE, Boston, MA). Total protein was used as control for protein loading.

Genotypic differences of protein synthesis were tested via an as unpaired t-test.

### Proteosome activity assay

Flash frozen tissue was homogenized in ice-cold cytosolic extraction buffer (50 mM Tris-HCl pH 7.5, 250 mM Sucrose, 5 mM MgCl_2_, 0.5 mM EDTA, and 1 mM DTT). A bicinchoninic acid (BCA) assay (Pierce) was used to determine the protein concentrations. All samples were equally concentrated in proteasome assay buffer (50 mM Tris-HCl pH 7.5, 40 mM KCl, 5 mM MgCl_2_, and 1 mM DTT). Proteasome activity was determined in the presence of 28 µM ATP using the Suc-LLVY-AMC (18 µM, Boston Biochem #S280) fluorogenic substrate with and without proteasome inhibition (MG 132, 1 mM, Sigma). The plate was read at an excitation wavelength of 380 nm and an emission wavelength of 469 nm using a Spectramax M5 (Molecular Devices). Activity was calculated by subtracting the background (proteasome inhibited value) from the reading (proteasome activated value).

### Protein Aggregation Assays

Protein aggregates were measured using Proteostat (Enzo, ENZ-51023) following the manufacturer’s instructions. For this assay left ventricle myocardial lysate (Cell Signaling lysis buffer) was obtained, protein concentration assayed (BCA assay [Pierce]). Ten μg of protein loaded into a 96-well microplate and protein aggregates were analyzed using the Proteostat assay kit (Enzo Life Sciences) following the manufacturer’s instructions. Background readings were subtracted from sample recordings and were normalized to wild-type values.

### Quantibody® Mouse Inflammation and Periostin Arrays

Fresh LV tissue from 3 months old mice was homogenized and lysed in RIPA buffer (Sigma Aldrich) supplemented with protease inhibitor (Roche Inc.) using a Precellys homogenizer with the CKMix Tissue Homogenizing Kit. The supernatant was collected after centrifugation at 10,000×g for 10 min at 4 °C. The protein concentration was determined using the Bicinchoninic Acid (BCA) Assay (Thermo Fisher Scientific). The assay was performed using the Quantibody® Mouse Inflammation Array kit (QAM-INF-1-1, RayBiotech Inc.). The samples (tissue lysates) were diluted 2x using the sample diluent. 100 ul of both samples and standards were loaded on to the glass slide (labelled with the 40 different cytokines and chemokines) and incubated for 2 hours at room temperature. Samples and standards were decanted and from each well and washed with 150 µl of 1X Wash Buffer I at room temperature. The detection antibody cocktail was added to each well and incubated at room temperature for 1 hour. The samples were decantyed and washed with 150 µl of 1X Wash Buffer I at room temperature. A Cy3 equivalent dye-conjugated streptavidin was added to each well and incubated in dark at room temperature for 1 hour. The samples were decanted from each well, washed with 1x wash buffer and dried. The slide was visualised for signals using a laser scanner equipped with a Cy3 wavelength (green channel). The data was extracted, computed in the standard format as provided by the company and analyzed for relative levels of different cytokines in both WT and TG^AC8^ mice. Periostin was assessed in an array-based ELISA system (Growth Factor Quantibody Array, RayBiotech Life, Inc., Peachtree Corners, GA). Briefly, after a blocking step, 100uL of LV-tissue lysates were added on a slide containing a periostin antibody, and incubated 4C overnight. Nonspecific proteins were then washed off, and the arrays incubated with a cocktail of biotinylated detection antibodies, followed by a streptavidin-conjugated fluorophore. Signals were visualized using a fluorescence laser scanner. Relative quantification was calculated using the median, after subtracting the background intensity.

### PKA Activity

Enzymatic activity assays with heart tissue lysates were performed for cAMP-dependent protein kinase (PKA) using the PKA Kinase Activity Assay Kit (Abcam, ab139435), in accordance with the manufacturer’s instructions. The units are expressed in OD/mg protein/min.

### Western Blotting

Snap-frozen left ventricle (LV) tissue from 3 month old mice was homogenized and lysed in ice cold RIPA buffer (Thermo Fisher Scientific: 25 mM Tris-HCl (pH 7.6), 150 mM NaCl, 1% NP-40, 1% sodium deoxycholate, 0.1% SDS) supplemented with a Halt protease inhibitor cocktail (Thermo Fisher Scientific), Halt phosphatase inhibitor cocktail (Thermo Fisher Scientific) and 1 mM phenylmethyl sulfonyl fluoride, using a Precellys homogenizer (Bertin Instruments) with tissue homogenization kit CKMix (Bertin Instruments) at 4 °C. Extracts were then centrifuged at 10,000×g for 10 min at 4 °C and the protein concentration of the soluble fraction determined using the Bicinchoninic Acid (BCA) Assay (Thermo Fisher Scientific). Samples were denatured in Laemmli sample buffer (BioRad Laboratories) containing 355 mM 2-mercaptoethanol at 95oC for 5 minutes, and proteins (10-50 μg/lane) resolved on 4-20% Criterion™ TGX Stain Free™ gels (Bio-Rad Laboratories) by SDS/PAGE. Gels then exposed to UV transillumination for 2.5 minutes to induce crosslinking of Stain Free™ gel trihalo compound with protein tryptophan residues. Proteins were then transferred to low fluorescence polyvinylidene difluoride (LF-PVDF) membranes (BioRad Laboratories) using an electrophoretic transfer cell (Mini Trans-Blot, Bio-Rad). Membrane total protein was visualized using an Amersham Imager 600 (AI600) (GE Healthcare Life Sciences) with UV transillumination to induce and a capture fluorescence signal. Blocked membranes (5% milk/tris-buffered saline with Tween-20, TBST) were incubated with the following primary antibodies: anti-MYH6 (MA5-27820) at 1:2,500 working concentration, anti-ANP (PA5-29559) at 1:1,000, and anti-BNP (PA5-96084) at 1:1,000 from ThermoFisher Scientific; anti-MYH7 (ab173366) at 1:500, anti-α-Sk. Actin (ab179467) at 1:1,000, and anti-SERCA2 ATPase (ab91032) at 1:2000 from Abcam. Primary antibodies were then detected using horseradish peroxidase (HRP) conjugated antibody (Invitrogen) at 1:10,000. Bands were visualized using Pierce SuperSignal™ West Pico Plus ECL substrate kits (Thermo Scientific), the signal captured using an Amersham Imager 600 (AI600) (GE Healthcare Life Sciences) and quantified using ImageQuant TL software (GE Healthcare Life Sciences). Band density was normalized to total protein.

### RT-qPCR

RT-qPCR of LV tissue was performed to determine the transcript abundance of human AC8, genes that mediate neural autonomic input to LV (n=4 WT and 4 TG^AC8^ mice) and to detect genes regulating cytokines level in the heart (n=6 in WT and TG^AC8^). RNA was extracted from left ventricular myocytes (VM) with RNeasy Mini Kit (Qiagen, Valencia, CA) and DNAse on column digestion. The cDNA was prepared using MMLV reverse transcriptase (Promega). RT-qPCR was performed using a QuantStudio 6 Flex Real-Time PCR System (Thermo Fisher Scientific) with a 384-well platform. The reaction was performed with a FastStart Universal SYBR Green Master Kit with Rox (Roche) using the manufacturer’s recommended conditions; the sizes of amplicons were verified. Each well contained 0.5 μl of cDNA solution and 10 μl of reaction mixture. Each sample was quadruplicated and repeated twice using de novo synthesized cDNA sets. Preliminary reactions were performed to determine the efficiency of amplification. RT-qPCR analysis was performed using the ddCt method. Primers were selected with Primer Express 3.0 software (Applied Biosystems). Full list of primers used for amplification provided in **Table S.12.**

### RNASEQ

LV RNA was extracted from 8 of TGAC8 and WT animals. Following a quality control check, RNA was processed with a SMARTer Stranded Total RNA-Seq Kit - Pico Input Mammalian (Takara Bio USA, Inc.). 75 bp single end reads generated 30 to 40 million reads per library. Raw RNA sequencing (RNASeq) reads were aligned and after quality trimming was mapped to the UCSC mm10 mouse reference genome and cDNA of human AC8 and assembled using Tophat v2.0 to generate BAM files for each sample. Cufflinks v.2.1.1 was used to calculate FPKM (Fragments per Kilobase of transcript per Million mapped reads) for each sample. Differential gene expression analysis was performed with a Cuffdiff package (Cufflinks v2.1.1)

### LV Proteome analysis

Four LV samples from WT and TG^AC8^ mouse hearts were snap frozen in liquid nitrogen and stored at −80°C. On average, 2 mg of muscle tissue from each sample was pulverized in liquid nitrogen and mixed with a lysis buffer containing (4% SDS, 1% Triton X-114, 50 mM Tris, 150mM NaCl, protease inhibitor cocktail (Sigma), pH 7.6. Samples were sonicated on ice using a tip sonicator for 1 min with 3 sec pulses and 15 sec rest periods at 40% power. Lysates were centrifuged at +4°C for 15 min at 14000 rpm, aliquoted and stored at −80°C until further processing. Protein concentration was determined using commercially available 2-D quant kit (GE Healthcare Life Sciences). Sample quality was confirmed using NuPAGE® protein gels stained with fluorescent SyproRuby protein stain (Thermo Fisher).

In order to remove detergents and lipids 500 µg of muscle tissue lysate was precipitated using a methanol/chloroform extraction protocol (sample:methanol:chloroform:water – 1:4:1:3).^5^ Proteins were resuspended in 50 ul of concentrated urea buffer (8M Urea, 150 mM NaCl (Sigma)), reduced with 50 mM DTT for 1 hour at 36°C and alkylated with 100 mM iodoacetamide for 1 hour at 36°C in the dark. The concentrated urea/protein mixture was diluted 12 times with 50 mM ammonium bicarbonate buffer, and proteins were digested for 18 hours at 36°C, using trypsin/LysC mixture (Promega) in 1:50 (w/w) enzyme to protein ratio. Protein digests were desalted on 10 x 4.0 mm C18 cartridge (Restek, cat# 917450210) using Agilent 1260 Bio-inert HPLC system with a fraction collector. Purified peptides were speed vacuum dried and stored at −80°C until further processing.

A subset of 8 muscle samples (100 µg) each corresponding to 4 controls and 4 TGAC8 LVs and one averaged reference sample were labeled with 10-plex tandem mass spectrometry tags (TMT) using standard TMT labeling protocol (Thermo Fisher). 200 femtomole of bacterial beta-galactosidase digest (SCIEX) was spiked into each sample prior to TMT labeling to control for labeling efficiency and overall instrument performance. Labeled peptides from 10 different TMT channels were combined into one experiment and fractionated.

### High-pH RPLC fractionation and concatenation strategy

High-pH RPLC fractionation was performed in an Agilent 1260 bio-inert HPLC system using a 3.9 mm X 5 mm XBridge BEH Shield RP18 XP VanGuard cartridge and a 4.6 mm X 250 mm XBridge Peptide BEH C18 column (Waters). The solvent contained 10mM ammonium formate (pH 10) as mobile phase (A), and 10mM ammonium format and 90% ACN (pH 10) as mobile-phase B 9.

TMT labeled peptides prepared from the ventricular muscle tissues were separated using a linear organic gradient from 5% to 50% B over 100 min. Initially, 99 fractions were collected at 1 min intervals. Three individual high-pH fractions were concatenated into 33 master fractions at 33 min intervals between fractions (fraction 1, 34, 67 = master fraction 1, fraction 2, 35, 68 = master fraction 2 and so on). Combined fractions were speed vacuum dried, desalted and stored at −80°C until final LC-MS/MS analysis.

### Capillary nano-LC-MS/MS analyses

Purified peptide fractions were analyzed using UltiMate 3000 Nano LC Systems coupled to the Q Executive HF Orbitrap mass spectrometer (Thermo Scientific, San Jose, CA). Each fraction was separated on a 35 cm capillary column (3µm C18 silica, Hamilton, HxSil cat# 79139) with 200 µm ID on a linear organic gradient at a 500 nl/min flow rate. Gradient applied from 5 to 35 % in 205 min. Mobile phases A and B consisted of 0.1% formic acid in water and 0.1% formic acid in acetonitrile, respectively. Tandem mass spectra were obtained using Q Exactive HF mass spectrometer with a heated capillary temperature +280°C and spray voltage set to 2.5 kV. Full MS1 spectra were acquired from 300 to 1500 m/z at 120000 resolution and 40 ms maximum accumulation time with automatic gain control [AGC] set to 3×10^6^. Dd-MS2 spectra were acquired using a dynamic m/z range with fixed first mass of 100 m/z. MS/MS spectra were resolved to 30000 within of a maximum accumulation time, 120 ms with AGC target set to 2×10^5^. Twelve most abundant ions were selected for fragmentation using 28% normalized high collision energy. A dynamic exclusion time of 45 sec. was used to discriminate against the previously analyzed ions.

### Bioinformatics Analysis of the LV Proteome

Acquired raw data files from Q Exactive HF were converted to mascot generic format (MGF) using MSConvert, an open source software developed by ProteoWizard (http://proteowizard.sourceforge.net). Conversion filters were specified as follows: at MS level 1 with activation:HCD, threshold:count 900 most-intense, zeroSamples:remove Extra 1-, peakPicking;true 1-. Produced MGF files were searched in Mascot against the SWISS-PROT mouse database (02/06/2017) with the following parameters: enzyme trypsin/P, 2 missed cleavages, MS1 tolerance 20 ppm, MS2 tolerance 0.08 Da, quantification TMT10plex. Variable modifications were set to methionine oxidation, carbamidomethylation of cysteines, deamidation at glutamine and asparagine, carbamylation of lysin. Searched mascot data files were processed using commercially available Scaffold Q+ software package (Proteome Software, Inc). Files were merged in to one summary file using MudPIT algorithm, and researched against SWISS-PROT mouse database (02/06/2017), using XTandem search engine for deeper protein coverage with both protein prophet scoring algorithm and protein clustering analysis turned on. Raw reporter ion intensities from unique peptides were extracted into excel file and used in the final analysis.

Minor variations in protein amounts between TMT channels was adjusted by calculating a ratio between signal intensity in each TMT channel (In). Adjusted intensity for each channel (Incorr) was calculated by taking a sum of all intensities in each TMT channel divided by the average (µ) of all calculated sum intensities and multiplied by the initial intensity in each TMT cannel for each peptide: (∑I126+∑I127+∑I128+∑I129+∑I130+∑I131)/6=µ; ∑In/µ*In=Incorr.

Fold change for each unique peptide in each experiment was calculated by dividing Incorr by the median of all intensities in all TMT channels. The fold change between genotypes for each expressed protein was calculated by taking a median of fold change for all unique peptides of a given protein detected by mass spectrometry. Genotype differences were compared via Student’s t-test. A P value < 0.05 was considered to be significant.

### MR spectroscopy High Energy Phosphate

In vivo MRI/MRS experiments were performed on a Bruker spectrometer equipped with a 4.7- T/40-cm Oxford magnet and actively shielded gradients. A one-dimensional 31P chemical shift imaging (1D-CSI) sequence was used to obtain high-energy phosphate data, as previously described. ^6^ The PCr and [β-P] ATP peaks in 31P MR localized spectra were quantified by integration of the peak areas. ^6^

### ROS measurements

ROS measurements were conducted using electron paramagnetic resonance (EPR) spectroscopy as previously described.^7^ Snap-frozen heart tissue (apex, ∼20 mg) was homogenized in phosphate-buffered saline (PBS) containing protease inhibitor cocktail (Roche Applied Science, Indianapolis, IN) and 0.1 mM of the metal chelator, diethylenetriaminepentaacetic acid (DTPA), at pH 7.4. Nonsoluble fractions were removed by centrifugation at 15,000 g for 10 min (4°C). The homogenates were kept on ice and analyzed immediately. Stock solutions of 1-hydroxy-3-methoxycarbonyl-2,2,5,5-tetramethyl-pyrrolidine hydrochloride (CMH; Enzo Life Sciences, Farmingdale, NY) were prepared daily in nitrogen purged 0.9% (w/v) NaCl, 25 g/L Chelex 100 (Bio-Rad) and 0.1 mM DTPA, and kept on ice. The samples were treated with 1 mM CMH at 37 oC for 2 min, transferred to 50-μl glass capillary tubes, and analyzed immediately on a Bruker E-Scan (Billerica, MA) EPR spectrometer at room temperature. Spectrometer settings were as follows: sweep width, 100 G; microwave frequency, 9.75 GHz; modulation amplitude, 1 G; conversion time, 5.12 ms; receiver gain, 2 x 103; number of scans, 16. EPR signal intensities were normalized with respect to the protein concentrations of the tissue homogenates as determined by Pierce BCA protein assay kit (Life Technologies).

### Determination of mPTP-ROS Threshold

Experiments were conducted as described previously,^8^ using a method to quantify the ROS susceptibility for the induction of mPTP in individual mitochondria within cardiac myocytes. ^9^ Briefly, isolated cardiomyocytes were resuspended in HEPES buffer: 137 mM NaCl, 4.9 mM KCl, 1.2 mM MgSO_4_, 1.2 mM NaH_2_PO_4_, 15 mM glucose, 20 mM HEPES, and 1.0 mM CaCl_2_ (pH to 7.3). To assess the susceptibility of the mPTP to induction by ROS, cells were loaded with 100 nM tetramethylrhodamine methyl ester (TMRM; Invitrogen I34361) for at least 2 h at room temperature. Cells were imaged with an LSM-510 inverted confocal microscope, using a Zeiss Plan-Apochromat 63×/1.4 numerical aperture oil immersion objective (Carl Zeiss Inc., Jena, Germany) with the optical slice set to 1 µm. Images were processed by MetaMorph software (Molecular Devices, San Jose, CA). Line scan images at 2 Hz were recorded from ∼22 mitochondria arrayed along individual myofibrils with excitation at 543 nm and collecting emission at >560 nm, and the confocal pinhole was set to obtain spatial resolutions of 0.4 µm in the horizontal plane and 1 µm in the axial dimension. Repetitive laser scanning of this row of mitochondria in a myocyte loaded with TMRM results in incremental, additive exposure of only the laser-exposed area to the photodynamic production of ROS and consequent mPTP induction. The occurrence of mPTP induction is clearly identified by the immediate dissipation of ΔΨ in individual mitochondria and is seen at the point in time where “columns” of the line scan image suddenly lose TMRM fluorescence intensity and become black (Fig.6). The ROS threshold for mPTP induction (tmPTP) was determined as the average time necessary to induce mPTP in the exposed row mitochondria (N=3 in each genotype) (Fig. 6 M,N).

### Determination of autophagolysosome accumulation

Cardiomyocytes were loaded with the autophagy dye from CYTO-ID Autophagy detection kit (ENZO 51031-K200), according the manufacturer protocol (dilution 1:500 in HEPES buffer) and incubated for 30 minutes at room temperature. Then the dye was washed out by HEPES buffer and cell imaged with a confocal microscope (see above) in frame mode using 488 nm excitation and >505 nm emission filter. Images were processed by MetaMorph software. To discriminate the fluorescent spots representing autophagolysosomes from background and to ensure that all staining analyzed was only the true positive labelling, a defined threshold value was set for 488 nm excited fluorescence pixel intensity. The area of cell occupied by autophagolysosomes was expressed as fraction of total cell area. ^10^

## Supplemental Figures

**Figure S.1.**
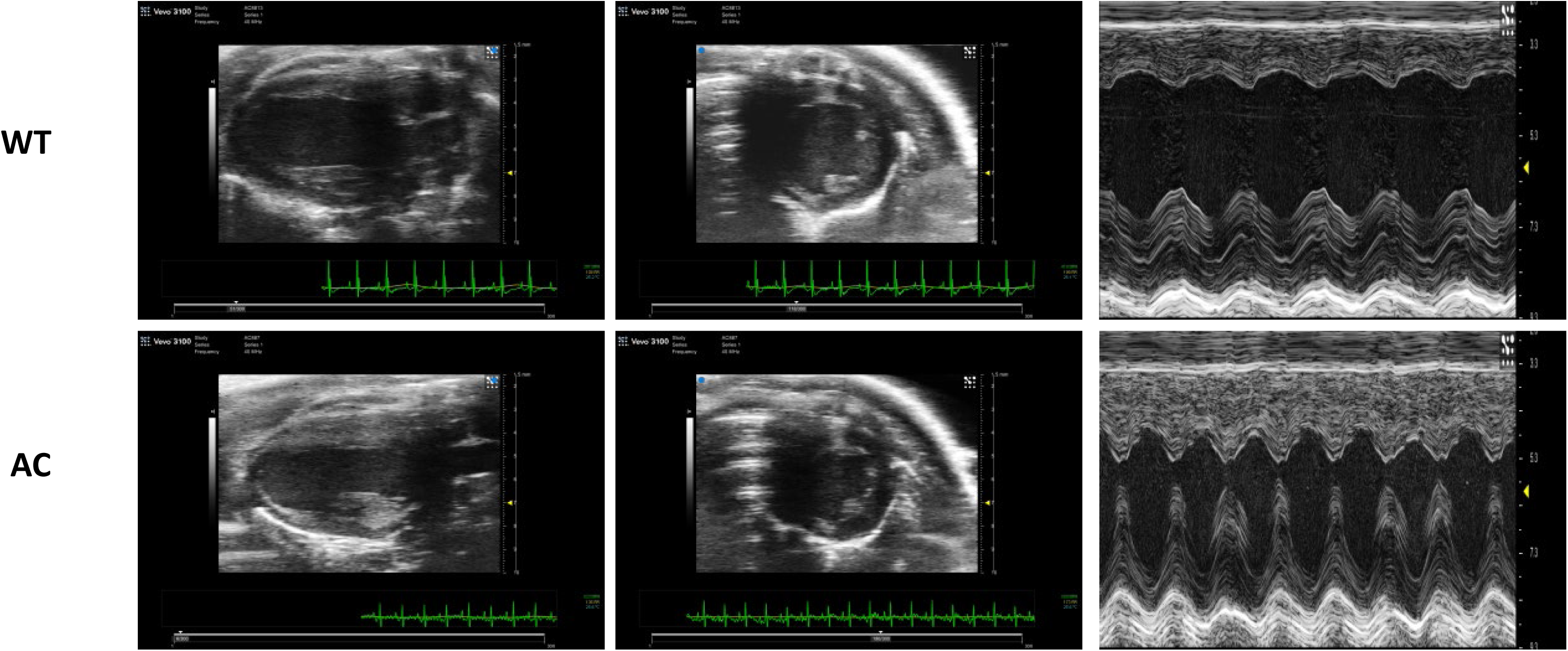
Representative images of Echocardiograms of TG^AC8^ and WT LV.

**Figure S.2.**
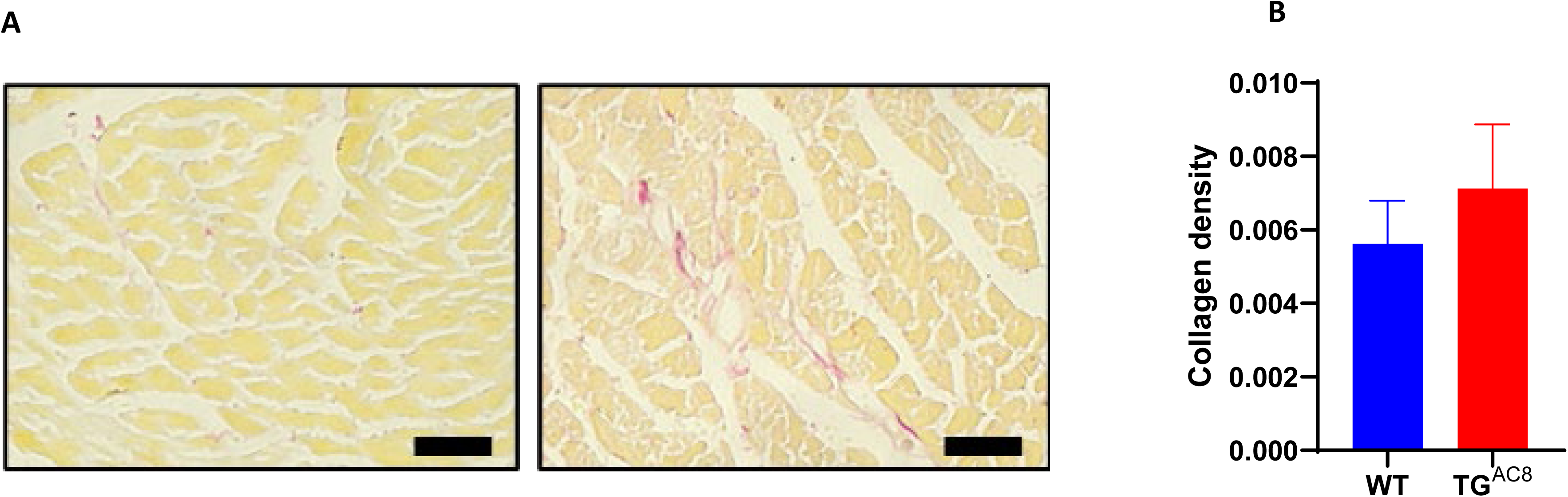
Representative LV sections, labeled with picrosirius red (A) and (B) average collagen density (picrosirius red labeling) in TG^AC8^ vs WT LV.

**Figure S.3.**
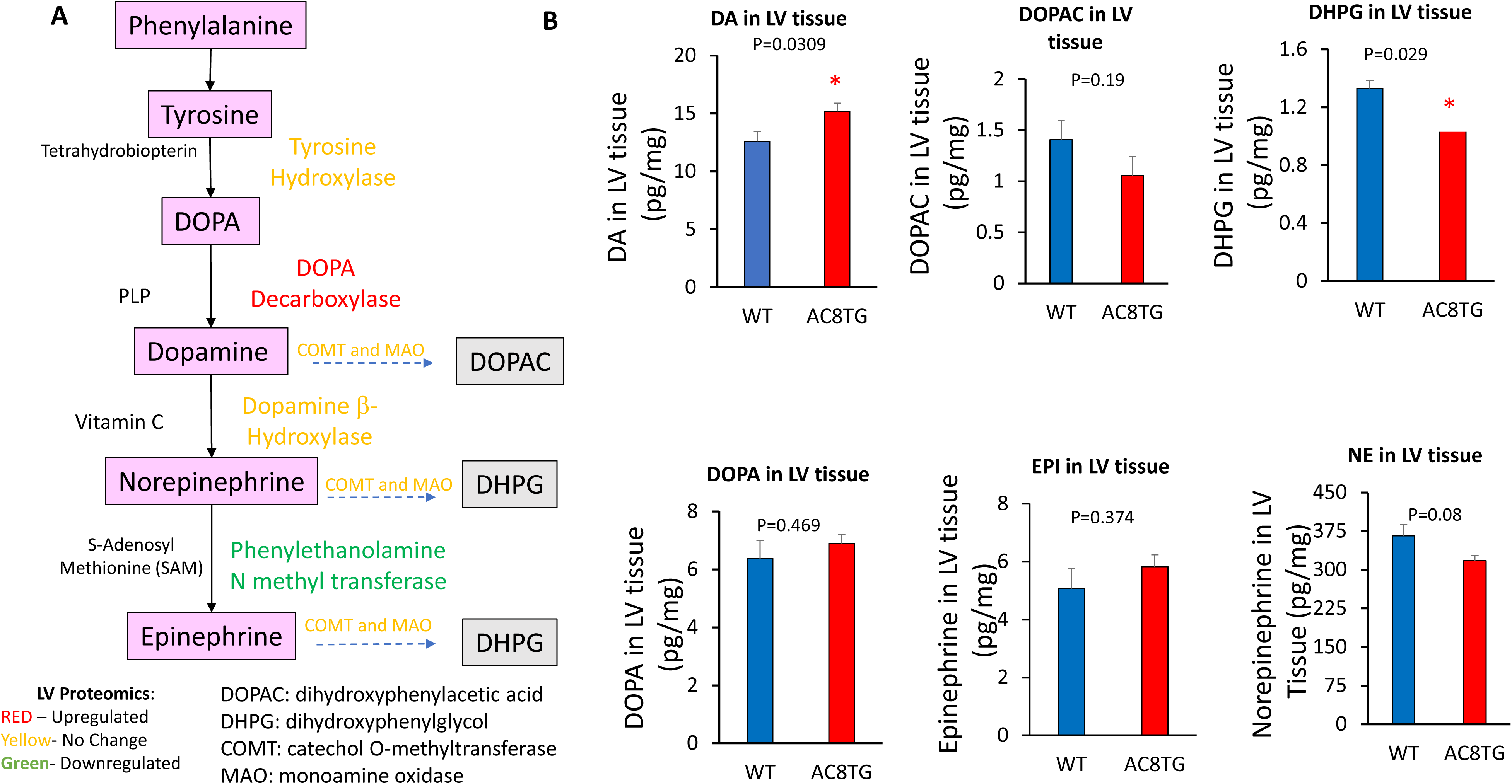
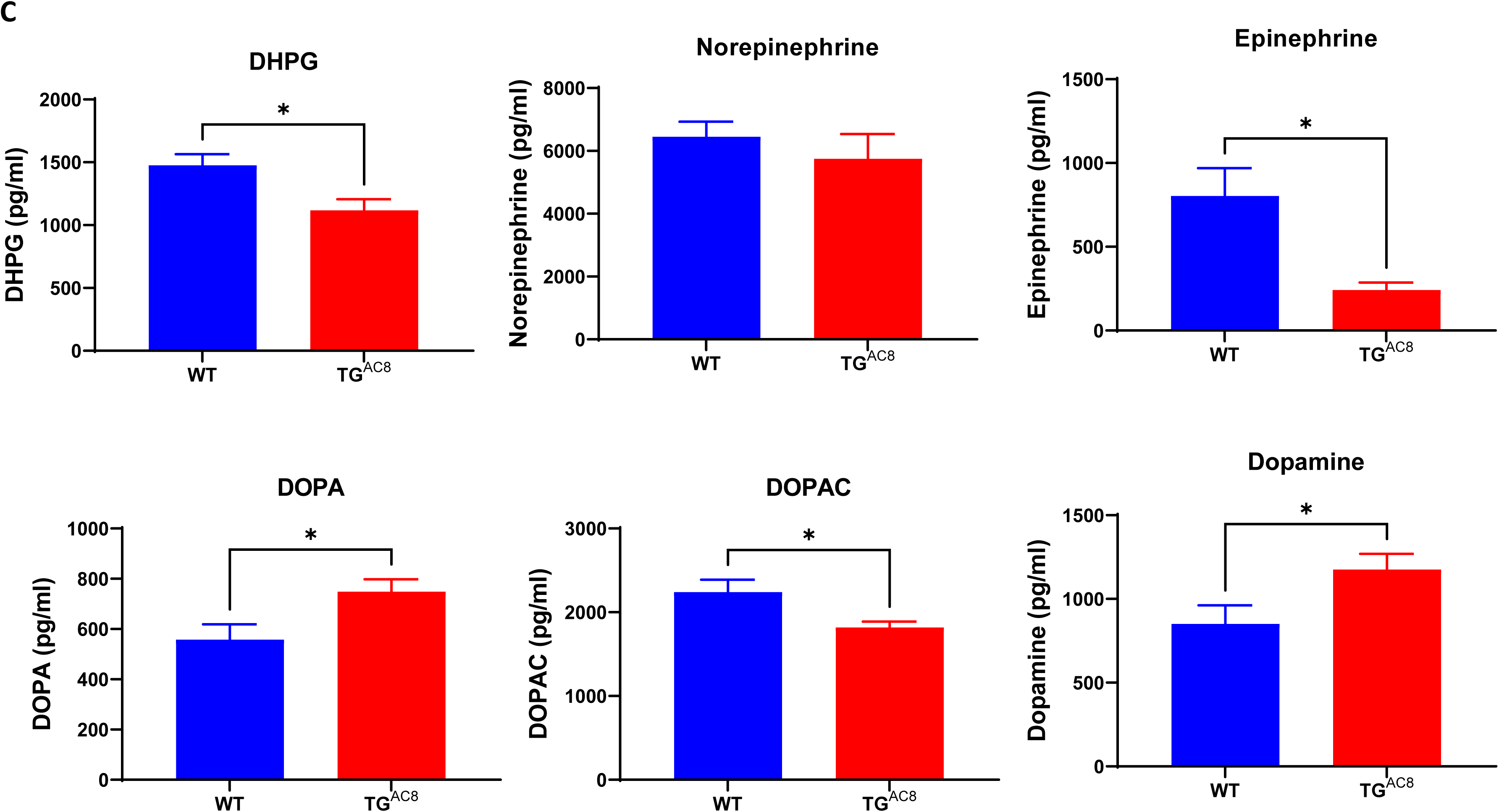
(A) Pathway for catecholamine synthesis and breakdown; (B) Myocardial catecholamine levels. (C) plasma catecholamine levels.

**Figure S.4.**
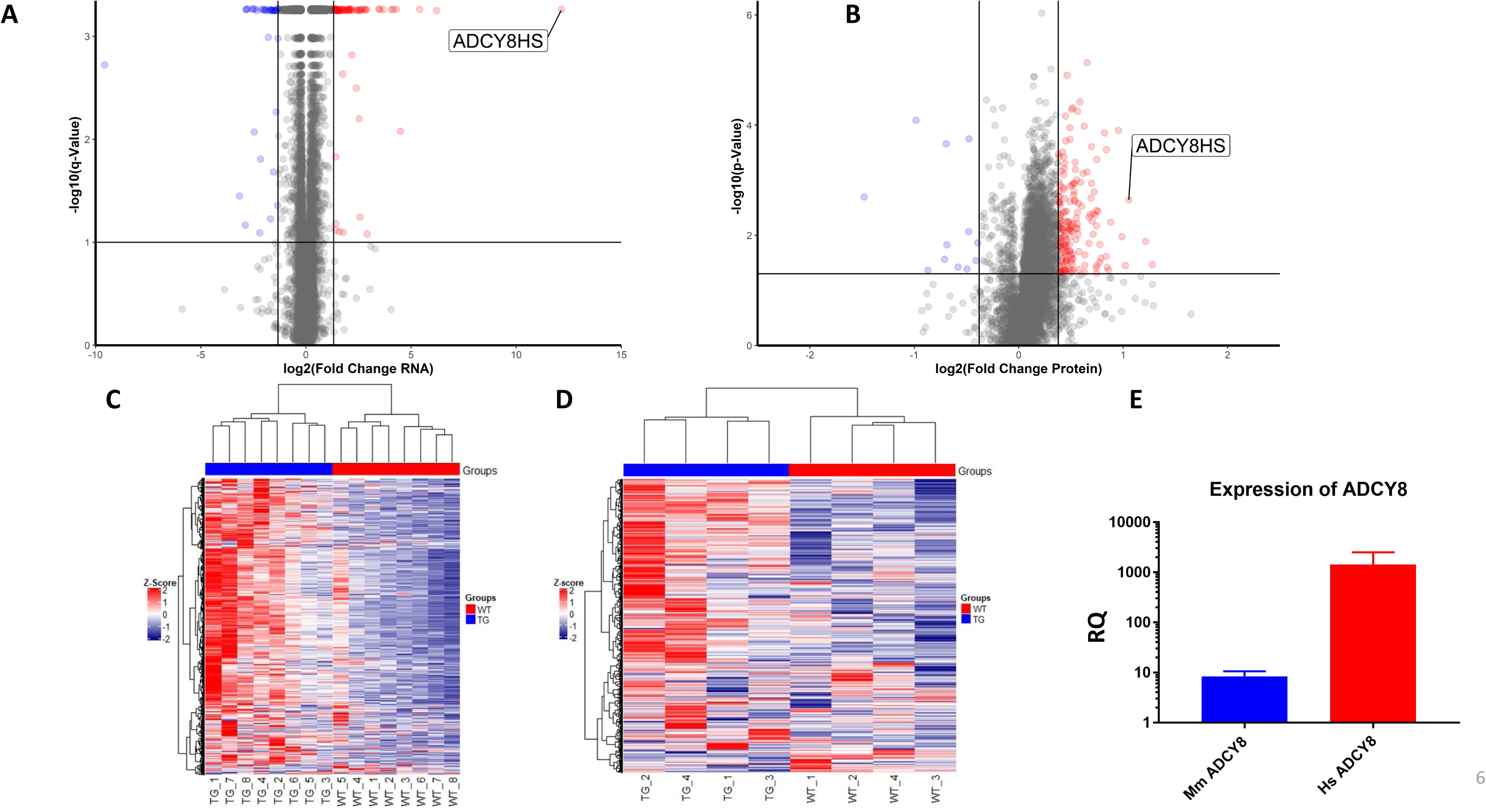
Volcano plots (A and B) and heat maps of transcripts (C) and proteins (D); (E) expression of mouse and human types of *Adcy8* in LV

**Figure S.5.**
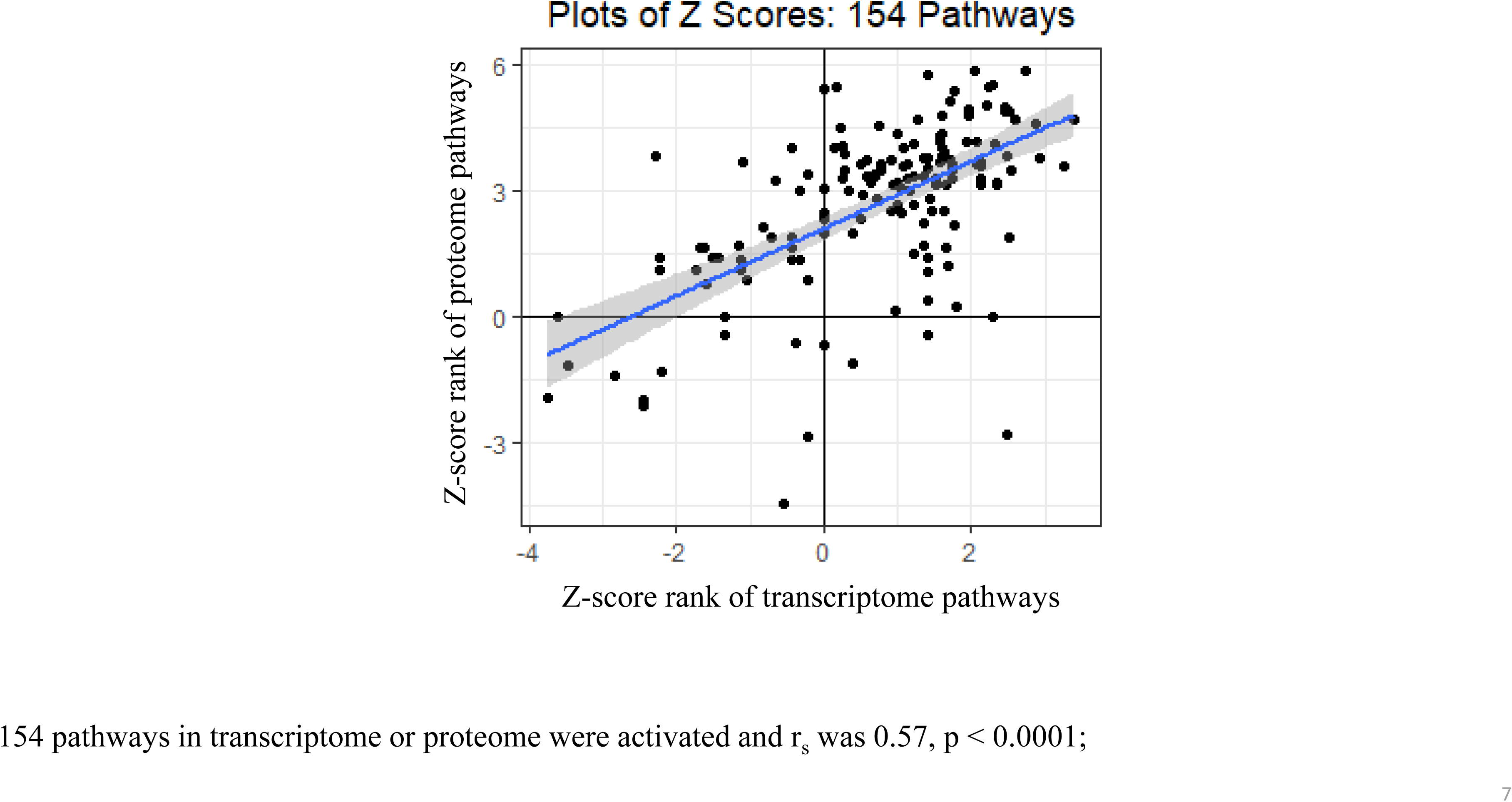
**Plot of the ranks of Z-scores of IPA transcriptome and proteome canonical pathways that significantly differed by genotype.**

**Figure S.6.**
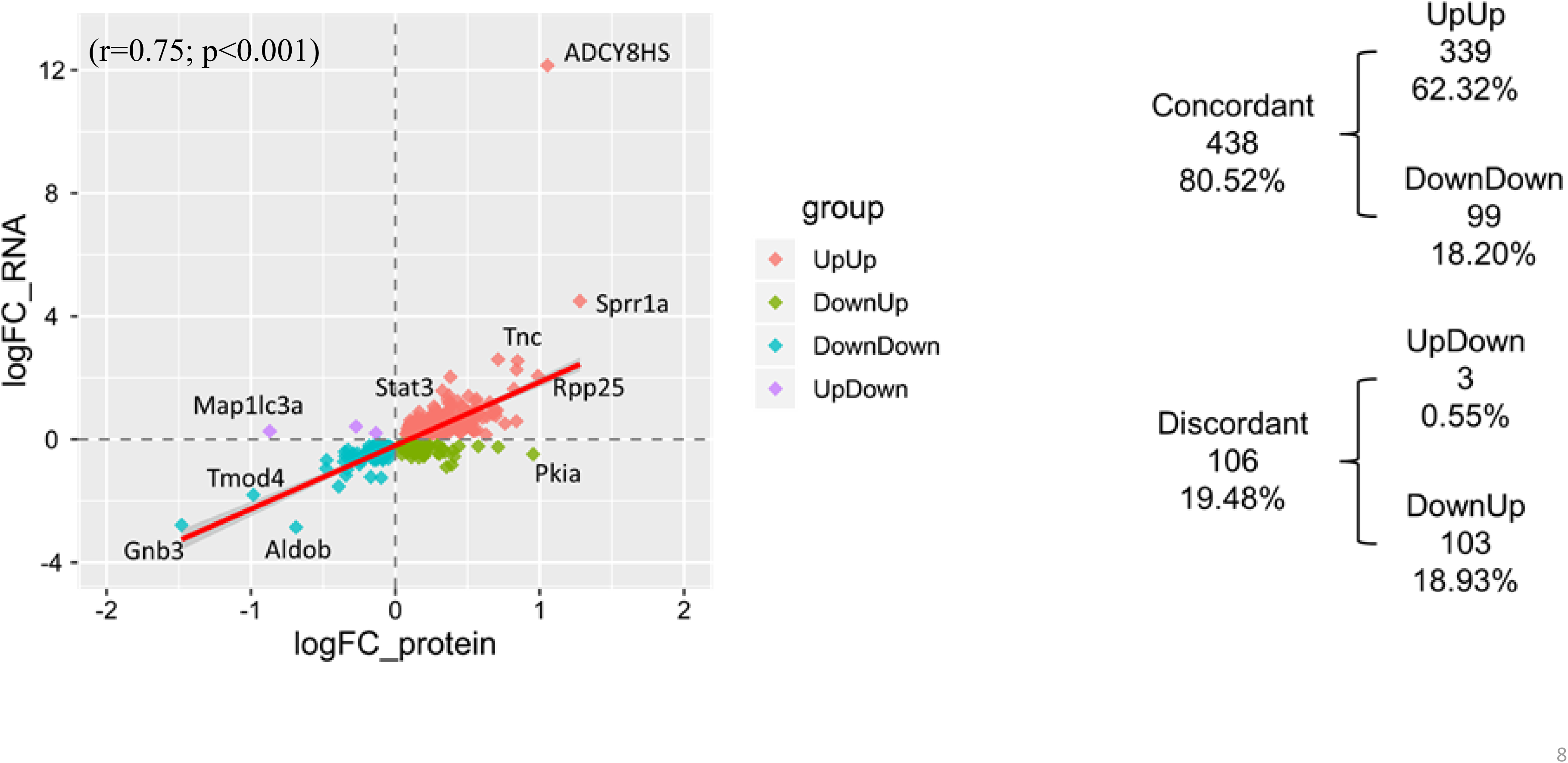
Correlation plot of 544 identified molecules of which *both* transcripts and proteins differed by genotype.

**Figure S.7.**
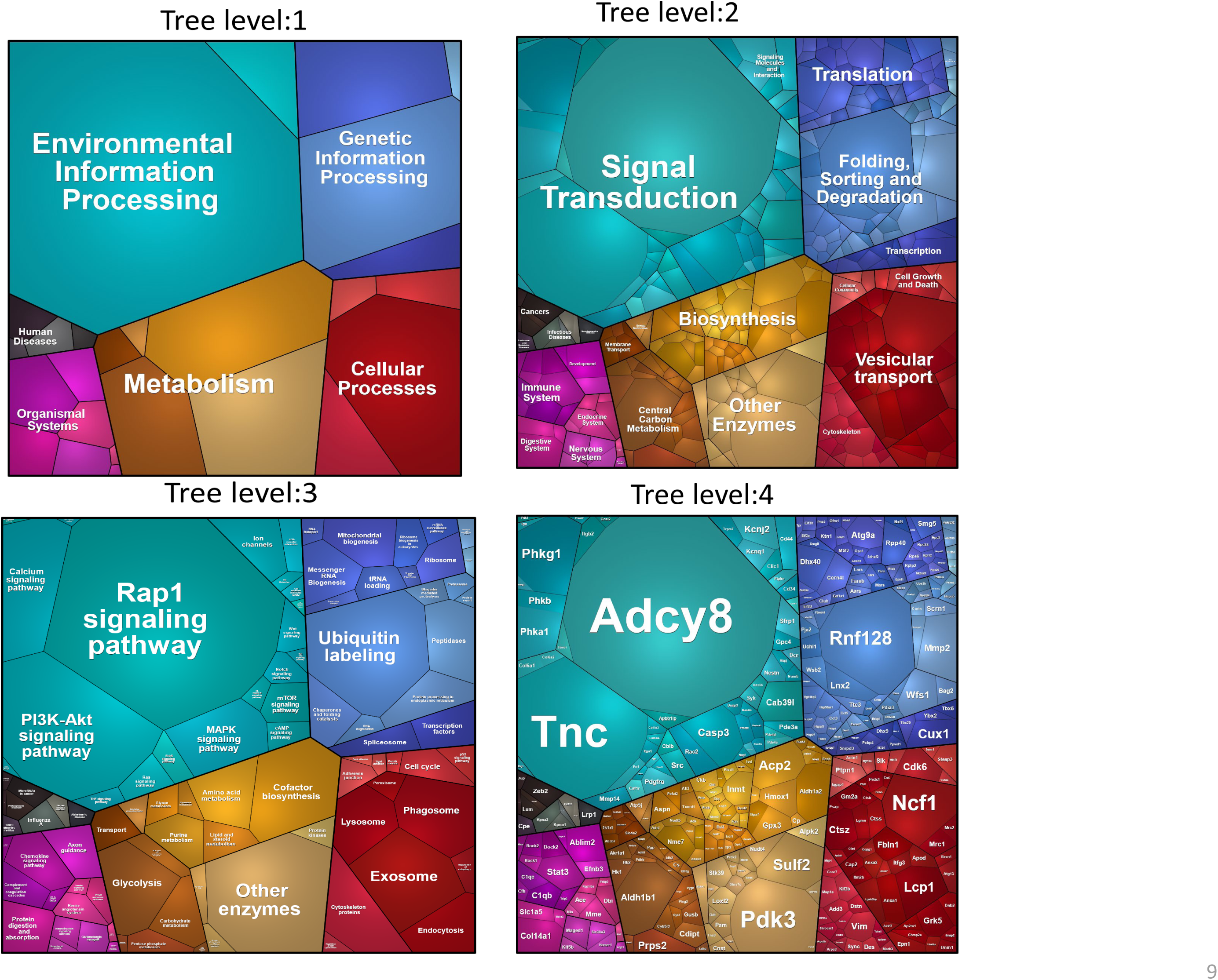
PROTEOMAP. Areas of polygons within each tier reflect genotypic differences in protein abundances, weighted by protein size.

**Fig S.8.**
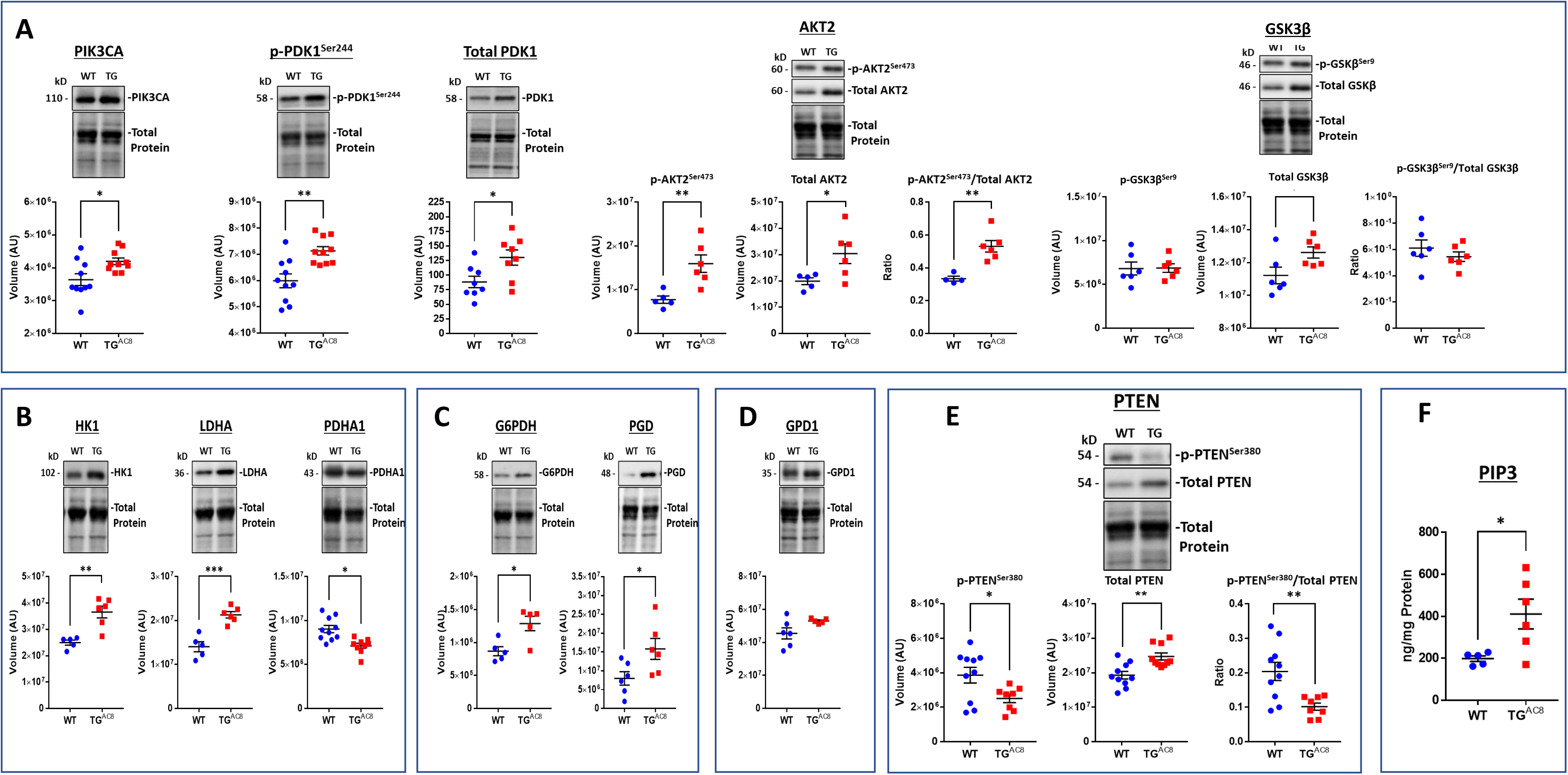
(A-E) WB and ELISA (F) analyses and of proteins that mediate PIP3 Kinase Signaling and Metabolism

**Fig S.9.**
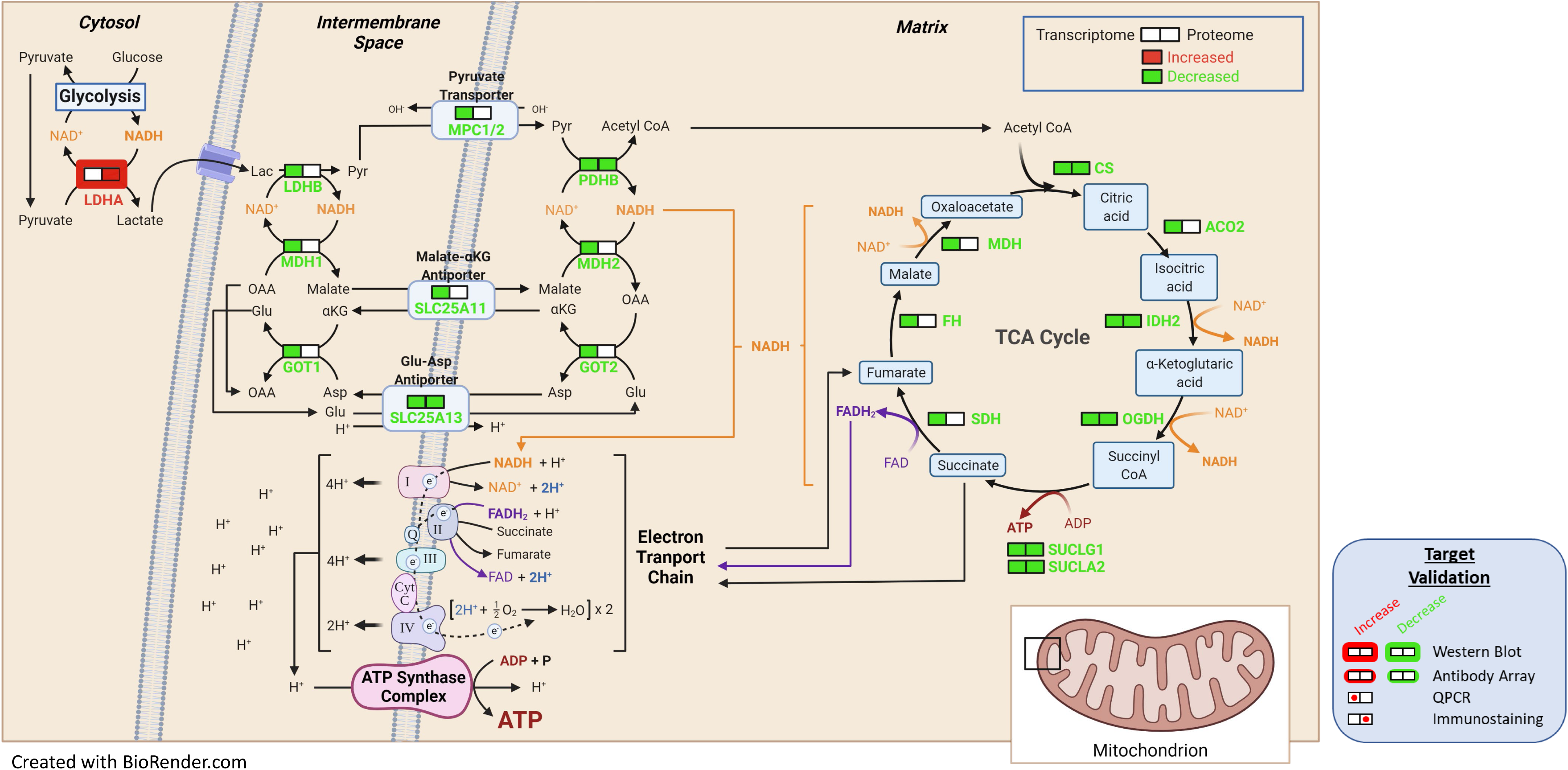
Detailed schematic malate-aspartate shuttle based on signals derived from bioinformatic analyses of the transcriptome and proteome and on selected WBs.

**Figure S.10.**
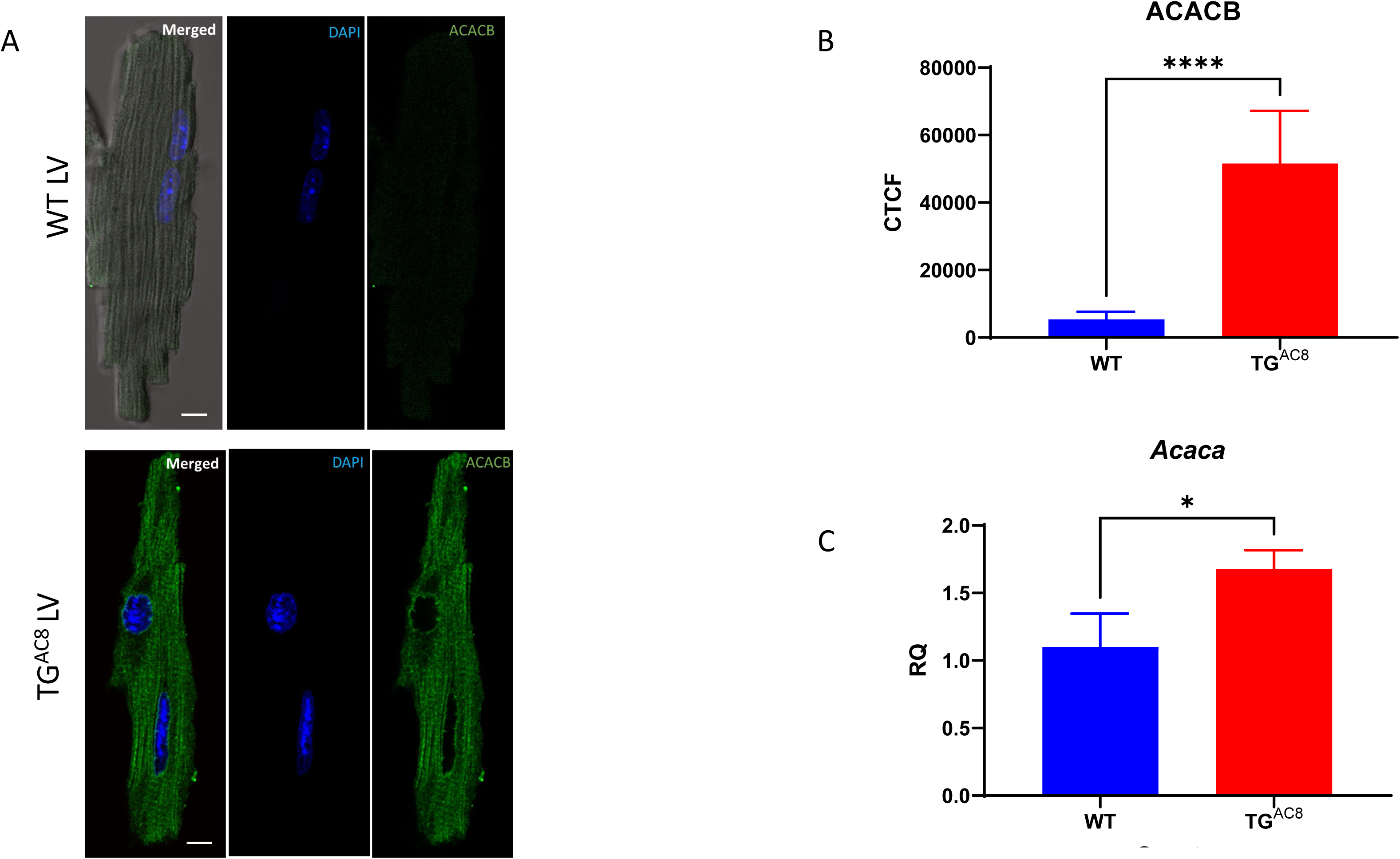
Representative examples of (A) ACACB immunolabeling of TG^AC8^ and WT LV myocytes; (B) average ACACB fluorescence in LV cardiomyocytes (n=25 for each group); (C) Relative Quantification of *Acaca* mRNA expression in LV tissue (n=4 for each group).

**Figure S.11.**
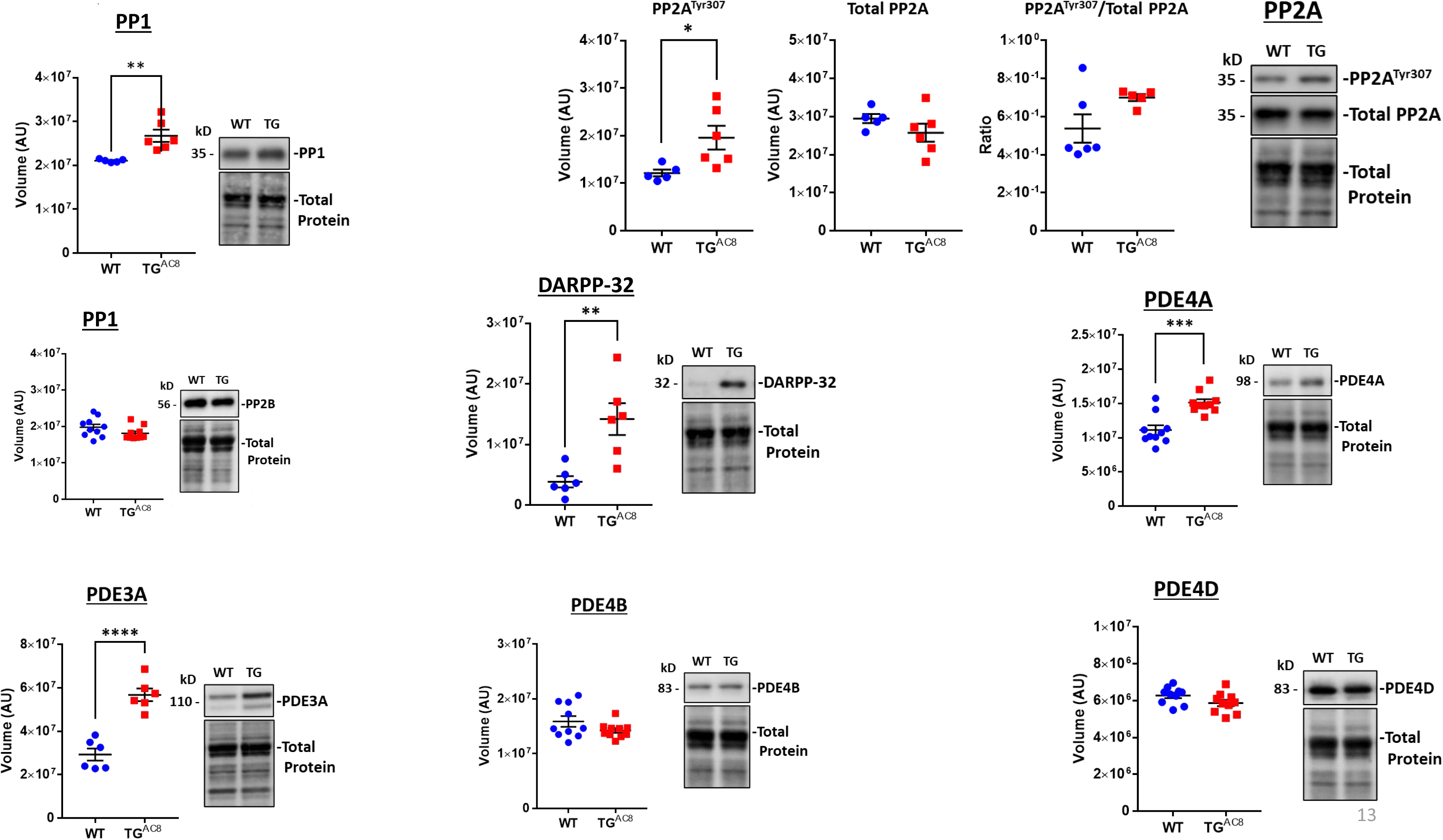
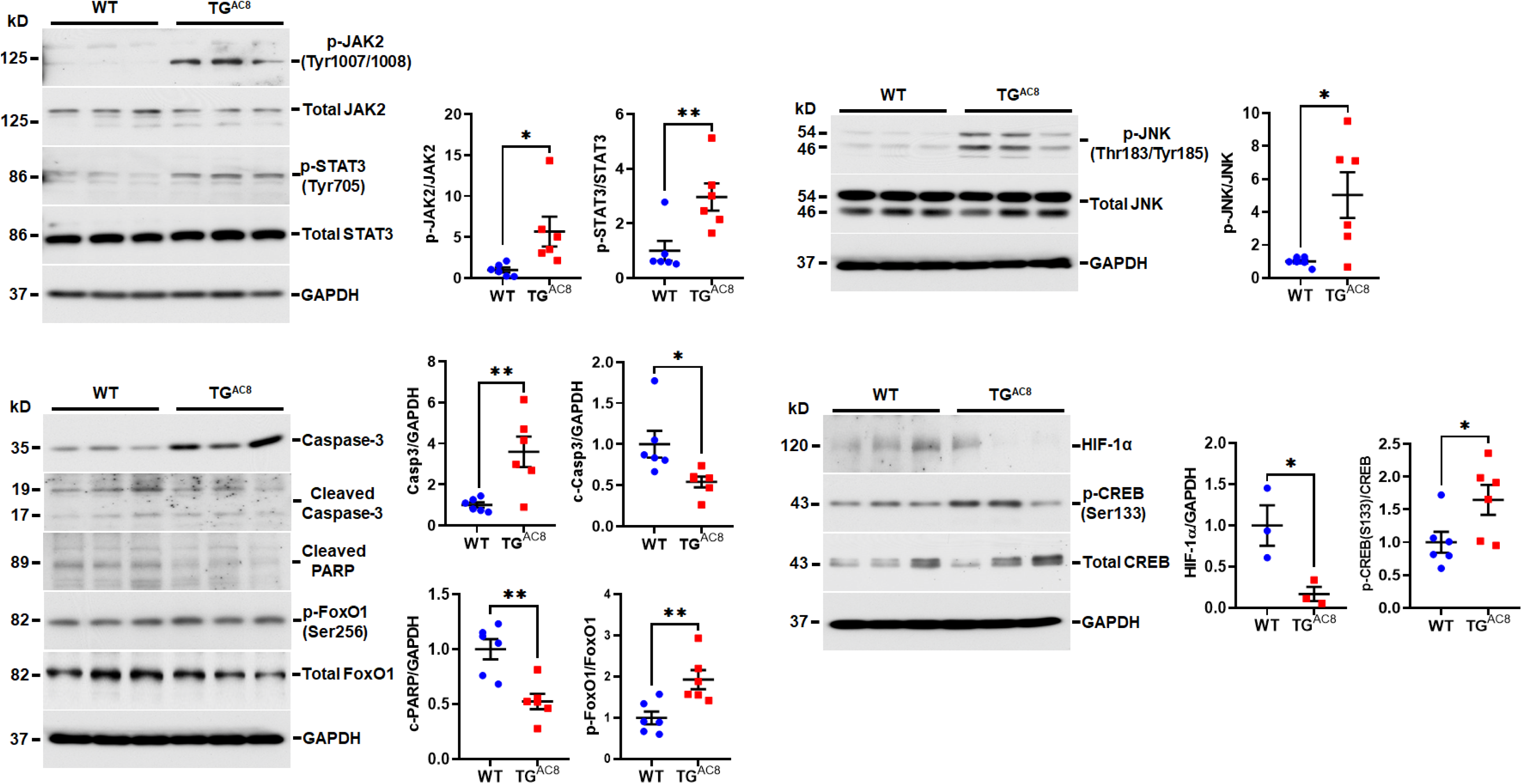
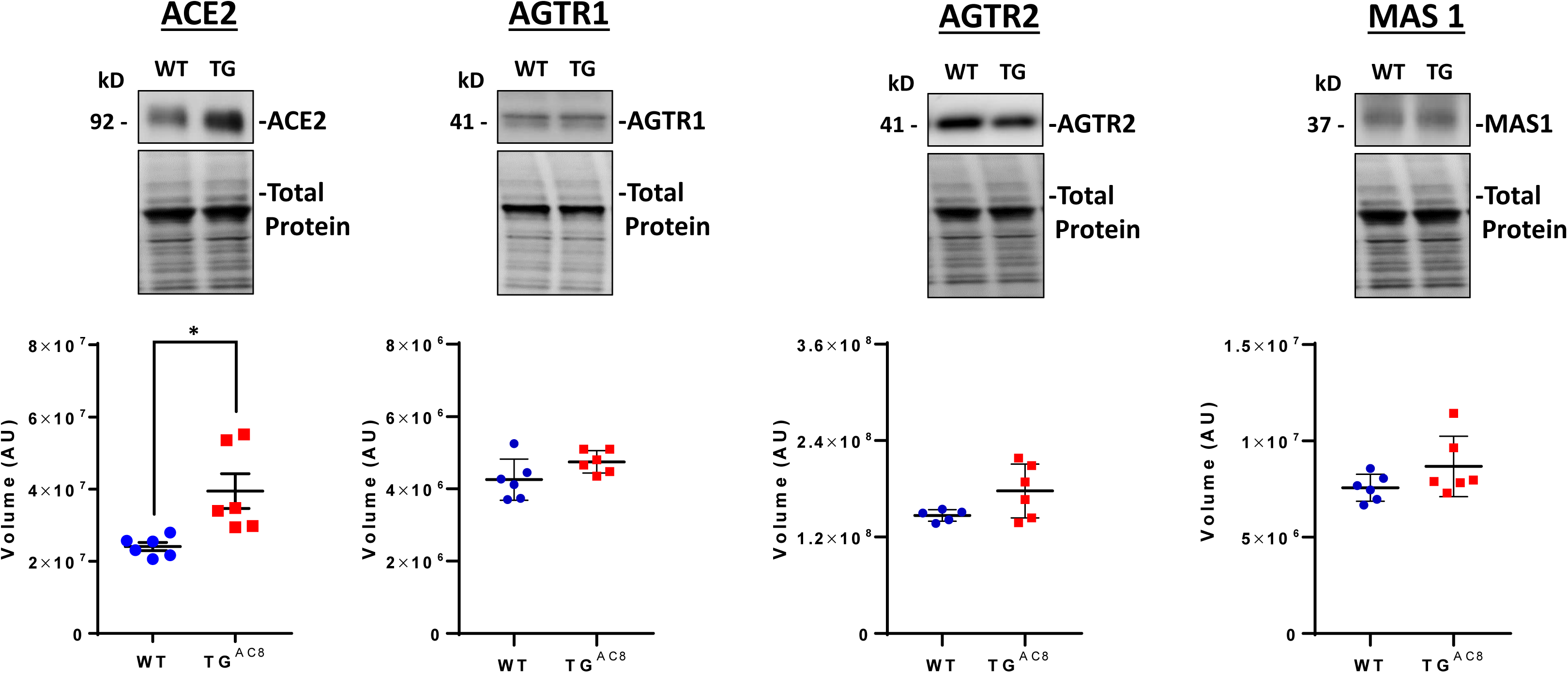
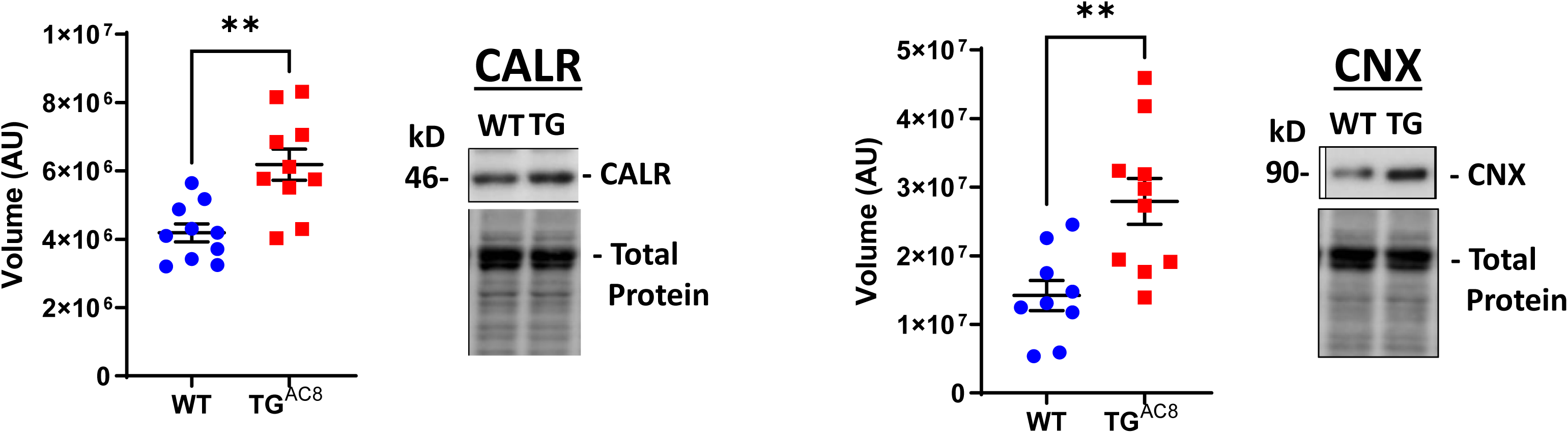
(A) Negative feedback adaptations on AC/PKA signaling (B) WB analysis of selected proteins involved in Jak/Stat/Jnk/Caspase signaling (C) WB analyses of selected proteins involved in angiotensin receptor signaling (D) WB analysis of Calnexin and Calreticulin, proteins involved in ER protein processing.

**Figure S.12.**
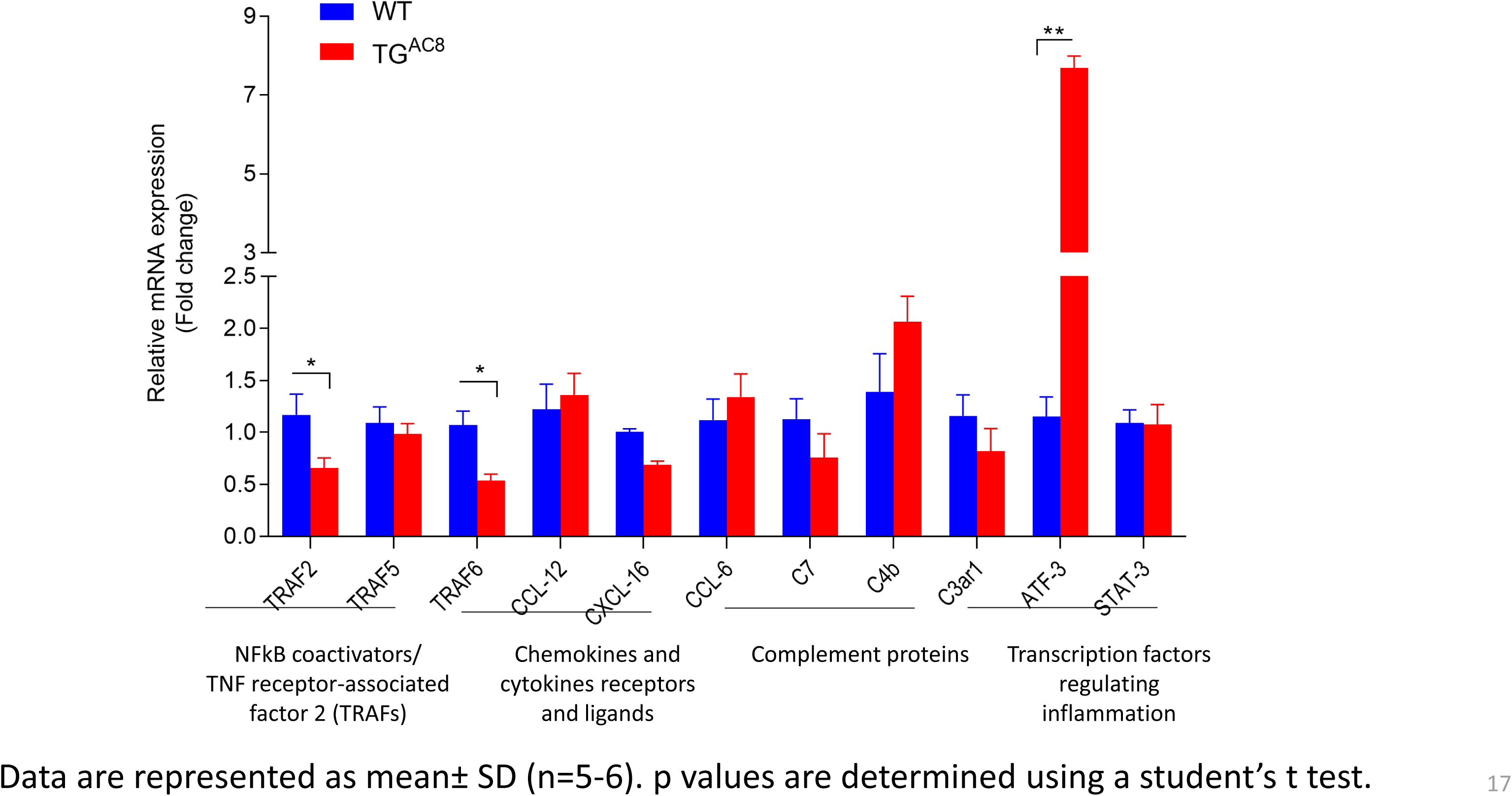
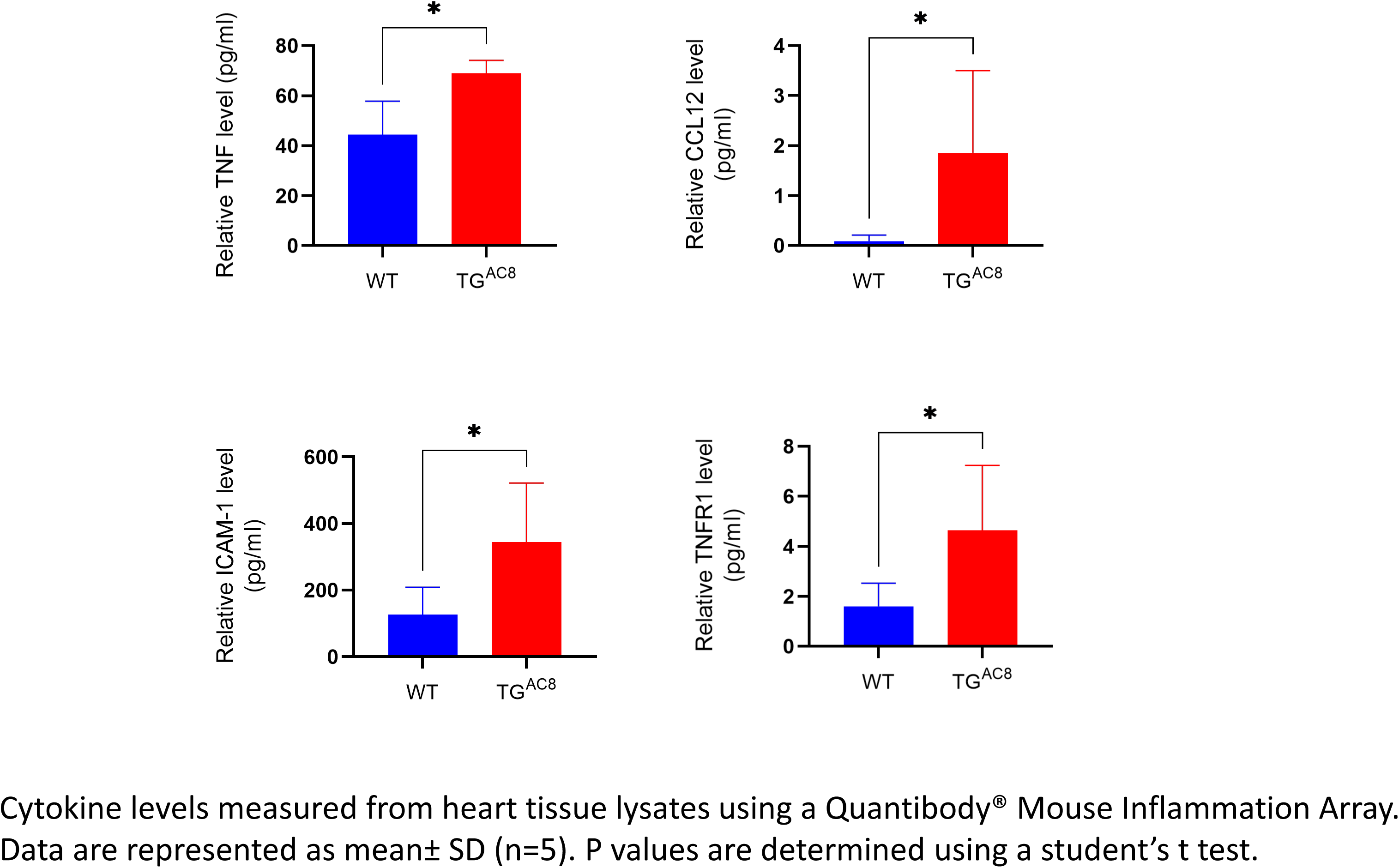
(A) . QRT-PCR analysis of genes regulating cytokines level in the heart. (B) Cytokines levels measured from heart tissue lysates.

**Figure S.13.**
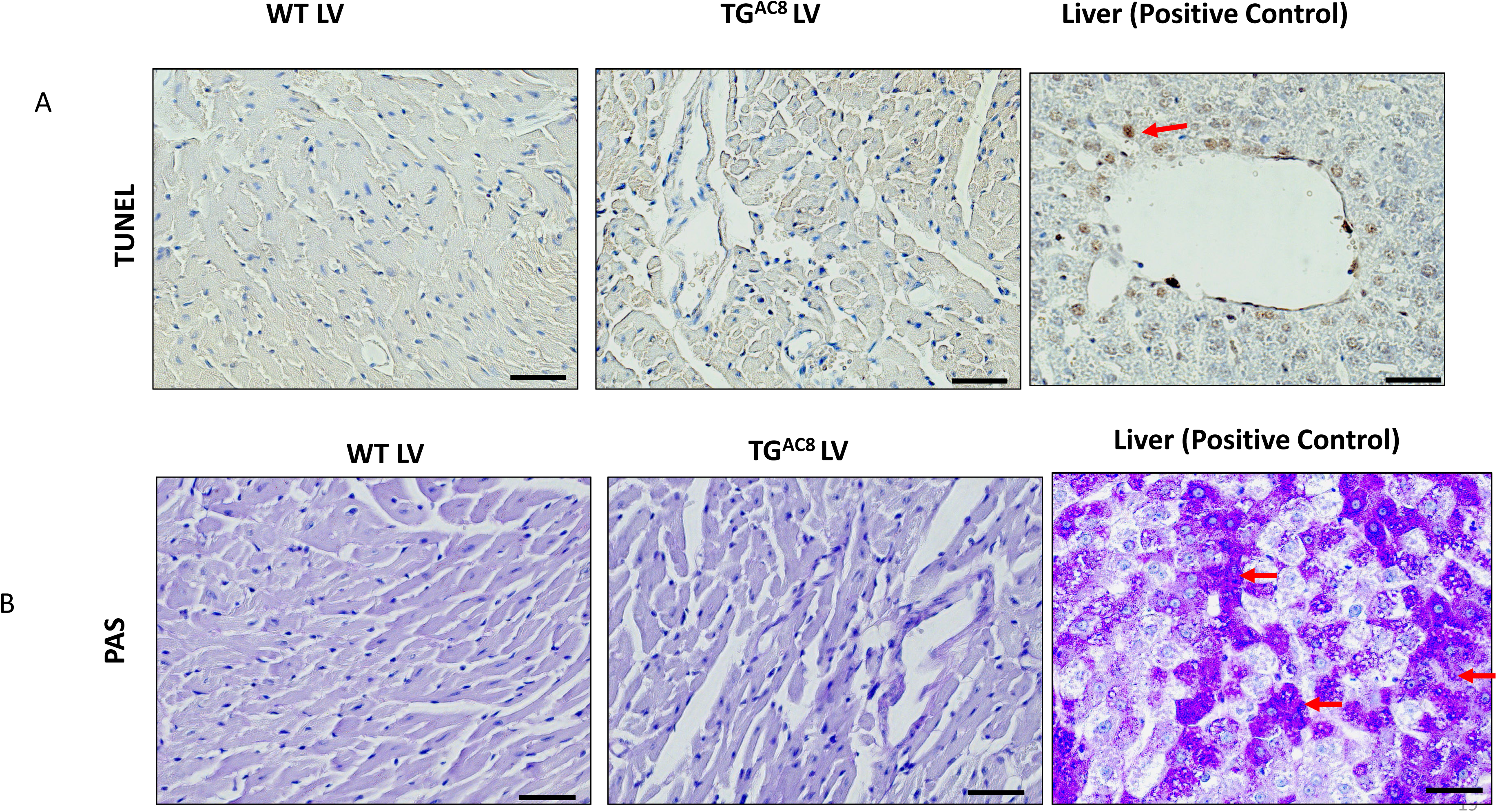
LV Tissue Staining for (A) apoptosis and (B) Glycogen

**Figure S.14.**
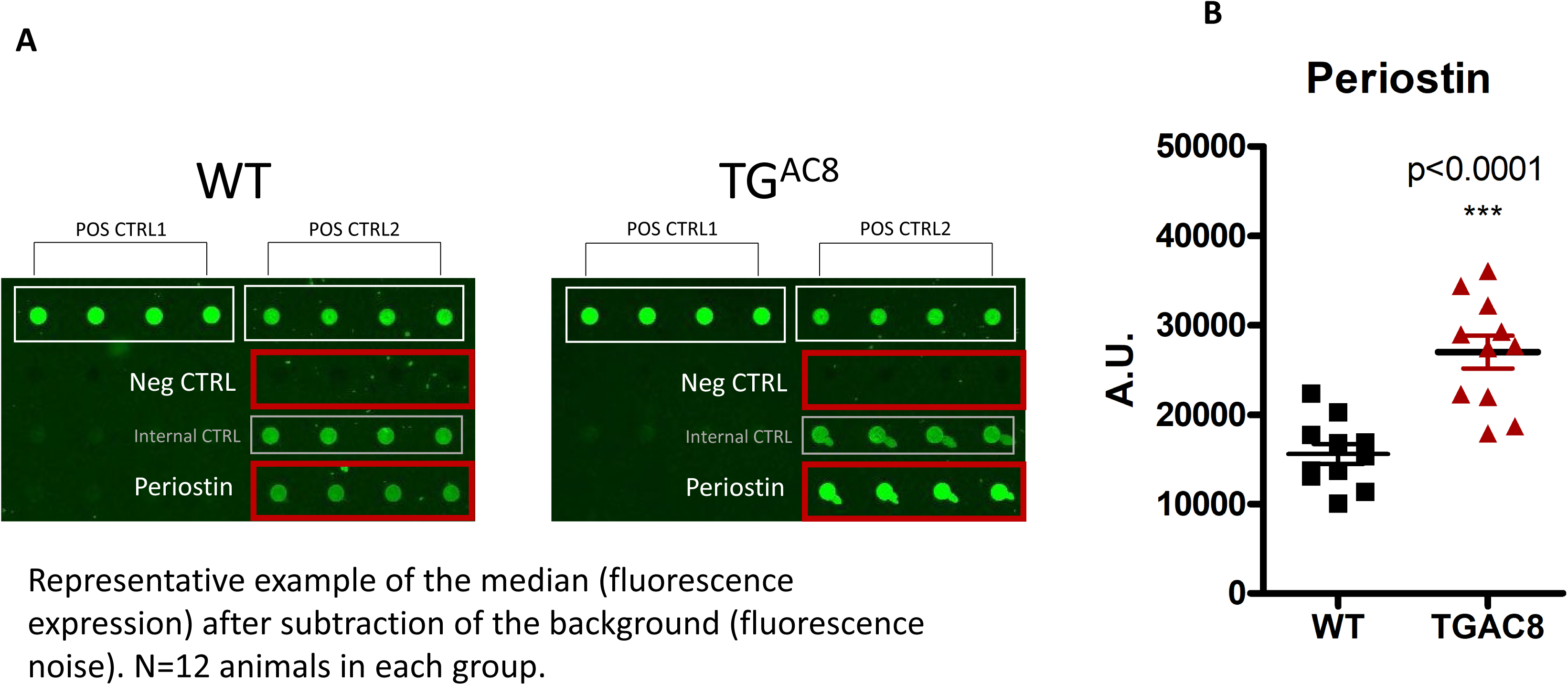
**Periostin levels detected in TGAC8 vs WT LV (Growth Factor Quantibody Array)**

**Figure S.15.**
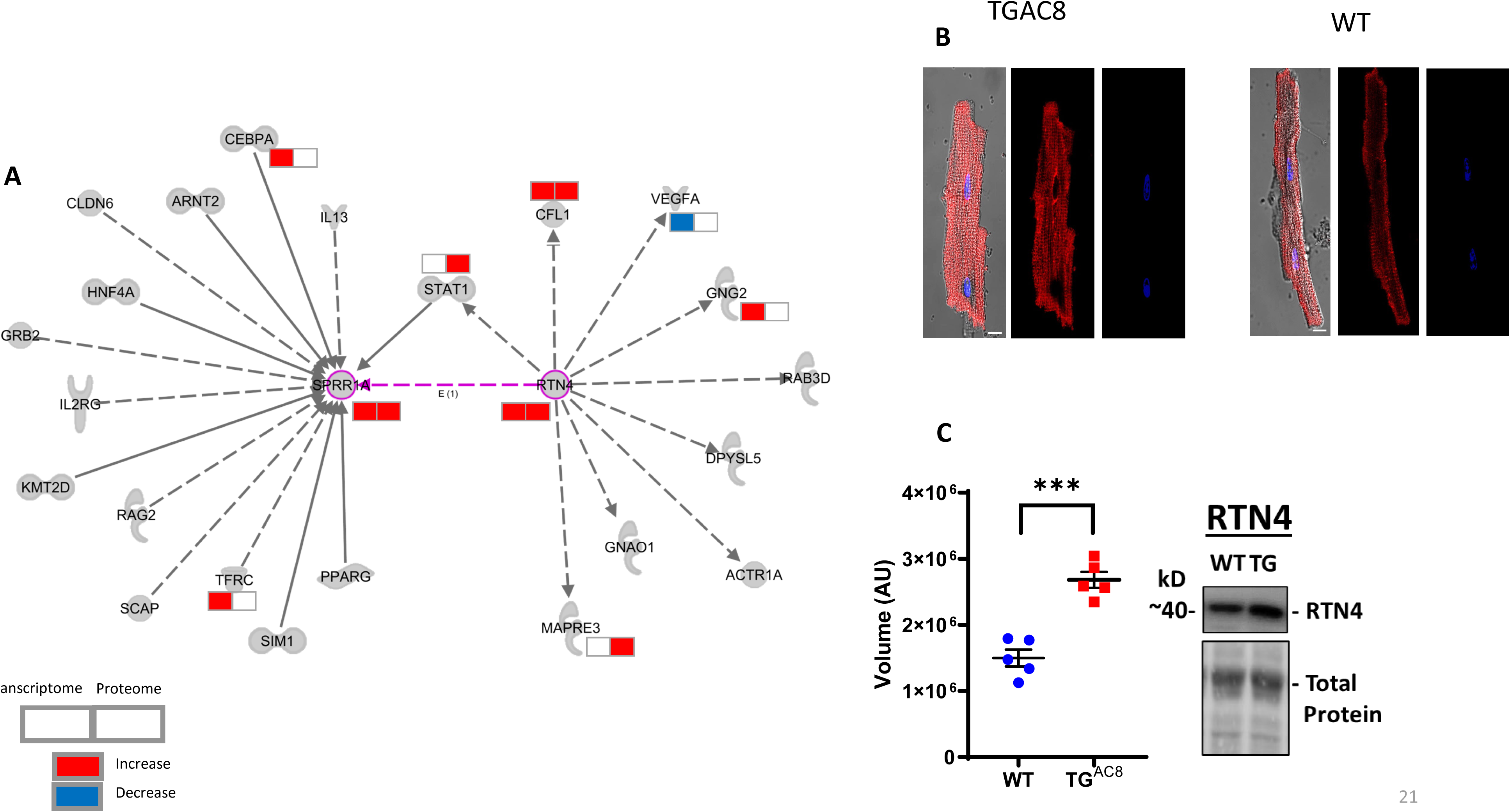
(A) Sprr1 signaling network. (B) Immunolabeling of Sprr1a in LV myocytes isolated from TG^AC8^ and WT. (C) WB analysis of Rtn4 expression

**Figure S.16.**
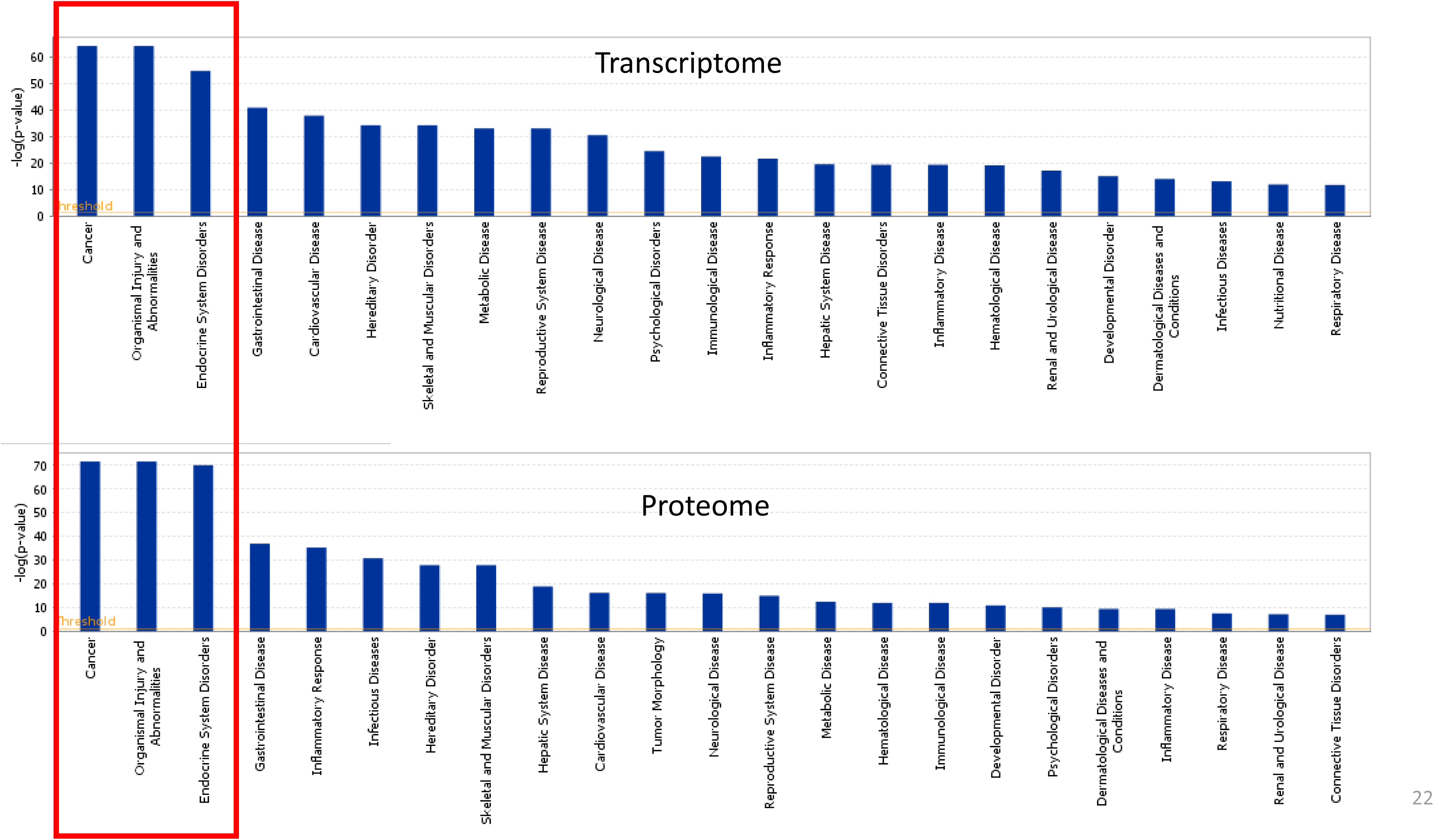
IPA representation of top. Disease-related functions within the LV transcriptome and proteome of TGAC8 and WT.

